# Formalizing host competence

**DOI:** 10.64898/2026.11.12.766225

**Authors:** Nathaniel L. Gibson, John M. Drake, Tara E. Stewart Merrill, Kailey M. McCain, Caroline Merriman, Kirsten M. Yeomans, Lynn B. Martin

## Abstract

Host competence – the capacity to acquire, sustain and transmit parasites or pathogens – is typically quantified for species, although it is also an individual property. We derive an individual-level competence index from infection biology and link it to reproduction numbers. Our index separates infectiousness and susceptibility (individual properties) from contact and abundance (environmental properties); combined, they yield host potential and capacity, analogues of vector potential and vectorial capacity. We simulated populations in four systems (Lyme disease, West Nile virus, chytridiomycosis, avian influenza), using experimental data where available and plausible values otherwise. Individual competence varied widely within and overlapped among species; species means approximated an established index for West Nile virus, to which the model was calibrated. Under the assigned variances, intraspecific variation was greatest in low-competence species, whose loads sit near the transmission threshold; there, load–contact covariation roughly halved or doubled mean potential. Where contact was measured (Lyme tick burdens), nearly all individuals of high-competence species replaced themselves across plausible tick survival, whereas replacement by most individuals of low-competence species depended on tick survival. In West Nile virus, the most infectious 20% of mourning doves supplied about half of transmission, so species averages could understate the role of some individuals.

## 1. Introduction

**Host competence** is a key concept in disease ecology. Defined as the capacity of a host to acquire, sustain, and transmit a parasite or pathogen to another host, vector or the environment, host competence ultimately determines whether a host will dampen, magnify or leave unchanged local risk of infection (Cortez & Duffy 2021; Gervasi et al. 2015). Indeed, host competence governs the dynamics of epidemics because it determines which hosts sustain transmission (DeMarchi et al. 2026; Pérez-Umphrey et al. 2026). It therefore also influences community-level relationships between, for example, diversity and disease (Keesing et al. 2010; LoGiudice et al. 2003; Stewart Merrill et al. 2022). As with many other complex traits, variation in host competence is not driven by simple processes operating at one level of organization (Downs et al. 2019; Huang et al. 2023). Rather, variation in host competence arises from an interplay of physiological, immunological, and behavioral forces, some of which are genetic, some of which are environmental, and some of which entail GxE interactions (Gervasi et al. 2015; Martin et al. 2021). This variation aggregates to the population and community levels, bridging heterogeneity in the internal biology of single hosts to the collective fate of hosts, vectors, and competitive parasites.

Despite its centrality, host competence remains operationally and quantitatively vague (Gervasi et al. 2015; Martin et al. 2016; Mommer et al. 2025; Stewart Merrill & Johnson 2020). Several similar terms have been proposed including reservoir competence (Brunner et al. 2008), realized reservoir competence (LoGiudice et al. 2003), and host quality (Mommer et al. 2025). Epidemiological terms such as the individual reproduction number (Lloyd-Smith et al. 2005) and the next-generation matrix (Diekmann et al. 2010 sensu Roberts & Heesterbeek 2013) have been offered, too. These latter terms iterate on a pathogen’s basic reproduction number, *R_0_*, to describe the role of individual hosts in transmission. The value of constructs such as competence is that they can bridge research on host variation with actionable outcomes (Kilpatrick et al. 2005). The current shortcoming of most efforts, though, is that most have been parameterized for types (i.e., species or life stages), usually by averaging measurements made on relatively few individuals, and rarely decomposed into organismal traits that vary among individuals (Fenton et al. 2015; Hofmeester et al. 2016; Levi et al. 2016; Streicker et al. 2013).

Ideally, an index of host competence would be formally described, starting from first principles and based on knowledge about variation in the phenotypic traits of individual host organisms (Huang et al. 2023; Stewart Merrill & Johnson 2020). It is through this route that variation in competence becomes common currency across scales of biological organization (Downs et al. 2019) from host-pathogen coevolution to seasonal cycles of risk to spillovers of novel infectious agents. Indeed, such an index could be useful to probe more explicitly the role of individual host heterogeneity in the ecological processes of transmission (Hawley & Altizer 2011; Kilpatrick et al. 2006; Leitch et al. 2026; VanderWaal & Ezenwa 2016), especially when community diversity is low (Violle et al. 2012) or the environment itself causes stress (Gervasi et al. 2015).

Vector biology offers some useful guidance to formalizing host competence. Specifically, vector biology has long been focused on two related concepts: vector competence and vectorial capacity. Vector competence is analogous to host competence, specifically quantifying the probability that an individual vector acquires, supports, and transmits a pathogen once exposed. Vectorial capacity, by contrast, is a population-level composite of vector competence, host biting rates, survival over the period of transmissibility, and the density of the focal vector population (Smith et al. 2012). Because vector competence and vectorial capacity share a mathematical framework, individual and higher-level processes can be understood together, providing practical insight unavailable otherwise (Garrett-Jones 1964; Hardy et al. 1983; Smith et al. 2012). Consider climate change, which is expected to alter arboviral disease risk in the future. Because vector competence and vectorial capacity are directly related, models can incorporate individual responses (such as thermal tolerance), population demographics, and ecological processes, to generate predictions (Mordecai et al. 2019). Further, when combined with host use measures (e.g., the identity of vector blood meals), vector potential can provide insight into the role of vector populations on local infection risk (Bauer et al. 2026).

Here we derive a host competence index, from first principles, and use it to ask three questions (full derivation in Appendix S1). First, can a single expression, built from the biology of infection, identify and capture the critical processes that shape competence? Second, when the index is applied to individuals rather than species, where does variation in competence arise, and which host traits drive it? Third, when does variation in individual host competence matter for transmission in a community? We address these questions by deriving a general equation for the individual trait, defining its population- and community-level extensions, and examining it in representative systems. In doing so, we demonstrate the usefulness of our theory as both a tool for empirical applications and its heuristic capacity for identifying new directions of research.

## 2. Deriving host competence, potential and capacity

Host competence is a composite of physiological, immunological, and behavioral factors that together determine whether a single host generates meaningful infection risk for other hosts (Becker et al. 2020; Downs et al. 2019; Martin et al. 2016). Our goal is a quantity that describes what an individual host contributes to pathogen transmission, built up from the biology of infection and connected explicitly to reproduction numbers (i.e., number of new infections caused by one host). We construct this index in four steps that follow the main events in a transmission cycle: i) whether an exposed host becomes infected, ii) what happens within a host once it is infected, iii) how often an infected host contacts susceptible recipients, and iv) how many such hosts there are in a population. The first two are properties of the individual host and can be measured in isolation. The last two are properties of the ecological setting. Keeping the two kinds of term separate allows competence to be scaled up to populations and communities, just as vector competence is scaled up to vectorial capacity (Garrett-Jones 1964; Smith et al. 2012).

### Susceptibility and Infectiousness

The first aspect of competence involves encountering another infected host or vector, but as this stage entails variation in both parties, we separate the role of a focal host (and pathogen) from other factors. Subsequently, we identify this first aspect of host competence as **susceptibility**, the dose-dependent infection probability of a focal host given encounter or ε(D), the probability that a host becomes infected when exposed to a dose D.

**Infectiousness** subsequently describes how an infection proceeds within a host: how much pathogen it carries, for how long, and how often that load can reach a susceptible host. Consider a host individual infected at time zero and let τ denote infection age. The within-host pathogen load of this individual follows a trajectory L(τ). Load affects transmission through φ(L), the probability that a transmission-relevant contact made at load L infects a reference recipient (for example, another host or a vector); φ is assumed to be saturating in load (Fraser et al. 2007; Ke et al. 2021; Marc et al. 2021). In our scenarios, the host remains a source only while it is alive and has not cleared infection, which we write as Σ(τ) = exp(−∫ ^τ^ [µ(u) + γ(u)] du), where µ is the mortality hazard and γ the clearance hazard. Finally, infection often changes behavior: sickness behavior can suppress activity and contact, and anti-vector behavior such as grooming can stop vectors from completing a blood meal (Adelman & Martin 2009; Keesing et al. 2009). We capture these possible effects with b(τ), the rate at which an infected individual realizes transmission-relevant contacts relative to that individual’s rate before infection (b = 1 when infection does not alter behavior). Integrating over the infection gives **infectiousness**:

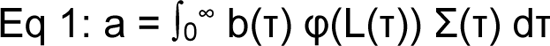

The quantity *a* has units of time. It is the number of fully infectious, fully active days that an infected host provides, which we call effective infectious days. It is conditional on infection, matching the experimental concept of competence in other work in which animals were deliberately infected and their infectiousness to vectors or conspecifics measured (Brunner et al. 2008; Komar et al. 2003). Because b and Σ can depend on load, *a* can represent sickness behavior, pathogen tolerance, and/or virulence within a single formula. We keep φ and b definitionally distinct: φ describes what happens given a contact and b describes how often contacts happen, so that the effect of pathogen load is never counted twice.

Multiplying the two individual-level terms gives host competence, the expected number of effective infectious days generated per exposure:

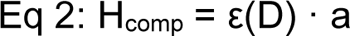

Competence is therefore often described as the product of two host properties: susceptibility and infectiousness. Transmissibility entails how an infected host behaves, how often it encounters susceptible recipients, and how vectors respond to it. Accordingly, transmissibility includes both individual host (e.g., behavior that changes at the time or after an infection: sickness behavior, grooming, anti-vector defense) and environmental/ecological components (e.g., interactions with other hosts and vectors) that must be separated. For instance, behavior that changes during or at the time of infection is included in infectiousness as is behavior that changes focal host exposure (affecting susceptibility). We keep the host-intrinsic parts of transmission within competence and move its ecological part, the rate of contact with susceptible recipients (θ*_i_*), into host potential (see below). Vector host choice and vector competence belong to the vector half of the transmission cycle. Competence thus is a wholly host quantity, defined relative to a reference recipient whereas measures of transmissibility and host abundance convert this competence into host potential and host capacity.

### Contact rate, potential, and capacity

Again, whether a competent host actually generates new infections depends on how many susceptible recipients it encounters. Those factors are set largely by the density of vectors that bite it, of larval ticks that attach to it, of conspecifics it meets, or of the environments into which it sheds. We denote this transmission-relevant contact rate as θ*_i_* (contacts per unit time).

Individual behaviors that change contact after infection are already contained in b, and individual behavior that changes exposure before infection is contained in ε. Subsequently, θ*_i_* is the ecological baseline against which both are measured. **Host potential** is then:

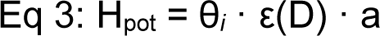

which is dimensionless, expressed as the expected number of new infections that arise from one pathogen exposure of the focal host. **Host capacity**, the contribution of a population of hosts of type *i*, weights potential by the abundance of that type, n_i_, assuming equal exposure across types (Appendix S1, Section 4.6):

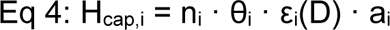

This progression from host competence to potential to capacity mirrors the progression from vector competence to vector potential (Bauer et al. 2026) to vectorial capacity (Garrett-Jones 1964; Smith et al. 2012). It also connects to familiar species-level indices. Specifically, Eq. 2 maps to the reservoir competence index of Komar et al. (2003), Eq. 3 corresponds to indices that weight infectiousness by vector feeding (Kilpatrick et al. 2006), and Eq. 4 relates to reservoir potential and the fraction of infected vectors fed by each host species (LoGiudice et al. 2003; Mather et al. 1989). What is new here is that each term is defined for individuals, derived from a single expression, and given explicit units.

Unlike vectorial capacity, in which the biting rate is squared, contact appears only once in Eq. 4, which covers direct transmission or host-to-vector transmission. Vectorial capacity squares the biting rate because one short-lived vector bites once to acquire the pathogen and again to transmit it, at the same rate and on the same kind of host. For hosts, the contacts that expose a host to a parasite and those behaviors through which it later transmits are generally distinct (or at least separated by significantly more time than vector biting). *Borrelia burgdorferi* (*Bb*), for instance, is acquired by *Peromyscus* mice mainly from nymphs early in summer and transmitted to larvae later in the season. There is even evidence that different vector stages have different host preferences (Brunner & Ostfeld 2008). Also, infection itself can change host behavior. Eq. 1 captures these effects through b. On the other hand, covariance among terms comprising competence is probably common and sometimes even strong (see below): if individuals that are exposed often also become very infectious, their contributions to epidemics could be exceptional (Dye & Hasibeder 1986; Woolhouse et al. 1997). The squared biting rate of vectorial capacity is the special case in which exposure-side and transmission-side contact rates are identical and perfectly correlated. Whereas over a full host–vector–host cycle the biting rate still appears twice, once in θ*_i_* and once in V (Eq. 6), what does not carry over to hosts (compared to vectors) is the product of a host’s own exposure and transmission contacts, which is more plausible given the longer lives and more complex behaviors of hosts.

### General form and the special case used here

Eq. 1 above is general. In our examples below, we use a special case of it under three assumptions. (A1) Behavior does not change with load or infection age, b(τ) ≡ b, so b leaves the integral. This case removes the dependence of sickness behavior on pathogen load, but it retains individual differences in behavior or susceptibility. (A2) Infectiousness is lost only through death (γ ≡ 0), so Σ(τ) becomes the survival function S(τ), and clearance of pathogens in acute infections is represented by load itself returning to zero. (A3) Infectiousness is a Hill function of load, φ(L) = L^n^/(K^n^ + L^n^), with half-saturation load K and steepness n (Almocera et al. 2018; Ke et al. 2021; Marc et al. 2021). The Hill form is a phenomenological default, chosen because it represents well the load-infectiousness relationships in some of our example systems; φ can also be taken from a dose–response curve, because a donor’s load sets the dose delivered to a recipient (e.g., φ(L) = 1 − e^−rL^; Brouwer et al. 2017; Lunn et al. 2019) or derived from within-host models (Della Marca et al. 2023; Gubbins 2024). Given A1–A3 (the mechanisms that bound the integral in different infections are described in Appendix S1, Section 7), infectiousness is expressed as:

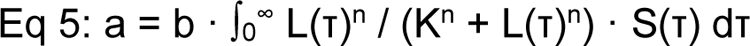

### Relation to reproduction numbers

Infectiousness (*a*) is not a reproduction number, so *a* = 1 has no meaning on its scale. A threshold appears only once contact is included and the transmission cycle is closed. For an infected individual of type *i*, the expected number of new infections it generates is:

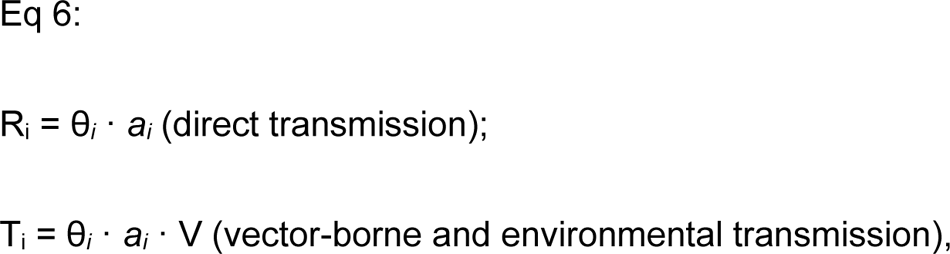

Where R_i_ is the individual reproduction number of Lloyd-Smith et al. (2005), whose mean over newly infected hosts, weighted by exposure and susceptibility, is R_0_ (Appendix S1, Section 4.6). Recipient susceptibility is contained in φ, so the focal host’s own ε does not appear. For directly transmitted pathogens, R_i_ = 1 separates individuals that more than replace themselves from those that do not. For vector-borne and environmentally transmitted pathogens, θ*_i_* · *a* counts infected vectors, or pathogens deposited in the environment, but this step is only half of the cycle. Self-replacement then requires multiplying by V, the expected number of host infections produced per infected vector (or per unit of environmental pathogen). The result, T_i_, is the number of host infections produced through one host-vector-host cycle; for identical hosts it is a type-reproduction number (Hartemink et al. 2008; Roberts & Heesterbeek 2003), and T = 1 is exactly equivalent to R_0_ = 1 because R_0_ = √T. Because θ*_i_* and V are rarely measured at the individual host level, we display thresholds in two ways below: i) the fraction of individuals that would replace themselves across a range of effective contact rates, and, ii) for Lyme disease, where larval tick burdens provide θ*_i_* directly. Like R_0_, R_i_ and T_i_ describe invasion into a largely susceptible population and do not account for the depletion of susceptibles at endemic equilibrium. If contact rates or pathogen load trajectories vary with time of year rather than with infection age, though, the appropriate object is a time-periodic reproduction number (Bacaër & Guernaoui 2006), making our formulation a special case.

Full derivations, including the next-generation formulation from which Eq. 6 follows (Diekmann et al. 1990, 2010; van den Driessche 2017), are given in Appendix S1.

## 3. Quantifying host competence among and within disease systems

To probe the promise of this form of individual competence, we parameterized Eq. 5, then simulated host populations of three different host species for four disease systems. We focused on species for which data on susceptibility, within-host load trajectory, Hill parameters, and survival were available; where specific parameters were not well defined or unavailable, we used biologically reasonable assumptions. For West Nile virus, species-mean daily viremia and among-individual ranges came directly from experimental infections (Komar et al. 2003). For Lyme disease, susceptibility and pathogen load for species were drawn from estimates in individuals from field studies (Barbour et al. 2015; Brunner et al. 2008; Keesing et al. 2009; LoGiudice et al. 2003). In *Bd*, loads and load-dependent mortality were inferred from empirical observations (Briggs et al. 2010; Knapp et al. 2022; Ribeiro et al. 2019; Vredenburg et al. 2010). For avian influenza, experimental data on load and shedding provided reasonable estimates (Brown et al. 2012; Costa et al. 2011). Other parameter values were chosen to be plausible based on the literature; among-individual variances were drawn from the literature sources as reported in Appendix S2. To translate individual-level variation into intraspecific distributions of host competence, we used Monte Carlo simulation, drawing 4,000 individuals for each host species. Each parameter was drawn from its distribution (Table 1), and competence was computed from the drawn values. The full list of parameters, values, distributions, and sources is given in Appendix S2.

**Table 1.** The four disease systems organized by the terms of Section 2. Infectiousness, *a*, depends on the load trajectory, the map from load to infectiousness (φ), and individual behavior (b); susceptibility (ε) and transmissibility (θ*_i_*) are listed separately; the last column gives the mechanism that bounds the infectiousness integral. Values are plausible rather than fitted except where a source is given; full parameter values are in Appendix S2.

| **System<br>(Load x Transmission route)** | **Infectiousness *a*: load trajectory and infectiousness φ** | **Behavior b, susceptibility ε, transmissibility θ<sub>*i*</sub>** | **How the infection ends** |
|---|---|---|---|
| West Nile Virus<br>(acute x vector-borne) | Daily mean log<sub>10</sub> viremia from Komar et al. (2003) with an individual offset (normal; SD from reported ranges) and viremia duration (truncated normal); φ = Hill on log<sub>10</sub> titer (half-saturation 10<sup>8.79</sup> PFU/mL, n = 4.5; zero below 10<sup>5</sup>) | b lognormal (SD 0.15, assumed); ε = 1 (all exposed birds infected; Komar et al. 2003); θ<sub>*i*</sub>V = mosquito biting rate (unmeasured; varied in Fig. 1B); infection of the mosquito is in φ; dissemination and transmission are in V | Load route: viremia clears (or the bird dies) within 2–6 days |
| Lyme disease<br>(chronic x vector-borne) | Plateau spirochete load (lognormal), logistic growth rate and inflection time (normal); φ = Hill (K = 10<sup>4</sup>, n = 1.5) | b = feeding survival of attached larvae, i.e., grooming (lognormal; Levi et al. 2016); ε fixed per species (0.05–0.95); θ<sub>*i*</sub> = larvae attaching per host per 72 h ÷ 3 (post-grooming larval burden ÷ feeding survival; Levi et al. 2016) | Seasonal window: 60-day larval season; no infection-induced mortality; host survival within the season not modeled |
| *Bd*<br>(progressive x environmental) | Initial load and carrying capacity (lognormal), growth rate (normal); φ = Hill (K = 10<sup>4</sup> zoospore equivalents, n = 2.5) | b lognormal; ε beta-Poisson (dose 1,000 zoospores); θ<sub>*i*</sub> = contact rate × zoospore persistence | Lifespan (tolerant bullfrog; mean 5 years) or disease-induced death (load-dependent hazard above a threshold; leopard and mountain yellow-legged frogs) |
| LPAI<br>(acute x direct or environmental) | Peak cloacal shedding (lognormal) and time to peak (normal), acute rise and fall; φ = Hill (K = 10<sup>4</sup> EID<sub>50</sub>/mL, n = 2) | b lognormal; ε beta-Poisson (dose 10<sup>4</sup> EID<sub>50</sub>); θ<sub>*i*</sub> = flock and water contact × viral persistence in water | Load route: shedding clears within about three weeks; non-lethal |

We selected these host-pathogen sets for three reasons: i) they differ in within-host load dynamics (i.e., acute, chronic, or progressive) among host species, ii) each involves different transmission routes (i.e., vector-borne versus direct/environmental; Table 1), and iii) each is either a well-studied multi-host zoonotic pathogen or one of conservation concern. Within one system (chytrid), we deliberately simulated hosts where infection was and was not lethal to identify the effect of pathogen-mediated mortality.

### West Nile virus

West Nile virus is an acute, mosquito-borne flavivirus maintained in North America in cycles between *Culex* mosquitoes and birds. Viremia rises steeply over two to four days and is cleared within about a week, so the infectious period is set by the rise and fall of viremia (Table 1), although some species, notably corvids, often die before clearing infection (Komar et al. 2003). Komar et al. (2003) experimentally infected a wide range of North American bird species with the NY99 strain and summarized each species’ reservoir competence as susceptibility × mean daily infectiousness × duration of infectiousness. From their daily viremia data, we parameterized three focal species spanning the widest range of that index: blue jay (*Cyanocitta cristata*; 2.55), house sparrow (*Passer domesticus*; 1.59) and mourning dove (*Zenaida macroura*; 0.19). We later consider American robin (*Turdus migratorius*; 1.08) and European starling (*Sturnus vulgaris*; 0.22), partly because robins are often regarded as the key amplifying host in eastern North America (Hamer et al. 2009; Kilpatrick et al. 2006). Competence is also a property of the host–pathogen strain combination rather than of the host alone. NY99 was soon displaced by the WN02 and SW03 genotypes. Wild house sparrows collected in 2012–2013 were less competent for NY99 than earlier sparrows, and in those birds, contemporary WN02 and SW03 isolates produced peak viremias about 1.2 log_10_ PFU/mL higher than NY99, a net increase for circulating strains (Duggal et al. 2014). Our parameterization describes competence for NY99 in birds sampled shortly after its introduction.

### Lyme disease

Lyme disease is caused by infection with the bacterium, *Borrelia burgdorferi*, transmitted to hosts by *Ixodes* ticks (Ostfeld et al. 2026). In the white-footed mouse (*Peromyscus leucopus*), the principal reservoir for *B. burgdorferi*, infection is chronic and well-tolerated (i.e., no pathology), with bacterial load persisting near the threshold for transmission to feeding ticks for much of the infectious period (Barbour et al. 2015; Brunner et al. 2008; Keesing et al. 2009). Infected mice suffer little mortality from infection, so the infectious period is bounded not by clearance but by the larval transmission season. Eastern chipmunks (*Tamias striatus*), by contrast, have been described as moderately competent hosts (Brunner et al. 2008; Ostfeld et al. 2026), perhaps because they encounter and/or host many nymphal ticks (Hamer et al. 2012). In contrast, eastern gray squirrels (*Sciurus carolinensis*) appear to have low infectivity to feeding ticks, which at least partly explains their comparatively low competence at the species level (Brunner et al. 2008).

### Chytridiomycosis

Chytridiomycosis, caused by the fungus *Batrachochytrium dendrobatidis* (*Bd*), is a progressive, persistent infection in some tolerant amphibian reservoirs. In other species, infection is lethal, truncating survival and hence transmission (Briggs et al. 2010; Greenspan et al. 2012; Ribeiro et al. 2019). Fungal load accumulates on the skin and is shed into shared aquatic habitat, so transmission occurs primarily into the environment. The American bullfrog (*Lithobates catesbeianus*) lives quite long and tolerates high fungal burden for extended periods, so it acts as a persistent environmental source. For *Bd* in the bullfrog, natural lifespan bounds the infectious period. By contrast in vulnerable hosts, such as the mountain yellow-legged frog (*Rana muscosa*) and the northern leopard frog (*Rana pipiens*), accumulating fungal load is not tolerated and drives mortality. Death thus truncates the load trajectory and shortens the infectious period of individuals in these species.

### Avian influenza

Low-pathogenicity avian influenza (LPAI) is an acute, directly and environmentally transmitted viral infection of waterbirds, in which the infectious period is bounded by the rise and fall of fecal–cloacal shedding rather than by mortality. Competence for LPAI varies markedly across three co-occurring hosts largely due to differences in shedding: the mallard (*Anas platyrhynchos*) is considered a major reservoir. The wood duck (*Aix sponsa*) is an intermediate shedder whereas the laughing gull (*Leucophaeus atricilla*) is a comparatively poor shedder (Brown et al. 2012; Costa et al. 2011; Jourdain et al. 2010). We choose LPAI here because the best individual shedding data are available for this pathogen (Hénaux & Samuel 2011; Latorre-Margalef et al. 2009).

Across all four systems, intraspecific variation in infectiousness was substantial (Fig. 1A). Within species, infectiousness spanned one to two orders of magnitude, and in the least competent species three or more (e.g., laughing gull). Coefficients of variation within host species rose from 0.13–0.45 in the most competent species of each system (white-footed mouse 0.13, blue jay 0.35, mallard 0.43, bullfrog 0.45) to 0.97–2.07 in the least competent species (gray squirrel 0.97, mourning dove 1.08, laughing gull 2.07); the exception was chytridiomycosis, in which all three species varied similarly (0.35–0.45). In every case, our index varied as parameterized, with competence declining from the most-recognized reservoir to the least competent host species. Mean infectiousness fell from the mallard (4.5 effective infectious days) through the wood duck (2.4) to the laughing gull (0.22) in avian influenza; from the blue jay (2.1) and house sparrow (2.0) to the mourning dove (0.33) in West Nile virus, a quarter of which never reached an infectious viremia; from the white-footed mouse (23) through the eastern chipmunk (7.9) to the eastern gray squirrel (1.5) in Lyme disease; and from the bullfrog (1,700) to the northern leopard frog (44) and mountain yellow-legged frog (17) in chytridiomycosis.

**Figure 1.**
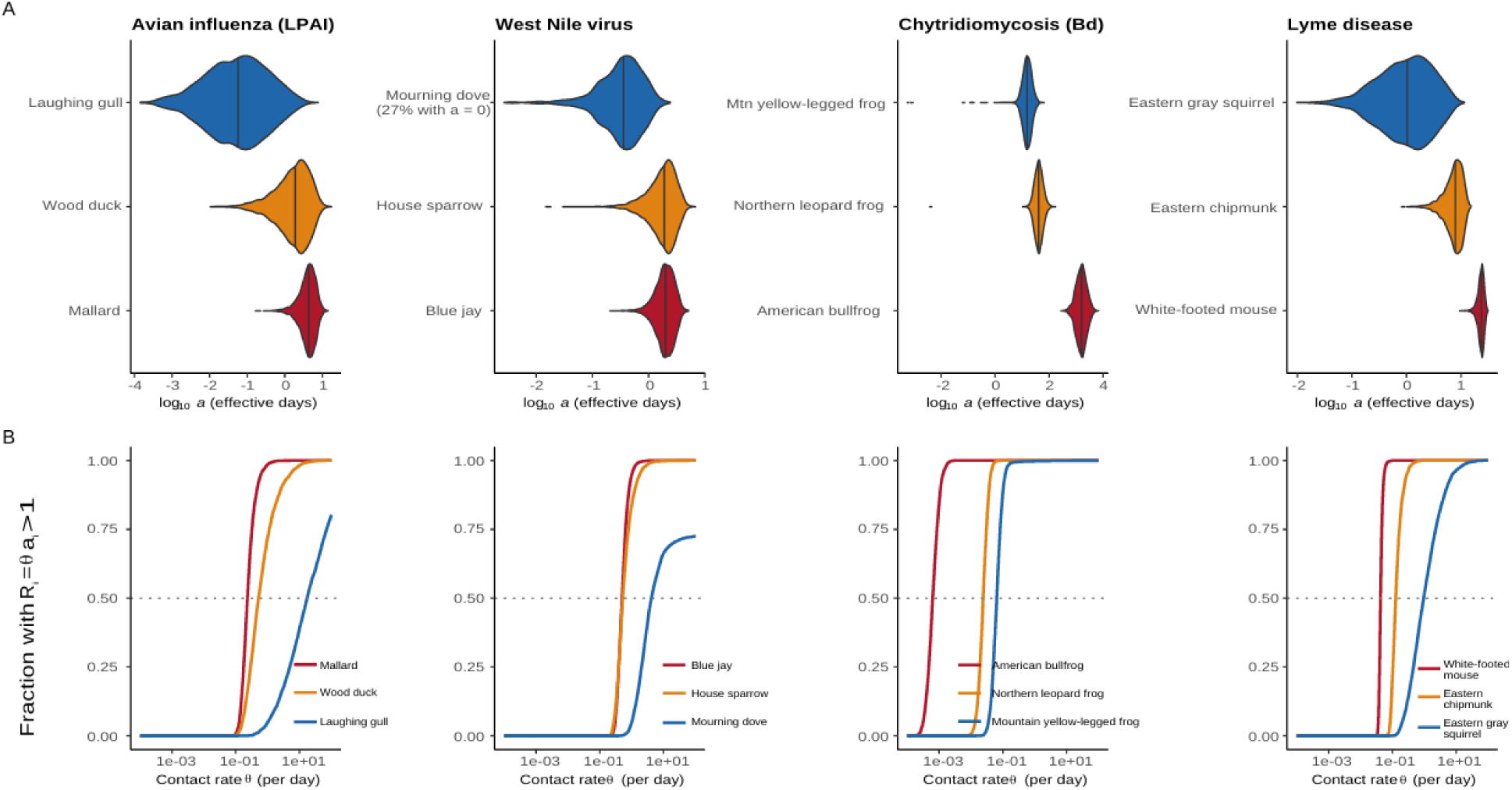
Individual infectiousness and self-replacement in four disease systems. (A) Distributions of infectiousness, *a* (Eq. 5; effective infectious days, log_10_ scale), for 4,000 simulated infected individuals per species; vertical lines within violin plots mark medians. West Nile virus species were parameterized from daily viremia data in Komar et al. (2003), with an assumed among-individual SD of 0.15 (log scale) in behavior, b; individuals whose viremia never reached 10^5^ PFU/mL have *a* = 0, and their percentage is given beside the species name. Lyme disease is integrated over a 60-day larval-tick season. (B) Fraction of infected individuals whose individual reproduction number, R_i_ = θ*_i_* · a_i_, exceeds 1 across a range of effective contact rates, θ*_i_*, assuming all individuals share the same contact rate. Here θ*_i_* (θ*_i_*V in Eq. 6) is the rate of transmission-relevant contacts that each generate a new host infection, so for vector-borne and environmental systems it includes the vector or environmental half of the cycle. The contact rate at which a curve crosses 0.5 is the rate at which the median individual replaces itself; the steepness of each curve reflects within-species variation.

Where individuals fall relative to the threshold for self-replacement is even more revealing (Fig. 1B). Species differed quite a bit in the breadth of infectiousness. Highly competent species were competent because of a high mean and low variance, whereas low-competence species were on average less competent but also more variably so. The ecological ramifications of these patterns become obvious when intraspecific variation is considered relative to the individual threshold. For instance, in avian influenza, the median mallard would replace itself once its effective contact rate exceeded about 0.23 per day and the median wood duck above about 0.5 per day. The median laughing gull, however, would need about 18 effective contacts per day; at a given contact rate, most gulls would therefore dampen transmission, but some gulls (in the upper tail of the distribution) could cause more than one infection under conditions in which the typical gull could not. Similarly for West Nile virus, the median blue jay and house sparrow would replace themselves at similar contact rates (about 0.5 per day), but the median mourning dove would do so only at about 4 per day and a quarter of doves never would. In Lyme disease and chytridiomycosis, where infections persist for months to years, break-even contact rates were one to three orders of magnitude lower (e.g., about 0.04 per day for the median white-footed mouse and 0.0006 per day for the median bullfrog). Chytridiomycosis is the only system studied here in which survival, whether natural lifespan (bullfrog) or disease-induced death (leopard and mountain yellow-legged frogs), bounds the infectious period. This disease highlights some important caveats for host competence. First, intraspecific variance in competence will probably differ among systems. In the case of chytrid, the average individual in a species is representative of most individuals. This outcome arises because fungal loads in all three species rise far above the threshold for infectiousness, so differences among individuals in load matter little; what varies is survival and, under our assumed spread, behavior.

Our individual-based results for West Nile virus (WNV) also compare reasonably well with established species-level metrics but with some important caveats. The means of our simulated individuals mostly agreed with Komar et al.’s (2003) species ranking, as expected because our Hill function was calibrated to their data (Fig. 3C): blue jay (2.1 vs 2.55), house sparrow (2.0 vs 1.59), American robin (0.90 vs 1.08), European starling (0.25 vs 0.22) and mourning dove (0.33 vs 0.19). However, because our index derives from individual measures, we revealed that individuals within species may be quite distinct in their propensity to cause new infections. For instance, the most infectious 20% of jays supplied about a third of their species’ transmission but about half of all infections in doves came from this same 20%. It also emphasizes that host species can play comparable roles in the same system through different pathways. Robins, for example, rank below house sparrows in within-host infectiousness, yet they are widely regarded as key amplifying hosts because *Culex* feed on them disproportionately (Hamer et al. 2009; Kilpatrick et al. 2006).

A notable finding from our simulations is that the duration of the infectious period strongly influences the extent of variation in infectiousness within but also among species. Infectiousness values differ by roughly five orders of magnitude across systems, being lowest in acute systems where the infectious period spans days (i.e., avian influenza, West Nile virus) and higher in systems where infections can be tolerated for years in some reservoirs (i.e., Lyme disease, chytridiomycosis). These differences partly reflect our choice of transmission window (i.e., a 60-day larval season for Lyme disease, natural lifespan for bullfrogs), and because contact rates are unmeasured in most systems, magnitudes cannot be compared across systems, limiting comparisons of competence to within systems.

## 4. Covariation among parameters underlying competence

In the simulations above, each component of competence was drawn independently. Under the known and estimated variances, a variance decomposition of those simulations (Saltelli et al. 2010; Appendix S3) showed that load magnitude explained most among-individual variation in infectiousness in every system except chytridiomycosis, and more so in low-competence species (first-order Sobol indices 0.84–0.95) than in high-competence ones (0.52–0.76), because the loads of low-competence species sit near the threshold at which small differences in load produce large differences in infectiousness; in chytrid frogs, natural lifespan or behavior dominated instead. However, covariation between aspects of competence is probably common and worthy of consideration (Vazquez-Prokopec et al. 2016). Behavioral syndromes can couple susceptibility and peak load in avian mycoplasmosis (Hawley et al. 2011), sickness behaviors can depress transmissibility of a variety of bacterial and other pathogens at high loads (Adelman & Martin 2009), and prior immune experience can reduce infectiousness and favor host survival (Tarasiuk et al. 2023). Because transmission depends on the product of infectiousness and behaviors affecting contact once infected, the covariation most likely to matter is between them. We therefore let each individual’s contact rate vary (CV = 1; Tufts et al. 2021) and linked it to load magnitude with a bivariate Gaussian copula (Nelsen 2006), which leaves each marginal distribution unchanged, for correlations from −0.7 (heavily infected individuals make fewer host- or vector-directed contacts) to +0.7 (heavily infected individuals make more). For a high- and a low-competence species in each system, we then compared the resulting distributions of individual potential, θ_i_ · a_i_ (Fig. 2A–D), focusing on their upper tails: the percentage of individuals above the 99th percentile of the uncorrelated case (Fig. 2E), the percentage that would replace themselves at a mean contact rate at which 10% do so without covariation (Fig. 2F), and the share of transmission contributed by the most infectious 10% of individuals (Fig. 2G).

**Figure 2.**
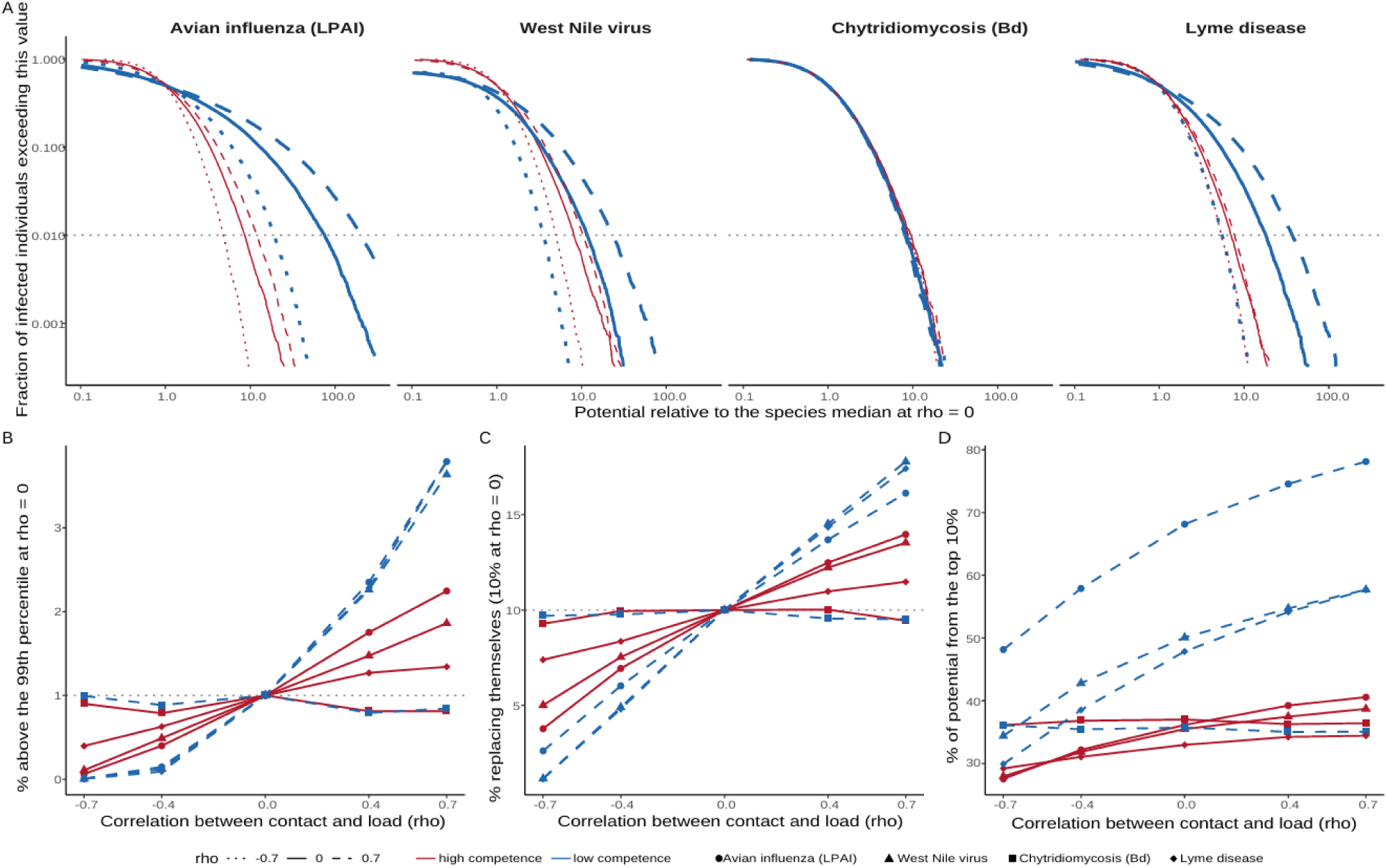
Covariation between contact and load reshapes the upper tail of transmission potential in low-competence species. Each individual’s contact rate, θ_i_ (among-individual CV = 1, similar to larval tick burdens on white-footed mice; Tufts et al. 2021), was linked to its load magnitude (peak, plateau or carrying-capacity load; for West Nile virus, the individual log_10_ viremia offset) by a Gaussian copula with correlation ρ, leaving both marginal distributions unchanged; each scenario comprises 40,000 simulated infected individuals per species (8,000 for the bullfrog). (A–D) Exceedance curves: the fraction of infected individuals whose potential, θ_i_ · a_i_, exceeds a given value, expressed relative to the species median at ρ = 0, for a high-competence (red, thin) and a low-competence (blue, thick) species in each system, with ρ = −0.7 (dotted; heavily infected individuals make fewer transmission-relevant contacts), 0 (solid; independent) and +0.7 (dashed; heavily infected individuals make more transmission-relevant contacts). Both axes are logarithmic, so the upper tail lies at the lower right; the gray dotted line marks 1% of individuals. (E) Percentage of individuals above the 99th percentile of the uncorrelated (ρ = 0) distribution, which is 1% by definition at ρ = 0. (F) Percentage of individuals that would replace themselves (R_i_ = θ_i_ · a_i_ > 1) at the mean contact rate at which 10% do so when ρ = 0. (G) Share of each species’ total potential contributed by its most infectious 10% of individuals. In E–G, red solid lines with filled symbols denote high-competence species and blue dashed lines with open symbols low-competence species; symbols identify systems.

Covariation between contact and load reshaped the upper tail of transmission potential far more than its center, and it did so mainly in low-competence species (Fig. 2). When the most heavily infected individuals also made the most contacts (ρ = +0.7), the percentage of laughing gulls, mourning doves and gray squirrels in the extreme upper tail (above the 99th percentile of the uncorrelated case) rose from 1% to 3.7–4.0%, and the 99th percentile itself rose 1.9–2.7-fold. When the most heavily infected individuals contacted least (ρ = −0.7), that tail all but disappeared (<0.05% of individuals) and the 99th percentile fell three- to fourfold (Fig. 2A, B, D, E). These shifts changed which individuals could sustain transmission. At a mean contact rate at which 10% of individuals replaced themselves without covariation, 16–17% of gulls, doves and squirrels did so under positive covariation, but only 1–2% under negative covariation (Fig. 2F). Likewise, the most infectious 10% of gulls, doves and squirrels supplied 57–78% of their species’ potential under positive covariation, but 30–48% under negative covariation (Fig. 2G). High-competence birds responded less (mallard and blue jay: 2.0–2.2% of individuals in the extreme tail and 14% self-replacing at ρ = +0.7), and species whose loads saturated infectiousness hardly at all (white-footed mouse: 1.3% and 11%; bullfrog and mountain yellow-legged frog: little changed (0.68–1.1%)). Mean potential changed correspondingly, by ×0.41–2.14 in laughing gulls, ×0.52–1.67 in mourning doves and ×0.57–1.58 in gray squirrels across ρ = −0.7 to +0.7, but by only ×0.81–1.17 in mallards and blue jays and ×0.93–1.05 in white-footed mice.

## 5. Interspecific variation in intraspecific variation in competence

The exercises above are solely theoretical. What would justify future work on H_comp_ based on individual hosts is evidence that it agrees with existing species-level measures and provides information they cannot. Unfortunately, there are relatively few multi-host pathogens for which extensive knowledge about host competence is available. One exception is Lyme disease where years of careful work have generated the necessary data (Brunner et al. 2008; Keesing et al. 2009; Levi et al. 2016; LoGiudice et al. 2003; Ostfeld et al. 2026). We used those data to simulate distributions of infectiousness for 4,000 individuals within 12 vertebrate host species of *Borrelia burgdorferi* using the same approach as Fig. 1. Combining infectiousness with each species’ larval tick burden (θ*_i_*) gave the number of infected larvae each infected host produces per season (Fig. 3A), revealing two interesting results. First, mean infectiousness of each host species tracked published realized reservoir competence for host species (Brunner et al. 2008; LoGiudice et al. 2003) across about two and a half orders of magnitude (R² = 0.79; Appendix S2, Tables S5–S6). This agreement is an internal consistency check rather than an independent validation because species’ susceptibility and loads were parameterized from the same field data. The two competence measures also differ in what they include. Realized reservoir competence is the probability that a larva feeding on a field-caught host becomes infected, so it folds in natural exposure and prevalence but not the number of larvae a host feeds or how long it remains infectious; our index conditions competence on infection and, once combined with tick burdens, counts infected larvae produced per host. That difference changes rankings: the eastern gray squirrel, a poor host by realized reservoir competence, produced about 70% as many infected larvae per infected individual as the eastern chipmunk (medians 283 vs 398), because more than five times as many larvae attach to it.

Second, we found that variation among individuals increased as mean competence decreased: the CV of infected larvae produced rose from 0.12 in the white-footed mouse to 1.7 in the Virginia opossum and 2.4 in white-tailed deer (Fig. 3A). This pattern arose largely because the pathogen loads of low-competence species sit near the threshold of the load–infectiousness curve, where small differences in load produce large differences in infectiousness. The most competent species, by contrast, sit on a saturated upper plateau. In the Lyme and avian influenza parameterizations, it is also partly an input artifact because assigned variances increase toward low-competence species. As hinted by Fig. 1, members of high-competence host species will probably play similar ecological roles for Lyme, however, individuals of low-competence species could be inconsequential or important, depending on context (Gervasi et al. 2015). Fig. 3B makes this more explicit. Every simulated white-footed mouse and nearly every eastern chipmunk, masked shrew and short-tailed shrew (>99%) would replace itself even if only about 1.5% of infected larvae went on to infect a host, as would most gray squirrels (87%). However, most individuals of the other host species would only do so if tick survival and host-finding were high. Even at the upper end of the plausible range (35%), 15% of white-tailed deer and 2% of opossums would not replace themselves. In West Nile virus, similar results hold. The median infectious mourning dove needs more than five times the effective contact rate of the median blue jay to replace itself (2.8 vs 0.49 per day), but the most competent doves match typical jays (Fig. 3D).

**Figure 3.**
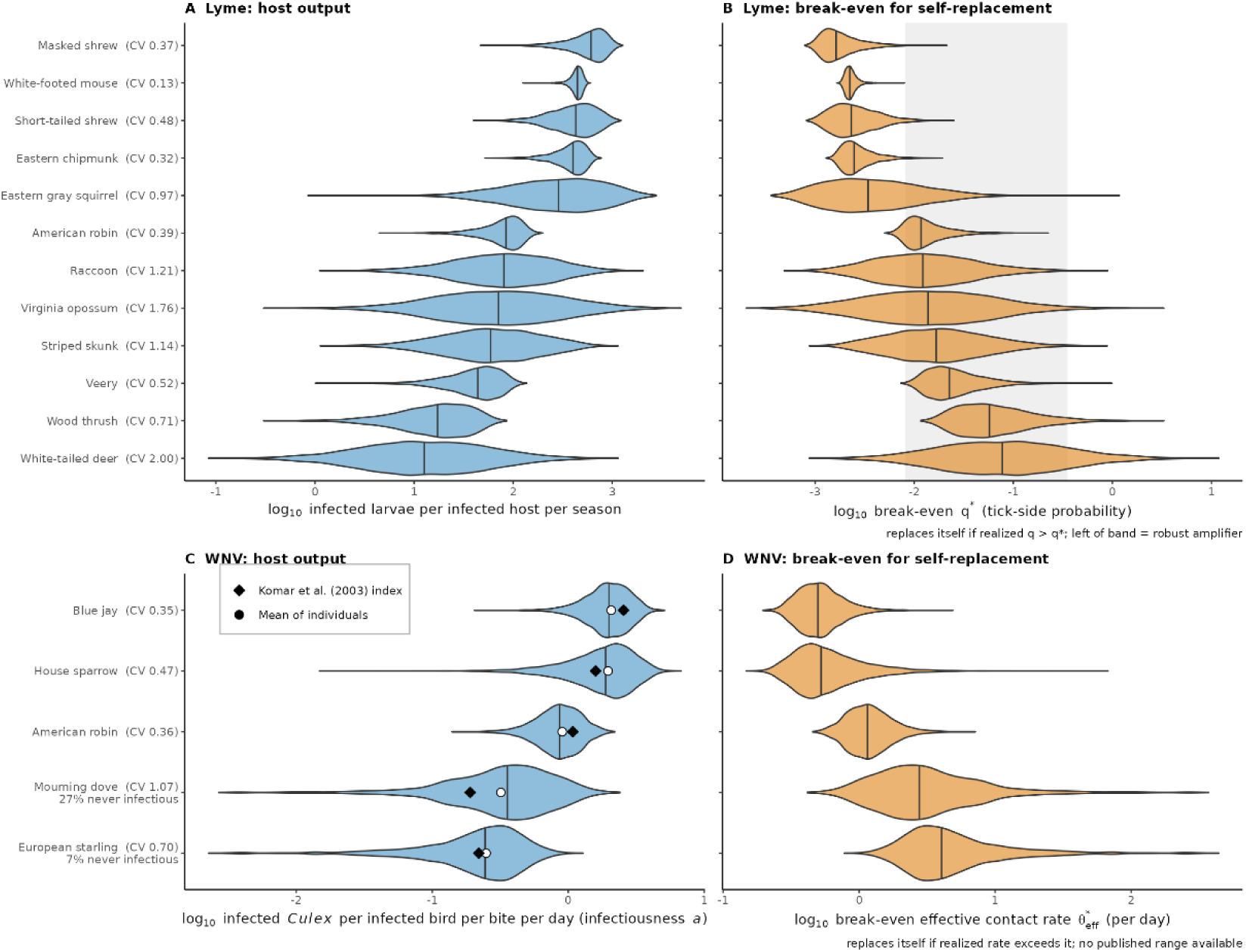
Individual output and self-replacement thresholds in Lyme disease and West Nile virus. (A) Lyme disease: infected larval ticks produced per infected host during a 60-day larval season (*a* · θ*_i_*, with θ*_i_* equal to larvae attaching per host per 72 h divided by 3, i.e., post-grooming burden ÷ feeding survival; Levi et al. 2016), for 4,000 simulated individuals of each of 12 species; coefficients of variation are given beside species names. (B) Break-even probability, q* = 1/(*a* · θ*_i_*), that an infected larva survives to molt and overwinter, finds a host as a nymph and infects it; an individual more than replaces itself if the realized probability exceeds q*. The shaded band (0.8–35%) spans products of published estimates of survival through molting and winter (18–47%; Lindsay et al. 1998), nymphal host-finding (15–83%; Tufts et al. 2021) and an assumed 30–90% probability that the host fed upon is susceptible and becomes infected. Host survival within the season is not modeled, so q* is a lower bound. (C) West Nile virus: infectiousness, *a*, expressed as the number of *Culex* infected per infected bird per mosquito bite per day, for 4,000 simulated individuals of five species spanning the range of the reservoir competence index of Komar et al. (2003). Species-mean daily viremia and among-bird ranges are from Komar et al. (2003, Table 2); individuals differ in their log_10_ viremia offset (SD estimated from reported ranges), duration of viremia and behavior (assumed SD 0.15, log scale); infectiousness to *Culex* is a Hill function of log_10_ viremia calibrated to Komar et al.’s mean daily infectiousness (half-saturation 10^8.79^ PFU/mL, n = 4.5, zero below 10^5^ PFU/mL). Individuals whose viremia never reached 10^5^ PFU/mL are excluded from the violins and their percentage is given beside the species name. Open circles: mean of simulated individuals; diamonds: Komar et al.’s species index. Among-bird variances rest on 2–6 birds per species and are approximate. (D) Break-even effective contact rate, θ*_eff_ = 1/*a*, the number of transmission-relevant bites per day, each weighted by the probability that the mosquito survives the extrinsic incubation period and infects another bird, that an individual needs to replace itself. Unlike panel B, no published range can yet be drawn, because *Culex* biting pressure on individual birds is site- and season-specific.

These results are important for two reasons. The first is mathematical: where competence distributions are widest, a species-level estimate could entail a mis-specified variance in the quantity to which models are most sensitive. There is a general expectation that averaging over individuals will bias species-level competence estimates substantially; often, assuming too little inter-individual variation overestimates transmission rate and epidemic size and makes intervention forecasts systematically pessimistic (Coffeng & de Vlas 2022). Disregarding intraspecific variation is thus inadvisable where trait–outcome relationships are strongly non-linear (Moran et al. 2016; Ruel & Ayres 1999). In our simulations, the resulting bias in species means was modest (Appendix S1, Section 9.1), so the stronger case for individual data rests on covariation (Fig. 2) and on questions that species means cannot answer, such as what fraction of a species’ individuals would replace themselves and how unevenly infected larvae are produced among them (Fig. 3). The same logic identifies where individual-level data will often be most informative: low-competence species whose loads sit near the transmission threshold. The second reason that these results matter is biological. Specifically, some forms of variance in competence can underpin superspreading (Lloyd-Smith et al. 2005). In many systems, a minority of hosts cause most secondary infections (Lloyd-Smith et al. 2005; Meagher & Friel 2022; Woolhouse et al. 1997). This can be particularly important when placed in the context of epidemics and outbreaks (O’Regan & Drake 2025; Taube et al. 2022). In our West Nile virus simulations, the most infectious 20% of individuals supplied about a third of their species’ transmission in blue jays but half in mourning doves. We assume that superspreading might derive from the most competent species, but perhaps this perspective is misguided given Figs 2 and 3. Instead in some cases, an exceptionally competent member of an otherwise poorly competent species may play an outsized role in its community. In this light, we might also more intensively and deliberately investigate covariation among aspects of H_comp_.

## 6. The ecological consequences of variation in individual host competence

The central goal of this paper was to derive and assess the promise of an individual-based metric of host competence. By collecting the appropriate data, one could scale up individual variation in physiology and behavior to higher-order effects, integrating sub-organismal, organismal and ecological processes toward a shared goal. To do so, however, H_comp_ cannot include processes external to the host including interactions between the host, its vectors, other hosts, and its abiotic environment. If host competence is a property of an individual, ecological processes must be kept separate. As with vector competence and vectorial capacity, host competence can be scaled to higher levels via two steps (Eqs. 3–4). First, individual competence can be weighted by among-host contact rates, θ*_i_*, to give host potential (H_pot_), the realized capacity of an individual to transmit a pathogen within a particular environment. Second, host potential can be scaled to relative host abundance, n, giving host capacity (H_cap_), a quantity that describes how much a given host population contributes to transmission. Host potential and capacity can take many forms. Generally, potential connotes the role of a given host individual in reference to its disposition to attract vectors, contact conspecifics, or otherwise transform its infectious potential into actual infections. In Lyme disease, one contributor to potential is larval *Ixodes* tick burden. In West Nile virus, the number of successful blood feeds by *Culex* mosquitoes, the vectors that transmit the virus, would best capture potential. In both cases, to quantify potential for an individual, H_comp_ would be multiplied by one or more of the above measures. Once H_pot_ is quantified, its ecological extension can be assessed more explicitly in one or more forms of host capacity, H_cap_. In most cases, H_cap_ will simply be given by Eq. 4, where n*_i_* is host population density. Of course, n*_i_* can be more nuanced, and may depend on the demographics of a focal host species. Individual differences in behavior are already partitioned among the terms: behavior that changes contact after infection (e.g., aggression in male rodents that transmit hantaviruses sensu Klein et al. (2004)) enters infectiousness through b, behavior that changes exposure enters susceptibility, and the density of recipients encountered enters θ, so many ad hoc modifications will be unnecessary.

Fig. 4 attempts to emphasize the implications of host competence, potential and capacity, using the 12 species from the Lyme disease system. Two things stand out. One is that competence alone does not determine the role of a species in a community – a phenomenon which has been observed in other systems (Johnson et al. 2024). For instance, an abundant host with modest competence for *B. burgdorferi* could pose an outsized risk to the community just because it attracts so many ticks (e.g., the gray squirrel, which rose from eighth in competence to fifth in potential and capacity). Another is that interspecific variation in interactions between hosts and vectors (potential) could be more consequential than competence or host abundance. Rank-order changes in overall risk to the community are most numerous between competence and potential (nine of 12 species); six species changed rank between potential and capacity, four of them among the lowest-ranked hosts, along with the white-footed mouse and masked shrew swapping first and second place. Community capacity, overall, was dominated by the white-footed mouse (36%), masked shrew (25%), short-tailed shrew (21%) and eastern chipmunk (13%) (Fig. 4B). White-tailed deer ranked low throughout; their main role in Lyme ecology comes from feeding adult ticks, and so sustaining tick populations, and perhaps from diverting larval meals away from competent hosts (Huang et al. 2019), effects that lie outside host competence and capacity.

**Figure 4.**
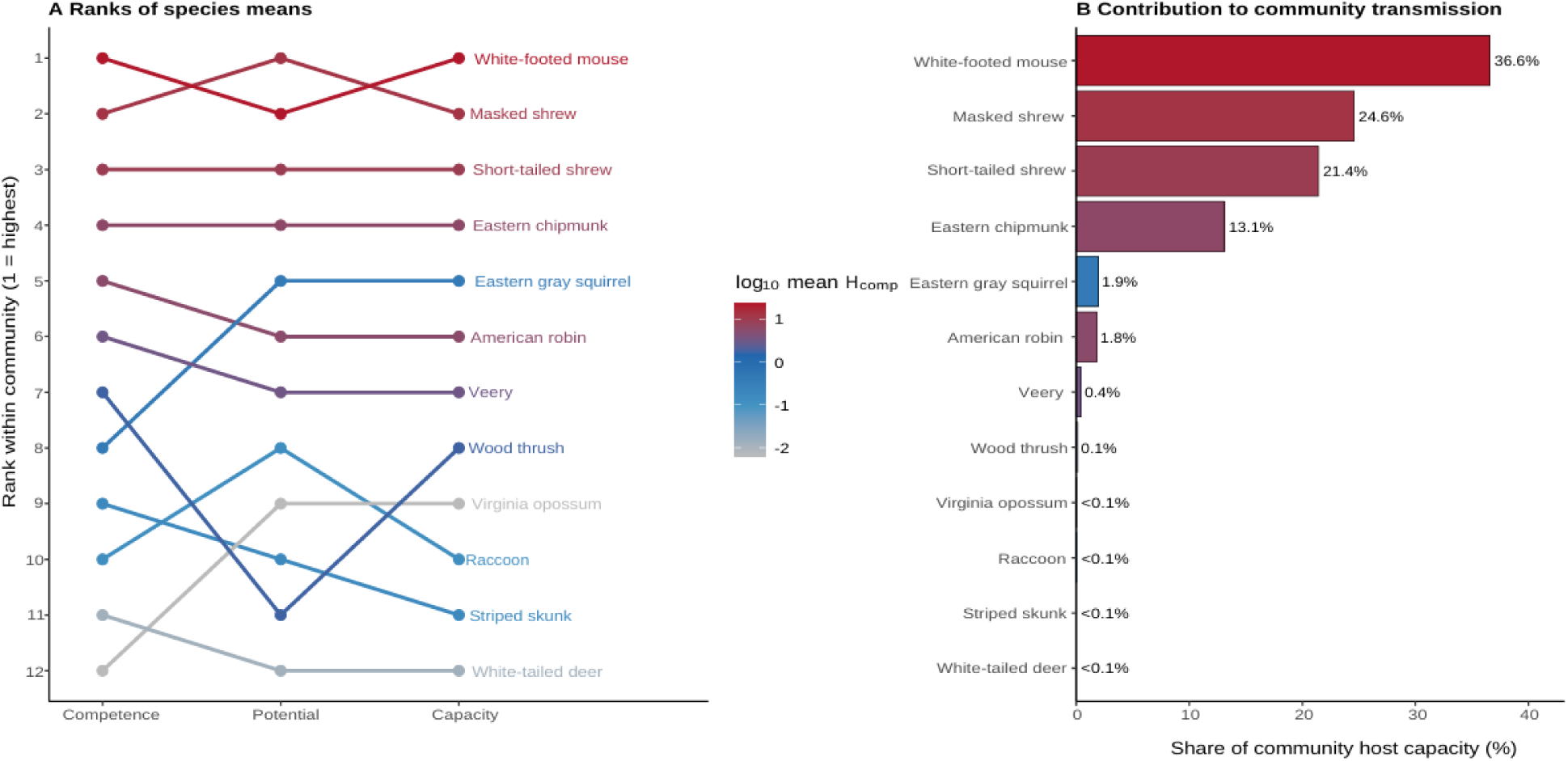
Host competence, potential, and capacity in a typical Lyme disease host community, computed with Eqs. 2–4. (A) Ranks of species means for competence, H_comp_ = ε · *a* (infectiousness over a 60-day larval season multiplied by the probability of infection given exposure); potential, H_pot_ = θ*_i_* · ε*_i_* · *a_i_* (θ = larvae attaching per host per 72 h ÷ 3); and capacity, H_cap_ = n*_i_* · θ*_i_* · ε*_i_* · *a_i_* (n = host density per ha). Colors show log_10_ mean H_comp_. (B) Each species’ share of total community capacity, i.e., of the infected larvae the community would produce if individuals of every species were equally likely to be exposed. Host densities, larval burdens and feeding survival are from Levi et al. (2016, Table 1); ε and loads are plausible rather than fitted values (Appendix S2); contact is counted once (Eq. 4), and nymphal host use is not modeled (see Section 2).

## 7. Further considerations

We have simplified two factors that should be considered more directly in the future: i) pathogen and/or vector encounter rates and ii) time broadly. First, encounters with parasites and vectors are heterogeneous among individual hosts (Brehm et al. 2025; Sih et al. 2018; VanderWaal & Ezenwa 2016). Some hosts are simply in the wrong place at the wrong time, but others have life histories, daily routines, or age-dependent needs that influence whether and how often they encounter pathogens. We decided to define H_comp_ conditional on exposure such that the only way that pathogen encounters could affect H_comp_ was through dose-dependent probability of infection, or susceptibility. As above, we appreciate that this approach might be questionable in some systems, but it was simplifying. More complex options are possible via adjustments of our base equation (Eq. 2); for example, Appendix S1 (Section 4.6) shows how exposure rates that differ among individuals weight their contributions to R_0_.

Second, H_comp_ could be understood as a process rather than a fixed trait. In a chronic infection, for instance, infectiousness may surge and fall (Harris et al. 2023), and when pathogen burden peaks, some behaviors could change, altering transmissibility (Herrera & Nunn 2019; Ruden & Adelman 2021). Likewise, hosts might die before they transmit, reducing their epidemiological effects in some systems but perhaps magnifying them in others, such as parasites that are transmitted only after their host dies (Rapti & Cáceres 2016). Prior pathogen exposure and experiences during development, too, could alter immunological aspects of host competence, typically reducing H_comp_ by lowering susceptibility, pathogen-dependent mortality, or even transmissibility. At the ecological scale, we can partly account for these effects by using thoughtful and natural forms of θ*_i_* and n*_i_*. However, it will also be critical to quantify effects on H_comp_ itself. We will usually not be able to capture all forms of competence in most populations; for instance, extreme competence is rare by nature. With large enough samples, though, one might adequately describe intraspecific heterogeneity. This is also why we separate competence from capacity and potential: H_comp_ describes what an individual host brings to transmission, whereas factors extrinsic to the host, stochastic and deterministic, captured by θ*_i_* and n*_i_*, determine which and how many new hosts and vectors it infects.

We think there are a few additional productive steps to refining H_comp_. Obtaining better susceptibility data is a priority. Susceptibility is markedly difficult to determine as an individual-level trait value, as a single exposure yields a binary outcome and an individual’s dose-response curve can rarely be estimated strictly from that individual. Such data are difficult and expensive to generate, but they are very important. There are presently alternative ways to estimate individual susceptibility from population variances (e.g., Dwyer et al. 1997), but alternative approaches warrant consideration (e.g., individual host cell susceptibility, changes in pathogen burden locally during the acute phase of infections). Another valuable exercise would be to assess the effects of repeated pathogen exposure, immunological memory, co-infection with particular pathogens, or all three on H_comp_. Immunological memory would very likely reshape susceptibility, and aspects of infectiousness, and perhaps even covariation among them. Relatedly, other organismal (e.g., tolerance), physiological (e.g., fever, stress, and acute phase responses), and behavioral (e.g., medication) phenomena could be included as modifiers of aspects of H_comp_. Competence is very likely to be labile within most taxa (Box 1, Figure 5; Buss & Hua 2023; Gervasi et al. 2015), underpinning the potential of some species to transition from incompetent to competent given context.

**Figure 5.**
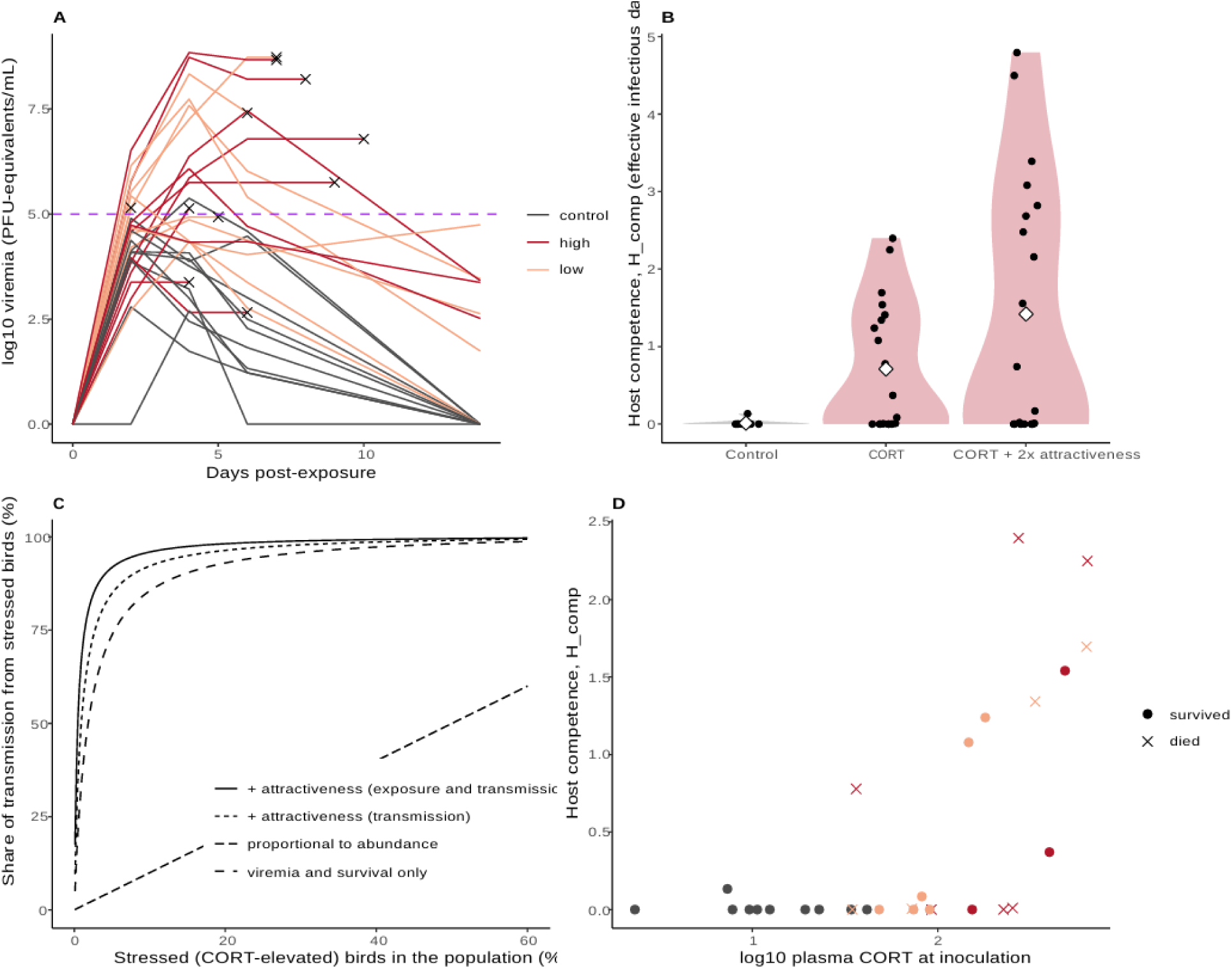
Stress transformed zebra finches from incompetent to competent hosts of West Nile virus. (A) Viremia of 30 zebra finches given empty (control) or corticosterone-filled (low or high dose) silastic implants and inoculated with WNV NY99 (Gervasi et al. 2017a). Viremia was measured by qPCR on days 2, 4, 6 and 14 and interpolated linearly on the log scale; birds that died (×) are held at their last measured titer until death. The dashed line marks the 10⁵ PFU/mL threshold for infecting ***Culex*.** (B) Host competence, H_comp_ = ε · *a*, of each bird (points; diamonds, means). Because all birds became infected, ε = 1 and H_comp_ equals infectiousness, *a*: the integral over the infection of the per-bite probability of infecting a mosquito, a Hill function of log₁₀ viremia (as in Fig. 3, zero below 10⁵). The right-hand group doubles the behavioral term *b* of CORT-implanted birds, reflecting their twofold greater attractiveness to ***Culex quinquefasciatus*** (Gervasi et al. 2016). (C) Share of transmission supplied by stressed birds in a population with a given proportion of CORT-elevated individuals, calculated from the group means in B. Lines show CORT acting through viremia and survival only (dotted), also doubling onward biting (dash-dot), or also doubling the rate at which stressed birds are bitten and hence infected (solid); the **gray** line is the share expected from abundance alone. (D) Plasma CORT at inoculation predicts individual H_comp_; X, birds that died. Colors as in A.

## 8. Conclusion

Our framework builds on robust ideas in disease ecology, vector biology, and epidemiology, to describe a measure of host competence amenable to most systems. Since at least 1999, there has been recognition of the need but lack of such a tool (Spielman 1999). Part of our inspiration, of course, comes from Anderson & May (1986) who described how the acquisition, persistence, and spread of a pathogen in a population is affected by biological properties and heterogeneity in transmission among hosts. Indeed, R_0_ represents for pathogens in a wholly susceptible population what we hope for with H_comp_ in similar contexts. We are sure that the forms of H_comp_ we propose here will not fit every system, but we believe the simple equation to be portable to most. At least, it will focus our attention on variation in individual host physiology and behavior most likely to have the strongest ecological consequences. It is costly to generate dose–dependent susceptibility data and viral, bacterial, and fungal load trajectories as well as pathogen-load dependent mortality rates for many host species, much less enough individuals. However, H_comp_ makes explicit why doing so has value. Importantly, H_comp_ could even inform us how we conduct some immunological research. Species in which H_comp_ is highly variable would be intriguing models to understand the mechanisms driving extreme competence and superspreading (Paull et al. 2011). Now, immunological research tends to proceed from the perspective of maximizing host health. Surely, H_comp_ has many shortcomings, but it is quantitatively defined for individuals, decomposable into measurable organismal traits, integrable over the within-host trajectory, linked to reproduction numbers, and pliable enough to suit most disease systems. Disease ecology is already among the most integrative forms of biology. We think formal, mathematical tools like H_comp_ will foster yet more progress.

## Acknowledgements

We thank the numerous colleagues and friends who have helped to shape this manuscript through great discussion and interest in the broader area of host competence. This work was supported by the National Science Foundation (2110070) and in part by the NSF Research Coordination Network “Integrating Organismal Biology into NEON” (2110233). The authors declare no conflicts of interest. During manuscript preparation, the authors used Claude (Anthropic; Claude Opus models) to help with 1) internal consistency checks for the math underlying equation derivation and 2) prepare, edit, and clean the R code used for simulations and figures. The tool was not used to generate data, results, citations, or scientific claims, and code edits did not alter simulation parameter values, model structure, or the data underlying any figure. NLG reviewed and verified all AI-assisted content and take full responsibility for the content of this publication.

## Statement of authorship

NLG, KMM, CM, KMY, and LBM designed the initial study; all authors conceptualized the final scope of the research. NLG, JMD, TESM, and LBM determined methodological approaches. NLG, JMD, and LBM performed derivation work. NLG and LBM analyzed data and generated figures. NLG wrote the initial draft of the paper, with all authors subsequently providing substantial revisions and feedback on the eventual final draft.

## Data

All data used in this research are openly available from the publications cited in the primary text and in Appendix S2. The code to run all simulations and generate figures is available via Figshare (https://doi.org/10.6084/m9.figshare.34747779). Data for Box 1 are available via the Dryad Digital Repository (https://doi.org/10.5061/dryad.1s9g1).

## Box 1: Implementing H_comp_ to understand the effects of physiological stress on passerine competence for West Nile virus

To illustrate how H_comp_ can be estimated from individual data, we re-analyzed two experiments on zebra finches (*Taeniopygia guttata*), an Australian species with no natural role in North American WNV transmission. In the first (Gervasi et al. 2017a), 30 adult finches received subcutaneous silastic implants that were either empty (controls, n = 10) or filled with corticosterone (CORT) at a low or high dose (n = 10 each). Two days later, all birds were inoculated with WNV NY99. Plasma CORT was measured at inoculation, viremia was quantified by qPCR on days 2, 4, 6 and 14, and survival was followed for 14 days. In a second experiment (Gervasi et al. 2016), other CORT-implanted and control finches were offered to *Culex quinquefasciatus* mosquitoes for blood-feeding, both in choice trials and housed alone. CORT-treated birds were almost twice as likely to be fed upon by mosquitoes, even though they behaved more defensively. We computed each bird’s infectiousness, *a* (Eq. 5), in three steps. First, we interpolated its log₁₀ viremia linearly between samples, holding birds that died at their last measured titer until death; this reproduces the published durations of infectiousness exactly, because they derive from the same samples. Second, we converted titer to the per-bite probability of infecting a mosquito with the Hill function used for WNV in the main text, which is zero below 10⁵ PFU/mL. Third, we integrated over the infection. Because every bird became infected (because we gave a high dose of WNV), susceptibility was 1 and H_comp_ = *a*. We represented greater attractiveness by doubling the behavioral term *b* for CORT birds. Finally, we used group means to estimate the share of transmission stressed birds would supply in a mixed population, first with attractiveness acting only on onward biting and then also on the rate at which stressed birds were themselves bitten and infected.

We found that stress converted an epidemiologically irrelevant host into a competent one. Only one of ten control birds ever exceeded the WNV transmission threshold, and only for about 1.5 days (Fig. 5 A, B). Among CORT-implanted birds, though, 13 of 20 birds crossed the 10^5^ threshold, some for nearly eight days, and mean H_comp_ rose 53-fold, to 0.71 (0.54 and 0.88 for the low and high doses); doubling attractiveness doubled it again (106-fold overall, by construction). Mortality did not offset these gains. The 11 CORT birds that died had higher H_comp_ (mean 0.90) than the nine that survived (0.48), because most were highly viremic for days before death, a trade-off the integral captures but a comparison of mortality rates would miss. Stress also raised the variance of H_comp_: seven CORT birds never became infectious, the coefficient of variation of H_comp_ was 1.14 (3.0 in controls), and the four most competent birds (20%) accounted for 56% of the group’s total H_comp_. CORT concentration at inoculation also predicted individual H_comp_ (Spearman ρ = 0.67, P < 0.001; Fig. 5 D).

All else equal, we found that stressed birds could dominate WNV transmission even when rare. If 5% of a population were stressed, these few birds would supply 74% of transmission through viremia and survival alone, and 92% if attractiveness also raised their exposure (Fig. 5 C). These results mirror the mean–variance pattern we detected in the other systems (Figs 1, 3). The unstressed zebra finches here behave like the low-competence species whose loads sat just below the transmission threshold, where small physiological shifts decided whether an individual transmits at all. An environmental factor (e.g., extreme weather, anthropogenic pollution) could therefore tip individuals across the threshold and produce a superspreader phenotype. For such hosts, species averages, especially measured under benign conditions, could understate the potential role of some birds. Intriguingly in this system, a simply measured individual trait, circulating corticosterone, served as a biomarker of competence.

## Box 2: Glossary

Host competence - H_comp_ = ε · *a*: the expected number of effective infectious days an individual host generates per exposure; the product of susceptibility and infectiousness.

Susceptibility - ε(D): the probability that a host becomes infected when exposed to dose D, including behavior that changes the dose received.

Infectiousness - *a*: effective infectious days, the integral over an infection of the per-contact probability of transmission at the current load, weighted by survival and by infection-induced changes in behavior (b).

Transmissibility - θ*_i_*: the rate of transmission-relevant contacts between a host and susceptible recipients (vectors, conspecifics or a contaminated environment); an ecological rather than an organismal property.

Host potential - H_pot_ = θ*_i_* · ε*_i_* · *a_i_*: the expected number of new infections arising from one exposure of the focal host.

Host capacity - H_cap_ = n*_i_* · θ*_i_* · ε*_i_* · *a_i_*: the contribution of a population of hosts to transmission.

Individual reproduction number - R_i_ = θ*_i_* · a_i_: the expected number of new infections caused by an infected individual; for vector-borne and environmental pathogens, multiplying by the vector- or environment-side term V gives the type reproduction number, T_i_.

Type reproduction number - T*_i_*: the expected number of infections of hosts of type *i* caused, through one full host–vector–host (or host–environment–host) cycle, by one infected individual of that type (Roberts & Heesterbeek 2003); T = 1 is equivalent to R_0_ = 1.

Next-generation matrix - the matrix whose element K_ji_ is the expected number of new infections of type *j* caused by one infected individual of type *i* over its infectious period; its dominant eigenvalue is R_0_ (Diekmann et al. 2010).

Reservoir - one or more epidemiologically connected populations or environments in which a pathogen can be permanently maintained and from which infection is transmitted to a defined target population (Haydon et al. 2002); a population-level role, distinct from the competence of individual hosts.

Superspreader - an infected individual that causes many more secondary infections than the average infected individual (Lloyd-Smith et al. 2005).

Extreme competence - competence far above the norm of a population or species, as in keystone hosts that contribute disproportionately to transmission (Martin et al. 2019).

Resistance - a host’s ability to prevent infection or limit pathogen load; in this framework, resistance lowers ε or *a*.

Tolerance - a host’s ability to limit the damage caused by a given pathogen load; by prolonging survival at high load, tolerance can raise *a*.

Symbols used in Eqs. 1–6 - τ, infection age; L(τ), within-host pathogen load; φ(L), the probability that a transmission-relevant contact at load L infects a reference recipient, here a Hill function with half-saturation load K and steepness n (distinct from abundance, n*_i_*); Σ(τ), the probability of remaining alive and infected, with mortality hazard µ and clearance hazard γ (S(τ) when infections end only through death); b(τ), contact by an infected individual relative to its rate before infection; D, dose; n*_i_*, abundance of hosts of type *i*; V, host infections produced per infected vector or per unit of environmental pathogen. A complete symbol table is given in Appendix S1, Section 2.

## References

Adelman, J. S., & Martin, L. B. (2009). Vertebrate sickness behaviors: adaptive and integrated neuroendocrine immune responses. Integrative and Comparative Biology, 49(3), 202–214.

Almocera, A. E. S., Nguyen, V. K., & Hernandez-Vargas, E. A. (2018). Multiscale model within-host and between-host for viral infectious diseases. Journal of Mathematical Biology, 77(4), 1035–1057.

Anderson, R. M., & May, R. M. (1986). The invasion, persistence and spread of infectious diseases within animal and plant communities. *Philosophical Transactions of the Royal Society of London. Series B*, Biological Sciences, 314(1167), 533–570.

Bacaër, N., & Guernaoui, S. (2006). The epidemic threshold of vector-borne diseases with seasonality: the case of cutaneous leishmaniasis in Chichaoua, Morocco. Journal of Mathematical Biology, 53, 421–436.

Barbour, A. G., Bunikis, J., Fish, D., & Hanincová, K. (2015). Association between body size and reservoir competence of mammals bearing Borrelia burgdorferi at an endemic site in the northeastern United States. Parasites & vectors, 8(1), 299.

Bauer, A. M., Burkett-Cadena, N. D., Reeves, L. E., Alto, B. W., & Campbell, L. P. (2026). Vector potential index: Bridging competence and contribution as an integrative measure of relative transmission capability. Ecology and Evolution, 16(1), e72705.

Becker, D. J., Seifert, S. N., & Carlson, C. J. (2020). Beyond infection: integrating competence into reservoir host prediction. Trends in Ecology & Evolution, 35(12), 1062–1065.

Brehm, A. M., Assis, V. R., Martin, L. B., & Orrock, J. L. (2025). Individual variation underlies large-scale patterns: Host conditions and behavior affect parasitism. Ecology, 106(1), e4478.

Briggs, C. J., Knapp, R. A., & Vredenburg, V. T. (2010). Enzootic and epizootic dynamics of the chytrid fungal pathogen of amphibians. Proceedings of the National Academy of Sciences of the United States of America, 107(21), 9695–9700.

Brouwer, A. F., Weir, M. H., Eisenberg, M. C., Meza, R., & Eisenberg, J. N. (2017). Dose-response relationships for environmentally mediated infectious disease transmission models. PLoS computational biology, 13(4), e1005481.

Brown, J. D., Berghaus, R. D., Costa, T. P., Poulson, R., Carter, D. L., Lebarbenchon, C., & Stallknecht, D. E. (2012). Intestinal excretion of a wild bird-origin H3N8 low pathogenic avian influenza virus in mallards (*Anas platyrhynchos*). Journal of Wildlife Diseases, 48(4), 991–998.

Brunner, J. L., LoGiudice, K., & Ostfeld, R. S. (2008). Estimating reservoir competence of Borrelia burgdorferi hosts: prevalence and infectivity, sensitivity, and specificity. Journal of medical entomology, 45(1), 139–147.

Brunner, J. L., & Ostfeld, R. S. (2008). Multiple causes of variable tick burdens on small-mammal hosts. Ecology, 89, 2259–2272.

Buss, N., & Hua, J. (2023). Host exposure to a common pollutant can influence diversity–disease relationships. Journal of Animal Ecology, 92(11), 2151–2162.

Coffeng, L. E., & de Vlas, S. J. (2022). Predicting epidemics and the impact of interventions in heterogeneous settings: standard SEIR models are too pessimistic. Journal of the Royal Statistical Society Series A: Statistics in Society, 185(Supplement_1), S28–S35.

Cortez, M. H., & Duffy, M. A. (2021). The context-dependent effects of host competence, competition, and pathogen transmission mode on disease prevalence. The American Naturalist, 198(2), 179–194.

Costa, T. P., Brown, J. D., Howerth, E. W., & Stallknecht, D. E. (2011). Variation in viral shedding patterns between different wild bird species infected experimentally with low-pathogenicity avian influenza viruses that originated from wild birds. Avian Pathology, 40(2), 119–124.

Della Marca, R., Loy, N., & Tosin, A. (2023). An SIR model with viral load-dependent transmission. Journal of Mathematical Biology, 86, 61.

DeMarchi, J. A., Carter, E. D., Gray, M. J., Lutz, K. R., Schrock, S. A., Searle, C. L., … & Wilber, M. Q. (2026). Many roads to reservoirs? How susceptibility and shedding shape host competence in amphibians. Functional Ecology, 40(3), 585–598.

Diekmann, O., Heesterbeek, J. A. P., & Metz, J. A. J. (1990). On the definition and the computation of the basic reproduction ratio R0 in models for infectious diseases in heterogeneous populations. Journal of Mathematical Biology, 28, 365–382.

Diekmann, O., Heesterbeek, J. A. P., & Roberts, M. G. (2010). The construction of next-generation matrices for compartmental epidemic models. Journal of the Royal Society Interface, 7, 873–885.

Downs, C. J., Schoenle, L. A., Han, B. A., Harrison, J. F., & Martin, L. B. (2019). Scaling of host competence. Trends in parasitology, 35(3), 182–192.

Duggal, N. K., Bosco-Lauth, A., Bowen, R. A., Wheeler, S. S., Reisen, W. K., Felix, T. A., Mann, B. R., Romo, H., Swetnam, D. M., Barrett, A. D. T., & Brault, A. C. (2014). Evidence for co-evolution of West Nile virus and house sparrows in North America. PLoS Neglected Tropical Diseases, 8, e3262.

Dwyer, G., Elkinton, J. S., & Buonaccorsi, J. P. (1997). Host heterogeneity in susceptibility and disease dynamics: tests of a mathematical model. American Naturalist, 150, 685–707.

Dye, C., & Hasibeder, G. (1986). Population dynamics of mosquito-borne disease: effects of flies which bite some people more frequently than others. Transactions of the Royal Society of Tropical Medicine and Hygiene, 80, 69–77.

Fenton, A., Streicker, D. G., Petchey, O. L., & Pedersen, A. B. (2015). Are all hosts created equal? Partitioning host species contributions to parasite persistence in multihost communities. The American Naturalist, 186(5), 610–622.

Fraser, C., Hollingsworth, T. D., Chapman, R., de Wolf, F., & Hanage, W. P. (2007). Variation in HIV-1 set-point viral load: epidemiological analysis and an evolutionary hypothesis. Proceedings of the National Academy of Sciences, 104, 17441–17446.

Garrett-Jones, C. (1964). Prognosis for interruption of malaria transmission through assessment of the mosquito’s vectorial capacity. Nature, 204, 1173–1175.

Gervasi, S. S., Civitello, D. J., Kilvitis, H. J., & Martin, L. B. (2015). The context of host competence: a role for plasticity in host–parasite dynamics. Trends in parasitology, 31(9), 419–425.

Gervasi, S. S., Burkett-Cadena, N., Burgan, S. C., Schrey, A. W., Hassan, H. K., Unnasch, T. R., & Martin, L. B. (2016). Host stress hormones alter vector feeding preferences, success, and productivity. Proceedings of the Royal Society B, 283(1836), 20161278.

Gervasi, S. S., Burgan, S. C., Hofmeister, E., Unnasch, T. R., & Martin, L. B. (2017a). Stress hormones predict a host superspreader phenotype in the West Nile virus system. Proceedings of the Royal Society B, 284(1859), 20171090.

Gervasi, Stephanie S.; Burgan, Sarah C.; Hofmeister, Erik et al. (2017b). Data from: Stress hormones predict a host superspreader phenotype in the West Nile virus system [Dataset]. Dryad.

Greenspan, S. E., Longcore, J. E., & Calhoun, A. J. (2012). Host invasion by Batrachochytrium dendrobatidis: fungal and epidermal ultrastructure in model anurans. Diseases of Aquatic Organisms, 100, 201–210.

Gubbins, S. (2024). Quantifying the relationship between within-host dynamics and transmission for viral diseases of livestock. Journal of the Royal Society Interface, 21(211), 20230445.

Hamer, G. L., Kitron, U. D., Goldberg, T. L., Brawn, J. D., Loss, S. R., Ruiz, M. O., Hayes, D. B., & Walker, E. D. (2009). Host selection by *Culex pipiens* mosquitoes and West Nile virus amplification. American Journal of Tropical Medicine and Hygiene, 80, 268–278.

Hamer, S. A., Hickling, G. J., Sidge, J. L., Walker, E. D., & Tsao, J. I. (2012). Synchronous phenology of juvenile *Ixodes scapularis*, vertebrate host relationships, and associated patterns of *Borrelia burgdorferi* ribotypes in the midwestern United States. Ticks and Tick-borne Diseases, 3(2), 65–74.

Hardy, J. L., Houk, E. J., Kramer, L. D., & Reeves, W. C. (1983). Intrinsic factors affecting vector competence of mosquitoes for arboviruses. Annual Review of Entomology, 28, 229–262.

Harris, T., Geard, N., & Zachreson, C. (2023). Correlation of viral loads in disease transmission could affect early estimates of the reproduction number. Journal of the Royal Society Interface, 20(202), 20220827.

Hartemink, N. A., Randolph, S. E., Davis, S. A., & Heesterbeek, J. A. P. (2008). The basic reproduction number for complex disease systems: defining R0 for tick-borne infections. The American Naturalist, 171, 743–754.

Hawley, D. M., & Altizer, S. M. (2011). Disease ecology meets ecological immunology: understanding the links between organismal immunity and infection dynamics in natural populations. Functional Ecology, 25(1), 48–60.

Hawley, D. M., Etienne, R. S., Ezenwa, V. O., & Jolles, A. E. (2011). Does animal behavior underlie covariation between hosts’ exposure to infectious agents and susceptibility to infection? Implications for disease dynamics. Integrative and Comparative Biology, 51(4), 528–539.

Haydon, D. T., Cleaveland, S., Taylor, L. H., & Laurenson, M. K. (2002). Identifying reservoirs of infection: a conceptual and practical challenge. Emerging Infectious Diseases, 8(12), 1468–1473.

Hénaux, V., & Samuel, M. D. (2011). Avian influenza shedding patterns in waterfowl: implications for surveillance, environmental transmission, and disease spread. Journal of Wildlife Diseases, 47(3), 566–578.

Herrera, J., & Nunn, C. L. (2019). Behavioural ecology and infectious disease: implications for conservation of biodiversity. Philosophical Transactions of the Royal Society B, 374(1781), 20180054.

Hofmeester, T. R., Coipan, E. C., Van Wieren, S. E., Prins, H. H. T., Takken, W., & Sprong, H. (2016). Few vertebrate species dominate the Borrelia burgdorferi sl life cycle. Environmental Research Letters, 11(4), 043001.

Huang, C.-I., Kay, S. C., Davis, S., Tufts, D. M., Gaffett, K., Tefft, B., & Diuk-Wasser, M. A. (2019). High burdens of *Ixodes scapularis* larval ticks on white-tailed deer may limit Lyme disease risk in a low biodiversity setting. Ticks and Tick-borne Diseases, 10, 258–268.

Huang, Z. Y. X., Halliday, F. W., & Becker, D. J. (2023). Host functional traits as the nexus for multilevel infection patterns. Trends in Ecology & Evolution, 38(12), 1125–1128.

Johnson, P. T. J., Stewart Merrill, T. E., Dean, A. D., & Fenton, A. (2024). Diverging effects of host density and richness across biological scales drive diversity–disease outcomes. Nature Communications, 15(1), 1937.

Jourdain, E., Gunnarsson, G., Wahlgren, J., Latorre-Margalef, N., Bröjer, C., Sahlin, S., Svensson, L., Waldenström, J., Lundkvist, Å., & Olsen, B. (2010). Influenza virus in a natural host, the mallard: experimental infection data. PLoS ONE, 5(1), e8935.

Ke, R., Zitzmann, C., Ho, D. D., Ribeiro, R. M., & Perelson, A. S. (2021). In vivo kinetics of SARS-CoV-2 infection and its relationship with a person’s infectiousness. Proceedings of the National Academy of Sciences, 118, e2111477118.

Keesing, F., Belden, L. K., Daszak, P., Dobson, A., Harvell, C. D., Holt, R. D., … & Ostfeld, R. S. (2010). Impacts of biodiversity on the emergence and transmission of infectious diseases. Nature, 468(7324), 647–652.

Keesing, F., Brunner, J., Duerr, S., Killilea, M., LoGiudice, K., Schmidt, K., … & Ostfeld, R. S. (2009). Hosts as ecological traps for the vector of Lyme disease. Proceedings of the Royal Society B: Biological Sciences, 276(1675), 3911–3919.

Kilpatrick, A. M., Kramer, L. D., Campbell, S. R., Alleyne, E. O., Dobson, A. P., & Daszak, P. (2005). West Nile virus risk assessment and the bridge vector paradigm. Emerging infectious diseases, 11(3), 425.

Kilpatrick, A.M., Daszak, P., Jones, M. J., Marra, P. P., & Kramer, L. D. (2006). Host heterogeneity dominates West Nile virus transmission. Proceedings of the Royal Society B: Biological Sciences, 273(1599), 2327–2333.

Klein, S. L., Zink, M. C., & Glass, G. E. (2004). Seoul virus infection increases aggressive behaviour in male Norway rats. Animal Behaviour, 67(3), 421–429.

Knapp, R. A., Joseph, M. B., Smith, T. C., Hegeman, E. E., Vredenburg, V. T., Erdman, J. E., Boiano, D. M., Jani, A. J., & Briggs, C. J. (2022). Effectiveness of antifungal treatments during chytridiomycosis epizootics in populations of an endangered frog. PeerJ, 10, e12712.

Komar, N., Langevin, S., Hinten, S., Nemeth, N., Edwards, E., Hettler, D., … & Bunning, M. (2003). Experimental infection of North American birds with the New York 1999 strain of West Nile virus. Emerging infectious diseases, 9(3), 311–322.

Latorre-Margalef, N., Gunnarsson, G., Munster, V. J., Fouchier, R. A. M., Osterhaus, A. D. M. E., Elmberg, J., Olsen, B., Wallensten, A., Haemig, P. D., Fransson, T., Brudin, L., & Waldenström, J. (2009). Effects of influenza A virus infection on migrating mallard ducks. Proceedings of the Royal Society B, 276(1659), 1029–1036.

Leitch, J. R., Jordan, N. R., Drake, J. M., Cant, M. A., & Alexander, K. A. (2026). Networks of Indirect Contact Promote Spread of an Environmentally Transmitted Pathogen in a Highly Social Species. Ecology and Evolution, 16(6), e73719.

Levi, T., Keesing, F., Holt, R. D., Barfield, M., & Ostfeld, R. S. (2016). Quantifying dilution and amplification in a community of hosts for tick-borne pathogens. Ecological Applications, 26(2), 484–498.

Lindsay, L. R., Barker, I. K., Surgeoner, G. A., McEwen, S. A., Gillespie, T. J., & Addison, E. M. (1998). Survival and development of the different life stages of *Ixodes scapularis* (Acari: Ixodidae) held within four habitats on Long Point, Ontario, Canada. Journal of Medical Entomology, 35(3), 189–199.

Lloyd-Smith, J. O., Schreiber, S. J., Kopp, P. E., & Getz, W. M. (2005). Superspreading and the effect of individual variation on disease emergence. Nature, 438(7066), 355–359.

LoGiudice, K., Ostfeld, R. S., Schmidt, K. A., & Keesing, F. (2003). The ecology of infectious disease: effects of host diversity and community composition on Lyme disease risk. Proceedings of the National academy of Sciences, 100(2), 567–571.

Lunn, T. J., Restif, O., Peel, A. J., Munster, V. J., De Wit, E., Sokolow, S., … & McCallum, H. (2019). Dose–response and transmission: the nexus between reservoir hosts, environment and recipient hosts. Philosophical Transactions of the Royal Society B: Biological Sciences, 374(1782), 20190016.

Marc, A., Kerioui, M., Blanquart, F., Bertrand, J., Mitja, O., Corbacho-Monné, M., … & Guedj, J. (2021). Quantifying the relationship between SARS-CoV-2 viral load and infectiousness. Elife, 10, e69302.

Martin, L. B., Addison, B., Bean, A. G., Buchanan, K. L., Crino, O. L., Eastwood, J. R., … & Grogan, L. F. (2019). Extreme competence: keystone hosts of infections. Trends in Ecology & Evolution, 34(4), 303–314.

Martin, L. B., Burgan, S. C., Adelman, J. S., & Gervasi, S. S. (2016). Host competence: an organismal trait to integrate immunology and epidemiology. Integrative and Comparative Biology, 56(6), 1225–1237.

Martin, L. B., Hanson, H. E., Hauber, M. E., & Ghalambor, C. K. (2021). Genes, environments, and phenotypic plasticity in immunology. Trends in Immunology, 42(3), 198–208.

Mather, T. N., Wilson, M. L., Moore, S. I., Ribeiro, J. M., & Spielman, A. (1989). Comparing the relative potential of rodents as reservoirs of the Lyme disease spirochete (*Borrelia burgdorferi*). American Journal of Epidemiology, 130, 143–150.

Meagher, J., & Friel, N. (2022). Assessing epidemic curves for evidence of superspreading. Journal of the Royal Statistical Society: Series A (Statistics in Society), 185(4), 2179–2202.

Mommer, L., Esser, H. J., van den Booren, S., Adelman, J. S., Bever, J. D., de Boer, W. F., … & Matson, K. D. (2025). Deconstructing host quality offers insight into disease ecology. Trends in Ecology & Evolution, 41 (3), 201–210.

Moran, E. V., Hartig, F., & Bell, D. M. (2016). Intraspecific trait variation across scales: implications for understanding global change responses. Global Change Biology, 22(1), 137–150.

Mordecai, E. A., Caldwell, J. M., Grossman, M. K., Lippi, C. A., Johnson, L. R., Neira, M., … & Villena, O. (2019). Thermal biology of mosquito-borne disease. Ecology Letters, 22(10), 1690–1708.

Nelsen, R. B. (2006). An introduction to copulas (2nd ed.). Springer.

O’Regan, S. M., & Drake, J. M. (2025). Finite mixture models of superspreading in epidemics. Mathematical Biosciences and Engineering, 22(5), 1081–1108.

Ostfeld, R. S., LaDeau, S. L., Oggenfuss, K., Canham, C. D., Fargione, M., Winchcombe, R. J., & Keesing, F. (2026). The ecology of Lyme disease: long-term data, surprises, and a synthesis. Proceedings of the National Academy of Sciences of the United States of America, 123(34), e2610907123.

Paull, S. H., Song, S., McClure, K. M., Sackett, L. C., Kilpatrick, A. M., & Johnson, P. T. (2012). From superspreaders to disease hotspots: linking transmission across hosts and space. Frontiers in Ecology and the Environment, 10(2), 75–82.

Pérez-Umphrey, A. A., Langwig, K. E., Adelman, J. S., Childs, L. M., Garrett-Larsen, J., Hawley, D. M., & Fleming-Davies, A. E. (2026). Infected host competence overshadows heterogeneity in susceptibility in shaping experimental epizootics. Ecology, 107(5), e70407.

Rapti, Z., & Cáceres, C. E. (2016). Effects of intrinsic and extrinsic host mortality on disease spread. Bulletin of mathematical biology, 78(2), 235–253.

Ribeiro, L. P., Carvalho, T., Becker, C. G., Jenkinson, T. S., Leite, D. D. S., James, T. Y., … & Toledo, L. F. (2019). Bullfrog farms release virulent zoospores of the frog-killing fungus into the natural environment. Scientific reports, 9(1), 13422.

Roberts, M. G., & Heesterbeek, J. A. P. (2003). A new method for estimating the effort required to control an infectious disease. Proceedings of the Royal Society B, 270(1522), 1359–1364.

Roberts, M. G., & Heesterbeek, J. A. P. (2013). Characterizing the next-generation matrix and basic reproduction number in ecological epidemiology. Journal of mathematical biology, 66(4), 1045–1064.

Ruden, R. M., & Adelman, J. S. (2021). Disease tolerance alters host competence in a wild songbird. Biology Letters, 17(10), 20210362.

Ruel, J. J., & Ayres, M. P. (1999). Jensen’s inequality predicts effects of environmental variation. Trends in Ecology & Evolution, 14(9), 361–366.

Saltelli, A., Annoni, P., Azzini, I., Campolongo, F., Ratto, M., & Tarantola, S. (2010). Variance based sensitivity analysis of model output. Design and estimator for the total sensitivity index. Computer Physics Communications, 181, 259–270.

Sih, A., Spiegel, O., Godfrey, S., Leu, S., & Bull, C. M. (2018). Integrating social networks, animal personalities, movement ecology and parasites: a framework with examples from a lizard. Animal behaviour, 136, 195–205.

Smith, D. L., Battle, K. E., Hay, S. I., Barker, C. M., Scott, T. W., & McKenzie, F. E. (2012). Ross, Macdonald, and a theory for the dynamics and control of mosquito-transmitted pathogens. PLoS pathogens, 8(4), e1002588.

Spielman, A. (1999). The Role of Surveillance in Interventions Directed Against Vector-Borne Disease. Ecosystem Health, 5(3), 141–145.

Stewart Merrill, T. E., & Johnson, P. T. (2020). Towards a mechanistic understanding of competence: a missing link in diversity–disease research. Parasitology, 147(11), 1159–1170.

Stewart Merrill, T. E., Calhoun, D. M., & Johnson, P. T. (2022). Beyond single host, single parasite interactions: Quantifying competence for complete multi-host, multi-parasite communities. Functional Ecology, 36(8), 1845–1857.

Streicker, D. G., Fenton, A., & Pedersen, A. B. (2013). Differential sources of host species heterogeneity influence the transmission and control of multihost parasites. Ecology letters, 16(8), 975–984.

Tarasiuk, K., Kycko, A., Świętoń, E., Bocian, Ł., Wyrostek, K., & Śmietanka, K. (2023). Homo- and heterosubtypic immunity to low pathogenic avian influenza virus mitigates the clinical outcome of infection with highly pathogenic avian influenza H5N8 clade 2.3.4.4.b in captive mallards (Anas platyrhynchos). Pathogens, 12(2), 217.

Taube, J. C., Miller, P. B., & Drake, J. M. (2022). An open-access database of infectious disease transmission trees to explore superspreader epidemiology. PLoS Biology, 20(6), e3001685.

Tufts, D. M., McClure, M., & Diuk-Wasser, M. A. (2021). *Ixodes scapularis* (Acari: Ixodidae) nymphal survival and host-finding success in the eastern United States. Journal of Medical Entomology, 58, 929–938.

van den Driessche, P. (2017). Reproduction numbers of infectious disease models. Infectious Disease Modelling, 2, 288–303.

VanderWaal, K. L., & Ezenwa, V. O. (2016). Heterogeneity in pathogen transmission. Functional Ecology, 30(10), 1606–1622.

Vazquez-Prokopec, G. M., Perkins, T. A., Waller, L. A., Lloyd, A. L., Reiner, R. C., Scott, T. W., & Kitron, U. (2016). Coupled heterogeneities and their impact on parasite transmission and control. Trends in Parasitology, 32(5), 356–367.

Violle, C., Enquist, B. J., McGill, B. J., Jiang, L., Albert, C. H., Hulshof, C., Jung, V., & Messier, J. (2012). The return of the variance: intraspecific variability in community ecology. Trends in Ecology & Evolution, 27(4), 244–252.

Vredenburg, V. T., Knapp, R. A., Tunstall, T. S., & Briggs, C. J. (2010). Dynamics of an emerging disease drive large-scale amphibian population extinctions. Proceedings of the National Academy of Sciences, 107(21), 9689–9694.

Woolhouse, M. E. J., Dye, C., Etard, J.-F., Smith, T., Charlwood, J. D., Garnett, G. P., et al. (1997). Heterogeneities in the transmission of infectious agents: implications for the design of control programs. Proceedings of the National Academy of Sciences, 94, 338–342.

