## Appendix S1 for "Formalizing host competence"

**Appendix** **S1.** **Host competence, potential and capacity: a derivation from the next-generation operator**

*Supporting Information for Gibson et al.,* *Formalizing* *host* *competence*

### 1. Purpose and result

Host competence has usually been written as a product of biologically sensible terms and justified by interpretation. Here we derive it instead, so that each term of the index used in the main text appears from a definition rather than an assertion. The definition we adopt is a natural one: the contribution an individual host makes to the next generation of infections. This is the quantity from which the basic reproduction number is built (Diekmann et al. 1990, 2010; van den Driessche 2017), and it is the host-side analogue of the way vectorial capacity is derived for mosquitoes (Garrett-Jones 1964; Smith et al. 2012).

The derivation differs from a conventional next-generation treatment in one structural respect, and that difference produces the framework used in the main text. The rate at which an infected host makes transmission-relevant contacts is split into two parts: an ecological rate, θ, set by the density of vectors, conspecifics or environmental pathogen the host encounters, and a dimensionless modifier, *b*(τ), by which infection changes that rate for this individual. Only the second is a property of the host, so only the second stays inside the host-level trait. What remains is

*a* = ∫_0_^∞^ *b*(τ) φ(*L*(τ)) Σ(τ) dτ, *H*_comp_ = ε(*D*) · *a*	(S1)

an infection-age integral of the host’s per-contact infectiousness, weighted by the probability that it is still alive and infected and by its own infection-induced behavior, multiplied by the probability that exposure produced infection in the first place. The consequence of this choice is that *H*_comp_ carries units of time rather than being dimensionless, so it is not itself a reproduction number and the value *H*_comp_ = 1 has no meaning (Section 5.1). The dimensionless quantities, and with them the thresholds, reappear as soon as the ecological terms are restored:

*H*_pot_ = θ · ε(*D*) · *a*, *H*_cap,i_ = *n*_i_ · θ_i_ · ε_i_(*D*) · *a*_i_, *R*_i_ = θ · *a*_i_, *T*_i_ = θ · *a*_i_ · *V*	(S2)

Sections 2–4 derive these expressions, Section 5 establishes the thresholds and shows why contact enters once rather than squared, Section 6 states the three assumptions under which Eq. S1 reduces to the equation simulated in the main text, Section 7 treats the upper limit of the integral, and Section 8 recovers three established species-level indices as special cases. Section 9 derives two analytic results that explain the paper’s main empirical patterns: why variation in competence rises as mean competence falls, and why covariation between load and contact (*b*) matters only for low-competence species. Section 10 sets out scope, identifiability and limitations.

### 2. Notation and biological setup

Consider a focal host individual *i* and let τ ≥ 0 denote infection age, the time since that individual became infected. Its within-host state is summarized by a pathogen-load trajectory *L*_i_(τ) ≥ 0. We use the following individual-level quantities.

| **Symbol** | **Meaning** | **Units** |
| --- | --- | --- |
| ε_i_(D) | probability the host becomes infected when exposed to dose D (susceptibility) | dimensionless |
| L_i_(τ) | within-host pathogen load at infection age τ | load |
| φ(L) | probability that a transmission-relevant contact made at load L infects the recipient (infectiousness per contact) | dimensionless |
| b_i_(τ) | rate of transmission-relevant contacts at infection age τ relative to the same individual’s uninfected baseline (behavior) | dimensionless |
| μ_i_(τ) | mortality hazard at infection age τ | time^−1^ |
| γ_i_(τ) | clearance (recovery) hazard at infection age τ | time^−1^ |
| Σ_i_(τ) | probability the host is still alive and still infected at infection age τ | dimensionless |
| θ_i_ | rate of transmission-relevant contacts with susceptible recipients, uninfected baseline (transmissibility) | time^−1^ |
| n_i_ | abundance or density of hosts of type i | hosts (or hosts area^−1^) |
| V | expected host infections produced per infected vector, or per unit of environmental pathogen | dimensionless |

The survival-of-infectiousness function is

Σ_i_(τ) = exp( −∫_0_^τ^ [ μ_i_(u) + γ_i_(u) ] du ), (S3)

so the host remains an active source only while it is both alive (μ) and not yet recovered (γ). Setting γ ≡ 0 recovers pure survival weighting (see Section 7).

Three pairs of quantities are kept definitionally disjoint throughout, and most of the structure of the framework follows from doing so.

- φ(*L*) versus *b*(τ). Infectiousness is within-host: given that a contact occurs, how likely is transmission at the current load. Behavior is between-host: how often contacts occur. Empirically φ is saturating in load (Fraser et al. 2007; Ke et al. 2021; Marc et al. 2021). Keeping quantities separate is what prevents the effect of load from being counted twice once behavior is allowed to depend on load.
- *b*(τ) versus θ. Both describe contact, but *b* is the change that infection causes in a focal individual host and θ is the baseline that change is measured against. The first is an organismal trait; the second is a property of the context.
- ε versus φ. Both are infection probabilities, but ε governs whether the focal host became infected and φ governs whether a recipient does. The focal host’s own ε therefore never appears in its onward transmission (Section 5.2).

### 3. Background: the next-generation operator

For a structured infectious population, the basic reproduction number is the spectral radius of the next-generation operator *K*, whose kernel *k*(ξ, η) is the expected number of new infections of state ξ produced by one newly infected individual of state η over its entire infection (Diekmann et al. 1990, 2010):

*R*_0_ = ρ(*K*), (*K*ψ)(ξ) = ∫ *k*(ξ, η) ψ(η) dη. (S4)

For compartmental models this reduces to *K* = *FV*^−1^ (Diekmann et al. 2010), and for ecological multi-host communities it becomes a next-generation matrix whose element *K*_ji_ is the expected number of type-*j* infections caused by a type-*i* individual (Roberts & Heesterbeek 2013; Hartemink et al. 2008). Hartemink et al. (2008) developed exactly this matrix for tick-borne systems, with each element carrying an explicit biological meaning, which is the closest existing precedent for treating host competence as a next-generation quantity.

Two issues are important here. First, an individual’s contribution to local infections is its column sum in *K*: the total expected secondary infections caused by an individual of type *i* is Σ_j_ *K*_ji_, a per-individual reproduction number rather than a population average. Second, because the elements are expected counts of secondary cases, the natural per-individual threshold at which an individual replaces itself (not necessarily the invasion threshold, which is ρ(*K*); Section 4.6) is Σ_j_ *K*_ji_ = 1. Section 4 describes that column sum explicitly when the relevant state is the within-host load trajectory, then parses its host-level and ecology-level parts.

### 4. Derivation

#### 4.1 The infection-age renewal kernel

Fix the focal individual *i*, infected at τ = 0 into an otherwise fully susceptible environment: the invasion regime in which *R*_0_ and the next-generation operator are defined. Define its instantaneous infectivity *A*_i_(τ) as the expected rate at which the host generates new infections at infection age τ. Decomposing that rate into still-active × opportunity × success,

*A*_i_(τ) = Σ_i_(τ) × *c*_i_(τ) × φ(*L*_i_(τ)), (S5)

where *c*_i_(τ) is the rate at which the individual makes transmission-relevant contacts with susceptible recipients at infection age τ. Integrating over the whole infection gives the expected number of secondary infections produced by this source,

ν_i_ = ∫_0_^∞^ *A*_i_(τ) dτ = ∫_0_^∞^ *c*_i_(τ) φ(*L*_i_(τ)) Σ_i_(τ) dτ. (S6)

This ν_i_ is the individual reproduction number of Lloyd-Smith et al. (2005): the mean of the offspring distribution for individual *i*, with *R*_0_ equal to the weighted mean of ν_i_ over newly infected hosts.

#### 4.2 Factoring contact into an ecological rate and a host modifier

The step that defines the present framework is the treatment of *c*_i_(τ). Write

*c*_i_(τ) = θ_i_ · *b*_i_(τ), with *b*_i_(0^−^) = 1, (S7)

where θ_i_ is the individual’s contact rate immediately before infection and *b*_i_(τ) is the factor by which infection changes it. As written, Eq. S7 is a reparameterization rather than an assumption: any positive contact trajectory can be split this way, and the normalization *b*(0^−^) = 1 fixes the split uniquely. Its content lies in what is then attributed to whom. We treat θ_i_ as a property of the ecological setting, because it is determined by the density of vectors that bite the host, larvae that attach to it, conspecifics it meets or substrate into which it sheds, etc., and we treat *b*_i_(τ) as a property of the host, because it is determined by sickness behaviors, grooming, anti-vector defense and morbidity. Substituting Eq. S7 into Eq. S6 and taking θ_i_ outside the integral, which requires that the baseline rate be constant over the infectious period (Section 10),

ν_i_ = θ_i_ ∫_0_^∞^ *b*_i_(τ) φ(*L*_i_(τ)) Σ_i_(τ) dτ ≡ θ_i_ · *a*_i_. (S8)

#### 4.3 Infectiousness

The integral in Eq. S8 is the host-side factor of transmissibility, which we call infectiousness:

*a*_i_ = ∫_0_^∞^ *b*_i_(τ) φ(*L*_i_(τ)) Σ_i_(τ) dτ. (S9)

Every quantity inside it can in principle be measured on an animal in isolation. Because the integrand is dimensionless and dτ has units of time, *a* has units of time; it is the number of fully infectious, fully active days the individual supplies, which we call effective infectious days. It is conditional on infection, which matches the experimental concept of competence in work where animals were deliberately infected and their infectiousness to vectors or conspecifics measured (Komar et al. 2003; Brunner et al. 2008). Because *b* and Σ may both depend on load, Eq. S9 represents sickness behavior, but also parasite tolerance and parasite virulence within a single integral.

#### 4.4 Susceptibility and competence

Equation S9 is conditional on the focal individual being infected. Competence, as conventionally defined, includes the ability to acquire infection as well as to transmit it. Multiplying by the probability of acquisition given exposure to dose *D* yields

*H*_comp,i_ = ε_i_(*D*) · *a*_i_, (S10)

the expected number of effective infectious days generated per exposure. Exposure itself, meaning the encounter rate that delivers dose *D* to the focal host, is deliberately outside this expression: competence is defined conditional on exposure, with ε carrying the dose dependence. Heterogeneity in encounter on the acquisition side is a population-level property, separated by construction from the organismal trait. Behavior that changes the dose an individual receives belongs to ε, while behavior that changes contact after infection belongs to *b*, so no behavioral effect is omitted or double-counted.

#### 4.5 Potential and capacity

Restoring the ecological rate to Eq. S10 gives host potential, the expected number of new infections arising from one exposure of the focal host,

*H*_pot,i_ = θ_i_ · ε_i_(*D*) · *a*_i_, (S11)

which is dimensionless because θ has units of inverse time and *a* units of time. Weighting potential by the abundance of hosts of that type gives host capacity, the contribution of a population,

*H*_cap,i_ = *n*_i_ · θ_i_ · ε_i_(*D*) · *a*_i_. (S12)

The progression from infectiousness and susceptibility to potential to capacity mirrors the progression from vector competence to vector potential (Bauer et al. 2026) to vectorial capacity (Garrett-Jones 1964; Smith et al. 2012) and is the reason the four terms can be measured and reported separately.

Notably, the availability of susceptible recipients, and any fixed per-contact scaling, are properties of the system rather than of the focal host. Carrying them explicitly inside θ makes *R*_i_ = θ_i_ · *a*_i_ a reproduction number, comparable across systems, with a meaningful threshold at 1. Absorbing them into an unmeasured θ instead leaves *H*_pot_ defined only up to a system-specific multiplicative constant, licensing within-system rankings alone. The simulations in the main text take the second route for every system except Lyme disease, where larval tick burdens supply θ directly, which is why magnitudes are not compared across systems there.

**4.6 From individual offspring numbers to R₀**

Individuals differ in the rate *x*_i_ at which they are exposed, in susceptibility ε_i_ and in abundance *n*_i_. If new infections fall on type *j* in proportion to *w*_j_ = *n*_j_ *x*_j_ ε_j_ / Σ_k_ *n*_k_ *x*_k_ ε_k_ (random mixing by attractiveness), then *K*_ji_ = ν_i_ *w*_j_, *K* has rank one, and R₀ = ρ(*K*) = Σ_i_ *w*_i_ ν_i_. (S13)

R₀ is therefore the mean of ν_i_ over newly infected individuals, weighted by exposure and susceptibility. It equals the unweighted mean E[ν_i_] only when infection is independent of ν_i_. When contact varies and exposure-side and transmission-side contact are identical (θ_i_ = *x*_i_), R₀ = E[θ] E[a] (1 + CV_θ_^2^) if a is independent of θ and ε is constant, so for CV = 1 R₀ is twice E[ν_i_].

### 5. Thresholds and reproduction numbers

#### 5.1 Why competence has units of time and no threshold

In a conventional treatment the individual-level quantity is ν_i_, which is dimensionless because it counts infections, so the threshold ν_i_ = 1 follows immediately. Here the dimensionless quantity has been deliberately split, and the host keeps only the part that does not contain θ. Here, ε, φ, *b* and Σ are dimensionless, dτ has units of time, so *a* and *H*_comp_ have units of time, and θ, with units of inverse time, is exactly what is needed to generate a dimensionless value. Consequently, *H*_comp_ is not a reproduction number, the value *H*_comp_ = 1 carries no information, and competence alone cannot say whether an individual amplifies or dampens transmission. That is not a defect of the index but the mathematical consequence of requiring that competence be a property of the host alone. Whether an individual replaces itself is a joint question about the host and its setting.

#### 5.2 Directly transmitted pathogens

From Eq. S8, the expected number of new infections caused by an individual that is already infected is

*R*_i_ = θ · *a*_i_, (S14)

with *R*_i_ = 1 separating host individuals that more than replace themselves from those that do not, and R₀ = Σ_i_ *w*_i_*R*_i_ (Eq. S13), which equals E[*R*_i_] when infection is independent of *R*_i_. The focal host’s own susceptibility does not appear: ε governs whether this individual became infected, whereas the susceptibility of recipients is already carried by φ, which is the probability that a contact made at the current load infects the recipient. Including ε in *R*_i_ would count the focal host’s acquisition of a pathogen twice, once when it became infected and again in its onward transmission.

#### 5.3 Vector-borne and environmentally transmitted pathogens

For a vector-borne pathogen the next-generation matrix has the two-type form

*K* = [ 0 *K*_hv_ ; *K*_vh_ 0 ], *K*_vh_ = θ · *a*, *K*_hv_ = *V*, (S15)

where *K*_vh_ is the expected number of vectors infected by one infected host and *K*_hv_ the expected number of hosts infected by one infected vector. Here θ is the rate at which the focal host is bitten, φ is the probability that a biting vector becomes infected at the current host load, and *V* folds in vector survival through the extrinsic incubation period, vector competence and subsequent biting. The spectral radius of a matrix with this form is the geometric mean of the two half-cycles, *R*_0_ = √(θ *a V*), which is exactly the structure of vectorial capacity. Equivalently, and more usefully for a host-centered index, the two-step number of new infections from an individual host is

*T*_i_ = θ · *a*_i_ · *V*, (S16)

Which, for identical hosts, is the type-reproduction number *T*, with *T* = 1 exactly equivalent to *R*_0_ = 1 (Roberts & Heesterbeek 2003; Hartemink et al. 2008). Host competence is therefore the host’s share of one half of the cycle (Eq. S10), which is the appropriate host-side trait; the environmental case is identical with *V* reinterpreted as the expected host infections per unit of pathogen deposited.

#### 5.4 Why contact enters once rather than squared as in vectorial capacity

Vectorial capacity contains the squared biting rate because one short-lived vector bites once to acquire the pathogen and again to transmit it, at the same rate and on the same kind of host; the same rate therefore appears in both halves of the cycle and the square is unavoidable. The host-to-vector half cycle is probably too complex for this assumption. Accordingly, in Eq. S15, the focal host’s contact rate appears in *K*_vh_, the half-cycle that runs from this host to vectors. The rate at which the host was itself exposed appears in *K*_hv_, the half-cycle that runs from vectors to hosts, where it is a property of the recipient rather than of the focal individual. In the column of *K* belonging to an infected host, contact therefore appears exactly once; vector’s biting rate also appears in K_hv_ = V, so the closed cycle T_i_ is proportional to its square, as in vectorial capacity.

That formal statement matches the biology of many systems, meaning that the two contacts are separated in time, sometimes by weeks or months. For instance, *Borrelia burgdorferi* is acquired by *Peromyscus* mainly from nymphs early in summer and transmitted to larvae later in the season. Behaviors occurring in these periods may also involve different vector stages with different host preferences (Brunner & Ostfeld 2008). Further, infection itself could alter the host’s behavior between the two, which is precisely what *b*(τ) represents. Squaring θ would assert that exposure-side and transmission-side contact rates are the same number for the same individual, which for hosts is a strong and unrealistic empirical claim.

#### 5.5 What survives of the squared term: covariance

The squared biting rate for vectorial capacity implicates an important condition for hosts, though, in the form of trait covariance. Let θ_in_ and θ_out_ denote the exposure-side and transmission-side contact rates of the same individual. The probability that an individual becomes infected rises with θ_in_, so infected individuals are a biased sample of the population, over-represented for high θ_in_. If θ_in_ and θ_out_ covary, infected individuals are then also over-represented for high θ_out_, and the mean transmission-side contact rate among infected individuals exceeds the population mean by

E[θ_in_ θ_out_] / ( E[θ_in_] E[θ_out_] ). (S17)

For jointly lognormal contact rates with log-scale correlation ρ and coefficient of variation CV, this inflation factor is exp(ρσ²) with σ² = ln(1 + CV²), rising to 1 + CV² when ρ = 1. The product of mean contact rates is analogous to the squared biting rate of vectorial capacity and S17 is the further factor 1 + CV^2^ in the special case in which the two rates are identical and perfectly correlated, ρ = 1 and CV equal for both. For hosts, the magnitude of the inflation is an empirical question about the persistence of individual differences in contact, not a structural feature of the index.

### 6. The special case simulated in the main text

Equation S9 is general. The simulations use a special case obtained under three explicit assumptions, each stated here with its cost.

**(A1) Behavior does not change with load or infection age**, *b*(τ) ≡ *b*, so *b* leaves the integral. Individual differences in behavior are retained, since *b* varies among individuals; what is lost is any within-individual timing. Formally,

*a* = E[*b*] ∫ φ Σ dτ + ∫ Cov_τ_(*b*, φΣ) dτ, (S18)

so A1 discards the second term, which is the coincidence between when an individual is most infectious and when it is making the fewest or most contacts. The assumption fails conspicuously when sickness behavior is triggered above a load threshold that lies within the infectious range, because *b* then falls exactly where φ is highest; the product *b*(*L*)φ(*L*) becomes hump-shaped rather than saturating, and high-competence species, whose loads climb well past the threshold for transmission, are affected most.

**(A2) Infectiousness is lost only through death**, γ ≡ 0, so Σ(τ) becomes the survival function *S*(τ) and clearance in acute infections is represented by the load trajectory itself returning to zero. The cost is that for infections cleared rather than carried to death, dropping γ would overstate the infectious period if the modelled load did not in fact resolve; Section 7 details the condition under which it does.

**(A3) Infectiousness is a Hill function of load**, φ(*L*) = *L*^n^ / (*K*^n^ + *L*^n^), with half-saturation load *K* and steepness *n*. The Hill function is a phenomenological default rather than a necessary assumption, and φ need not be assumed at all: the true function can be fitted directly to paired load–transmission data. For example, the function can be described from a dose–response model because the donor’s shed load sets the challenge dose delivered to the recipient, which gives specific saturating shapes such as the exponential form φ(*L*) = 1 − e^−rL^ or a Beta–Poisson variant (Brouwer et al. 2017; Lunn et al. 2019). Functions could also be derived from coupled within-host models (Almocera et al. 2018; Della Marca et al. 2023; Gubbins 2024). The results of Section 9 require the sigmoidal Hill form (n>1).

Under A1–A3, Eq. S9 becomes the equation simulated in the main text,

*a* = *b* ∫_0_^∞^ [ *L*(τ)^n^ / (*K*^n^ + *L*(τ)^n^) ] *S*(τ) dτ. (S19)

### 7. What bounds the integral

The upper limit ∞ in Eqs. S9 and S19 is a formal convenience; what matters is that the integrand is driven to zero and the mechanism by which this happens. Three routes arise among the systems we considered (the load, survival and seasonal routes, as explained below), and a fourth bookkeeping device, a clearance hazard, must not be combined with the first.

**The load route.** The Hill map is monotone increasing with φ(0) = 0; as a function of time, it merely tracks the load. For an acute rise-and-fall trajectory with *L*(τ) → 0, the integrand vanishes through the load alone, even with the host fully alive. Clearance is then represented deterministically, through the shape of *L*(τ), and the effective upper limit is the resolution of detectable load.

**The survival route.** For a chronic plateau or a progressive approach to a carrying capacity, *L*(τ) → *L*_∞_ > 0, so φ does not vanish and the integrand is bounded only by *S*(τ) → 0. The effective upper limit is host death, so competence scales with longevity. This dependence can be supplied either by biology or by an artefact: if *S*(τ) carries a realistic mortality schedule, lifespan enters as a genuine, measurable competence trait, but if *S*(τ) is left flat and the integral is simply stopped at a numerical horizon, that arbitrary horizon sets the answer. For a tolerated lifelong infection (e.g., *B. burgdorferi* in *Peromyscus leucopus*), lifespan must be promoted to a first-class competence parameter with an explicit survival schedule.

**The seasonal route.** When transmission-relevant contacts occur only during part of the year, the integral is bounded by the transmission window regardless of load or survival, because θ and hence the opportunity for transmission is zero outside it. The Lyme simulations use a 60-day larval season on this basis. Strictly this amounts to replacing the constant θ of Eq. S7 with a seasonal θ(τ) that is zero outside the window, which is legitimate provided the host’s infection age and the calendar are aligned. Where they are not, the appropriate object is the time-periodic reproduction number (Bacaër & Guernaoui 2006).

**Avoiding double-counting.** The general weight of Eq. S3 admits a fourth route: a clearance hazard γ > 0 that removes the host from the infectious class while it is still alive and while load may still be positive. This is an alternative bookkeeping of the same biology as the load route, not an addition to it. One bounding of the integral must be chosen per individual. Under deterministic clearance, set γ ≡ 0, weight by survival alone and let *L*(τ) → 0 encode resolution; this is correct precisely when each modelled acute trajectory does return below threshold. Under stochastic clearance, set γ > 0, weight by Σ(τ), and let *L*(τ) be the load conditional on still being infected. Applying both a decaying load and a positive clearance hazard counts the same exit twice. At the individual level, parasite clearance is the deterministic event *L*(τ) → 0 or death. The population-level fraction still infected emerges from the distribution of those exit times across the simulated ensemble and should not be re-imposed as a per-individual hazard.

| **Load family (system)** | **L(τ) as τ → ∞** | **Integrand → 0 via** | **Effective upper limit** |
| --- | --- | --- | --- |
| Acute, cleared (WNV, LPAI) | → 0 | φ(L(τ)) → 0 (load) | resolution of load |
| Chronic plateau, tolerated (Lyme in Peromyscus) | → L_∞_ > 0 | θ(τ) → 0 (season) | transmission season |
| Progressive to carrying capacity, tolerated (Bd in bullfrog) | → L_cap_ > 0 | S(τ) → 0 (death) | host lifespan |
| Progressive, lethal (Bd in susceptible frogs) | rises, host dies | S(τ) → 0 (death) | time of death |

### 8. Recovering established species-level indices

Three widely used indices are special cases of Eqs. S10–S12, which makes explicit the assumptions under which each holds.

**Komar et al. (2003).** Their reservoir competence index was the product of susceptibility, mean daily infectiousness and the duration of infectiousness. Setting *b* ≡ 1 and Σ ≡ 1 over an infectious period of length *d*, and writing the mean of φ over that period as φ̄, Eq. S9 gives *a* = φ̄ *d*, so

*H*_comp_ = ε · φ̄ · *d*, (S20)

which is their index exactly. The index is therefore Eq. S10 evaluated for an average individual in daily steps, with no behavioral modifier and no mortality within the infectious period.

**Kilpatrick et al. (2006).** Weighting the Komar index by mosquito feeding preference and host abundance is precisely the step from Eq. S10 to Eqs. S11 and S12: feeding preference is the species-specific part of θ, and abundance is *n*. Their amplification fraction is thus a normalised *H*_cap_.

**Mather et al. (1989) and LoGiudice et al. (2003).** The fraction of infected vectors fed by each host species in a community is

*H*_cap,i_ / Σ_j_ *H*_cap,j_ = *n*_i_ θ_i_ ε_i_ *a*_i_ / Σ_j_ *n*_j_ θ_j_ ε_j_ *a*_j_, (S21)

Which, with exposure equal across species, is the quantity plotted as each species’ share of community capacity in the main text (Fig. 4B). Realized reservoir competence, the probability that a larva feeding on a field-caught host becomes infected, is a different quantity: it folds in natural exposure and prevalence, which Eq. S10 deliberately excludes. It also omits both the number of vectors a host feeds and how long it remains infectious, which Eqs. S11 and S9 supply. Their measures and ours therefore rank species differently whenever burden and infectivity are negatively related, which is the formal reason the eastern gray squirrel ranks low by realized reservoir competence yet produces about 70% as many infected larvae per infected individual as the eastern chipmunk (main text, Section 5).

What is new here is not any of these indices but that each term is defined for individuals, derived from a single expression, and given explicit units, so that the assumptions under which each species-level index holds can be stated rather than assumed.

### 9. Two analytic consequences of the derivation

The derivation yields two results that explain the main empirical patterns reported in the paper, results that would be difficult to anticipate from the index written as a product of terms.

#### 9.1 Averaging over individuals

Because *R*_0_ = E[*R*_i_] = E[θ *a*], when infection is independent of R_i_ and E[θ *a*] = E[θ] E[*a*] + Cov(θ, *a*), individual variation affects the mean number of secondary infections only through the covariance between contact and infectiousness, and, when exposure-side and transmission-side contact are identical, through the variance of contact, which raises R₀ by the factor 1 + CV² (Section 4.6). Variation that is independent of contact leaves the mean untouched and acts instead on the variance of the offspring distribution, which governs the probability of emergence and the degree of superspreading rather than the average (Lloyd-Smith et al. 2005). A second, separate gap arises because *a* is a nonlinear functional of load: by Jensen’s inequality, the mean of individual competences differs from the competence computed at mean traits, with the sign set by whether φ is convex or concave over the range of loads realized. These are distinct arguments for individual data and should not be conflated. The first issue concerns covariance, the second concerns nonlinearity, and in the simulations reported here the second issue proved modest whereas the first was strong. Across the 23 simulated species–pathogen combinations, mean *a* divided by *a* evaluated at mean parameter values was 0.82–1.15 in 20 and at most 1.41 (mourning dove 1.31, white-tailed deer 1.33, laughing gull 1.41), whereas load–contact covariation changed mean potential by ×0.41–2.14 (laughing gull; main text Fig. 2).

#### 9.2 Why variation rises as mean competence falls

The sensitivity of infectiousness to load is governed by the elasticity of the Hill function, which has a particularly simple closed form. Differentiating log φ with respect to log *L*,

d log φ / d log *L* = *n* *K*^n^ / (*K*^n^ + *L*^n^) = *n* [ 1 − φ(*L*) ]. (S22)

The elasticity is therefore *n* when load is far below the half-saturation point and falls to zero as φ approaches one. Two consequences follow directly, and both were observed in the simulations.

First, for small among-individual variation in log load with standard deviation σ_ℓ_, the coefficient of variation of infectiousness contributed by load is approximately *n*[1 − φ̄] σ_ℓ_. Species whose loads sit near the transmission threshold therefore convert load variation into competence variation with a gain of up to *n*, whereas species whose loads saturate infectiousness compress it toward zero. This difference is the mean–variance relationship reported in the main text, derived rather than observed: it is a property of the saturating load–infectiousness map, and of the variances assigned to each species, since the gain multiplies σ_ℓ_.

Second, the same elasticity controls how much covariation between load and contact can matter. As Cov(θ, *a*) = ρ σ_θ_ σ_a_ and σ_a_ scales with *n*[1 − φ̄], the effect of load–contact covariation on mean potential is proportional to *n*[1 − φ̄] and vanishes as φ̄ → 1. Covariation reshapes the upper tail of potential in low-competence species but does nothing at all in chytridiomycosis because fungal loads lie far above the threshold for infectiousness.

Evaluating Eq. S22 at each focal species’ central load tracks the simulated ordering only roughly.

| **Species** | **Mean φ at central load** | **Elasticity**  **n(1 − mean φ)** | **Simulated mean potential at ρ = +0.7 (relative to ρ = 0)** |
| --- | --- | --- | --- |
| Mallard | 0.96 | 0.08 | 1.17 |
| Laughing gull | 0.08 | 1.83 | 2.14 |
| Blue jay † | 0.81 | 0.86 | 1.14 |
| Mourning dove † | 0.14 | 3.84 | 1.67 |
| American bullfrog | 1.00 | 0.00 | 0.97 |
| Mountain yellow-legged frog | 1.00 | 0.00 | 1.00 |
| White-footed mouse | 0.92 | 0.12 | 1.05 |
| Eastern gray squirrel | 0.26 | 1.11 | 1.58 |

† For West Nile virus the Hill function is applied to log_10_ viremia rather than to load directly, so the elasticity is with respect to log titer and is not on the same scale as the other systems; the ordering within the system is unaffected, but the WNV rows cannot be ranked against the other rows.

Equation S22 is a local, pointwise approximation and captures only the amplitude channel, that is, how a change in load alters infectiousness at a given moment. It does not capture the duration channel, by which raising peak load also lengthens the time spent above the transmission threshold. Where the elasticity is near zero the amplitude channel contributes almost nothing and residual variation must come from duration, survival or behavior, which is exactly what the variance decomposition in Appendix S3 finds: lifespan dominates in the bullfrog, behavior in the mountain yellow-legged frog, and load timing in the mallard.

### 10. Scope, identifiability and limitations

- **Invasion regime.** Like *R*_0_ itself, Eqs. S8 and S14–S16 are defined in a fully or nearly susceptible environment. They quantify competence for invasion and early spread, not for an established endemic equilibrium where depletion of susceptibles matters.
- **Constancy of the baseline contact rate.** Taking θ_i_ outside the integral in Eq. S8 requires the uninfected baseline rate to be constant over the infectious period. For acute infections lasting days this condition is innocuous. For infections lasting months to years, it is not, and seasonal variation in vector activity is the obvious violation. A seasonal adjustment (i.e., Section 7) is the simplest way to accommodate this issue, and the time-periodic reproduction number (Bacaër & Guernaoui 2006) is the general one.
- **Transmissibility remains partly a community property.** Once the availability of susceptible recipients is explicit, potential and capacity are context-dependent through θ, consistent with the finding that reservoir status is not a fixed species trait but depends on context (Roberts & Heesterbeek 2013, 2020). The organismal-trait framing holds cleanly for ε, φ, *b* and Σ; θ is where context re-enters, by design.
- **Identifiability scales with generality.** A load-dependent *b* and a time-varying θ require data that are rarely available. The special case of Eq. S19 is what is estimable in most systems today; the general form is the scaffold.

*Susceptibility:* A particular identifiability problem deserves particular attention because it constrains what can be measured rather than what can be modelled. Susceptibility is defined as a dose–response function, ε(*D*), but a single individual exposed once yields a single binary outcome, and repeated exposure is not a remedy because the first exposure generates immune memory. An individual’s dose–response curve is therefore not identifiable from one host organism, no matter how well the experiment is designed. Four routes around this problem are available, each estimating something slightly different:

i) simultaneous independent exposures within one individual, where anatomy permits them, as with multiple skin sites for *Batrachochytrium dendrobatidis* or multiple attached nymphs for *B. burgdorferi*;
ii) dose–response measured ex vivo on cells or tissue from each individual and calibrated against a single in vivo challenge;
iii) a graded early-establishment measure, such as pathogen load in the first days after a single exposure, in place of a binary infected-or-not outcome; and
iv) hierarchical estimation of the distribution of ε across many individuals at several doses, which identifies its variance even though no individual value is identified, since heterogeneity in susceptibility flattens the population dose–response curve on log dose (Dwyer et al. 1997; Langwig et al. 2017).

Estimating the covariance between ε and *a* is harder still, because *a* is observable only in individuals that became infected, and those individuals are exactly the ones with high ε. The sample available for estimating the covariance is thus truncated by construction. A pre-exposure predictor of ε, measured on every individual before the outcome is known, is the only route we can see that avoids this truncation, and it is an argument for developing assay-based or biomarker-based proxies for susceptibility rather than relying on challenge outcomes alone.

### References

Almocera, A.E.S., Nguyen, V.K., Hernandez-Vargas, E.A. (2018). Multiscale model within-host and between-host for viral infectious diseases. Journal of Mathematical Biology 77, 1035–1057.

Bacaër, N., Guernaoui, S. (2006). The epidemic threshold of vector-borne diseases with seasonality: the case of cutaneous leishmaniasis in Chichaoua, Morocco. Journal of Mathematical Biology 53, 421–436.

Bauer, A.M., Burkett-Cadena, N.D., Reeves, L.E., Alto, B.W., Campbell, L.P. (2026). Vector potential index: bridging competence and contribution as an integrative measure of relative transmission capability. Ecology and Evolution 16, e72705.

Brouwer, A.F., Weir, M.H., Eisenberg, M.C., Meza, R., Eisenberg, J.N.S. (2017). Dose-response relationships for environmentally mediated infectious disease transmission models. PLoS Computational Biology 13, e1005481.

Brunner, J.L., LoGiudice, K., Ostfeld, R.S. (2008). Estimating reservoir competence of Borrelia burgdorferi hosts: prevalence and infectivity, sensitivity, and specificity. Journal of Medical Entomology 45, 139–147.

Brunner, J.L., Ostfeld, R.S. (2008). Multiple causes of variable tick burdens on small-mammal hosts. Ecology 89, 2259–2272.

Della Marca, R., Loy, N., Tosin, A. (2023). An SIR model with viral load-dependent transmission. Journal of Mathematical Biology 86, 61.

Diekmann, O., Heesterbeek, J.A.P., Metz, J.A.J. (1990). On the definition and the computation of the basic reproduction ratio R0 in models for infectious diseases in heterogeneous populations. Journal of Mathematical Biology 28, 365–382.

Diekmann, O., Heesterbeek, J.A.P., Roberts, M.G. (2010). The construction of next-generation matrices for compartmental epidemic models. Journal of the Royal Society Interface 7, 873–885.

Dwyer, G., Elkinton, J.S., Buonaccorsi, J.P. (1997). Host heterogeneity in susceptibility and disease dynamics: tests of a mathematical model. American Naturalist 150, 685–707.

Dye, C., Hasibeder, G. (1986). Population dynamics of mosquito-borne disease: effects of flies which bite some people more frequently than others. Transactions of the Royal Society of Tropical Medicine and Hygiene 80, 69–77.

Fraser, C., Hollingsworth, T.D., Chapman, R., de Wolf, F., Hanage, W.P. (2007). Variation in HIV-1 set-point viral load: epidemiological analysis and an evolutionary hypothesis. PNAS 104, 17441–17446.

Garrett-Jones, C. (1964). Prognosis for interruption of malaria transmission through assessment of the mosquito’s vectorial capacity. Nature 204, 1173–1175.

Gubbins, S. (2024). Quantifying the relationship between within-host dynamics and transmission for viral diseases of livestock. Journal of the Royal Society Interface 21, 20230445.

Hartemink, N.A., Randolph, S.E., Davis, S.A., Heesterbeek, J.A.P. (2008). The basic reproduction number for complex disease systems: defining R0 for tick-borne infections. American Naturalist 171, 743–754.

Ke, R., Zitzmann, C., Ho, D.D., Ribeiro, R.M., Perelson, A.S. (2021). In vivo kinetics of SARS-CoV-2 infection and its relationship with a person’s infectiousness. PNAS 118, e2111477118.

Kilpatrick, A.M., Daszak, P., Jones, M.J., Marra, P.P., Kramer, L.D. (2006). Host heterogeneity dominates West Nile virus transmission. Proceedings of the Royal Society B 273, 2327–2333.

Komar, N., Langevin, S., Hinten, S., Nemeth, N., Edwards, E., Hettler, D., Davis, B., Bowen, R., Bunning, M. (2003). Experimental infection of North American birds with the New York 1999 strain of West Nile virus. Emerging Infectious Diseases 9, 311–322.

Langwig, K.E., Wargo, A.R., Jones, D.R., Viss, J.R., Rutan, B.J., Egan, N.A., et al. (2017). Vaccine effects on heterogeneity in susceptibility and implications for population health management. mBio 8, e00796-17.

Lloyd-Smith, J.O., Schreiber, S.J., Kopp, P.E., Getz, W.M. (2005). Superspreading and the effect of individual variation on disease emergence. Nature 438, 355–359.

LoGiudice, K., Ostfeld, R.S., Schmidt, K.A., Keesing, F. (2003). The ecology of infectious disease: effects of host diversity and community composition on Lyme disease risk. PNAS 100, 567–571.

Lunn, T.J., Restif, O., Peel, A.J., et al. (2019). Dose–response and transmission: the nexus between reservoir hosts, environment and recipient hosts. Philosophical Transactions of the Royal Society B 374, 20190016.

Marc, A., et al. (2021). Quantifying the relationship between SARS-CoV-2 viral load and infectiousness. eLife 10, e69302.

Mather, T.N., Wilson, M.L., Moore, S.I., Ribeiro, J.M.C., Spielman, A. (1989). Comparing the relative potential of rodents as reservoirs of the Lyme disease spirochete (Borrelia burgdorferi). American Journal of Epidemiology 130, 143–150.

Roberts, M.G., Heesterbeek, J.A.P. (2003). A new method for estimating the effort required to control an infectious disease. Proceedings of the Royal Society B 270, 1359–1364.

Roberts, M.G., Heesterbeek, J.A.P. (2013). Characterizing the next-generation matrix and basic reproduction number in ecological epidemiology. Journal of Mathematical Biology 66, 1045–1064.

Roberts, M.G., Heesterbeek, J.A.P. (2020). Characterizing reservoirs of infection and the maintenance of pathogens in ecosystems. Journal of the Royal Society Interface 17, 20190540.

Smith, D.L., Battle, K.E., Hay, S.I., Barker, C.M., Scott, T.W., McKenzie, F.E. (2012). Ross, Macdonald, and a theory for the dynamics and control of mosquito-transmitted pathogens. PLoS Pathogens 8, e1002588.

van den Driessche, P. (2017). Reproduction numbers of infectious disease models. Infectious Disease Modelling 2, 288–303.

Woolhouse, M.E.J., Dye, C., Etard, J.-F., Smith, T., Charlwood, J.D., Garnett, G.P., et al. (1997). Heterogeneities in the transmission of infectious agents: implications for the design of control programs. PNAS 94, 338–342.
