## Appendix S2 for "Formalizing host competence"

Appendix S2. Parameter values, distributions and sources

### S1. How to read the tables

**Parameter**: the quantity and its units.

**Distribution among individuals**: how each simulated individual's value is drawn.

**Host**: species, or "All" for shared values.

**Value**: the mean of the distribution, or the fixed value.

**SD used in simulation**: the spread of the distribution as coded.

**Source and basis**: the citation, with one of the tags below.

**Distributions:**

- **Lognormal** draws are mean-preserving: *X* = exp(ln *m* − *s*^2^/2 + *sZ*), *Z* ~ N(0, 1), so the arithmetic mean equals the stated value *m*. The SD column gives *s*, the SD on the natural-log scale ("log-SD").
- **Normal** draws have the stated arithmetic SD. Where a floor is given, draws below it are set to the floor; for West Nile virus duration, draws outside the observed range are set to the range limits.
- **Beta-Poisson** susceptibility is ε = 1 − (1 − *r*)*^D^*, with per-particle infection probability *r* ~ Beta(α, β) and dose *D* lognormal.
- **Fixed** values are the same for every individual (SD "—").

**Literature**: the value, or the data it is computed from, is published for this host and pathogen.

**Derived**: computed from published data by the stated rule.

**Consistent**: no direct estimate, but within a published range for this or a closely related host.

**Plausible**: no published estimate; the reasoning is in Section S6.

**Assumed**: a scenario or design choice.

### S2. Parameter tables

#### Table S1. West Nile virus (NY99 strain)

Five passerine and columbiform species infected experimentally by mosquito bite (Komar et al. 2003). Load units: log_10_ PFU/mL serum. Infectiousness is *a* = ∫ *b* φ(*L*(τ)) dτ over 7 days, with φ applied to log_10_ titer. Ranges are converted to SDs by dividing by *d*_2_(*n*), the expected range of *n* standard normal draws (1.128, 1.693, 2.059 and 2.534 for *n* = 2, 3, 4 and 6).

**Table S1. West Nile virus parameters.**

| **Parameter** | **Distribution among individuals** | **Host** | **Value** | **SD used in simulation** | **Source and basis** |
| --- | --- | --- | --- | --- | --- |
| Daily mean viremia, days 1–6 (log_10_ PFU/mL) | Fixed (linear interpolation between days; day 0 = 0; day 7 below detection) | Blue jay | 8.5, 11.1, 12.1, 10.5, 2.2, <1.7 (*n* = 4) | — | Literature: Komar et al. 2003, Table 2. "<1.7" (below detection) coded as 1.0 |
|  |  | House sparrow | 7.8, 9.8, 10.3, 10.3, 8.4, 1.8 (*n* = 6) | — |  |
|  |  | American robin | 5.8, 8.9, 7.3, 4.6, 2.0, <1.7 (*n* = 2) | — |  |
|  |  | European starling | 5.3, 6.1, 4.9, 2.3, <1.7, <1.7 (*n* = 6) | — |  |
|  |  | Mourning dove | 4.8, 5.9, 5.6, 3.3, <1.7, <1.7 (*n* = 3) | — |  |
| Among-bird offset in log_10_ viremia (load magnitude) | Normal, mean 0; one offset per bird, added to every day | Blue jay | 0 | 1.70 | Derived: mean of the day-1 and day-2 ranges ÷ *d*_2_(*n*) (Komar et al. 2003, Table 2), using only days before any bird died |
|  |  | House sparrow | 0 | 1.50 |  |
|  |  | American robin | 0 | 0.75 |  |
|  |  | European starling | 0 | 0.73 |  |
|  |  | Mourning dove | 0 | 1.39 |  |
| Duration of detectable viremia (d) | Normal, limited to the observed range | Blue jay | 4.0 (range 3–5) | 0.97 | Literature (mean, range): Komar et al. 2003, Table 2 (viremia ≥10^1.7^); SD derived as range ÷ *d*_2_(*n*). Because φ = 0 below 10^5^, infectious duration is set by the titer |
|  |  | House sparrow | 4.5 (range 2–6) | 1.58 |  |
|  |  | American robin | 4.5 (range 4–5) | 0.89 |  |
|  |  | European starling | 3.2 (range 3–4) | 0.39 |  |
|  |  | Mourning dove | 3.7 (range 3–4) | 0.59 |  |
| Infectiousness to *Culex*, φ | Fixed | All | Hill on log_10_ titer: half-saturation 8.788 (10^8.788^ PFU/mL), *n* = 4.486; φ = 0 below 10^5^ | — | Derived: least-squares fit to mean daily infectiousness *i* (Komar et al. 2003, Table 10). Threshold: Komar et al. 2003; Sardelis et al. 2001 |
| Behavior *b* | Lognormal | All | 1 | 0.15 | Plausible (S6.1). Direction of among-bird differences: Darbro et al. 2007; Gervasi et al. 2016 |
| Host survival *S*(τ) | Fixed | All | 1 (not modeled) | — | Daily means after the first deaths are from survivors (Komar et al. 2003; S6.6) |
| Susceptibility ε | Fixed | All | 1 | — | Literature: Komar et al. 2003, Table 10 (s = 1.0). |
| Biting rate θ (per day) | Lognormal | All | 0.3 | 0.3 | Magnitude consistent with Caillouët et al. 2013 |
| Reservoir competence index *C*_i_ (comparison only) | Fixed | Blue jay | 2.55 | — | Literature: Komar et al. 2003, Table 10. Compared with simulated mean *a* in Fig. 3C |
|  |  | House sparrow | 1.59 | — |  |
|  |  | American robin | 1.08 | — |  |
|  |  | European starling | 0.22 | — |  |
|  |  | Mourning dove | 0.19 | — |  |
| Integration window; step (d) | Fixed | All | 7; 0.05 | — | All five species below detection by day 7 (Komar et al. 2003, Table 2) |

#### Table S2. Lyme disease (*Borrelia burgdorferi*) host community

Twelve hosts of larval *Ixodes scapularis*. Load units are relative to the half-saturation constant *K* = 10^4^; only *L*_max_/*K* affects φ. Infectiousness is *a* = *S* ∫ φ(*L*(τ)) dτ over a 60-day larval season, with logistic load growth *L*(τ) = *L*_max_ / (1 + exp(−*r*(τ − *t*_0_))). Infected larvae per infected host (Fig. 3A) = *a*θ; competence *H*_comp_ = ε*a*, potential *H*_pot_ = θε*a*, and capacity *H*_cap_ = *D*θε*a* (Fig. 4).

##### Table S2a. Within-host parameters

**Table S2a. Lyme disease: within-host parameters and feeding survival.**

| **Parameter** | **Distribution among individuals** | **Host** | **Value** | **SD used in simulation** | **Source and basis** |
| --- | --- | --- | --- | --- | --- |
| Susceptibility ε (probability of infection after an infected nymph bite) | Fixed | White-footed mouse | 0.95 | — | Plausible (S6.2). Ranking follows field prevalence and infectivity (Mather et al. 1989; Brunner et al. 2008). Mouse and short-tailed shrew values match the prevalence estimates of Brunner et al. 2008, Table 1 |
|  |  | Eastern chipmunk | 0.75 | — |  |
|  |  | Short-tailed shrew | 0.85 | — |  |
|  |  | Masked shrew | 0.75 | — |  |
|  |  | Eastern gray squirrel | 0.25 | — |  |
|  |  | Virginia opossum | 0.08 | — |  |
|  |  | Raccoon | 0.15 | — |  |
|  |  | Striped skunk | 0.15 | — |  |
|  |  | White-tailed deer | 0.05 | — |  |
|  |  | American robin | 0.65 | — |  |
|  |  | Veery | 0.50 | — |  |
|  |  | Wood thrush | 0.40 | — |  |
| Plateau load *L*_max_ (relative units) | Lognormal | White-footed mouse | 5×10^4^ | 0.5 | Short-tailed shrew: Calibrated (v9) so ε × φ(*L*_max_) equals published reservoir competence, 0.418 (LoGiudice et al. 2003); φ(*L*_max_) 0.49 vs infectivity 0.505 (Brunner et al. 2008, Table 1). Consistent where ε × φ(*L*_max_) approximates published reservoir competence: mouse, chipmunk, masked shrew, raccoon and veery (LoGiudice et al. 2003; Keesing et al. 2009). Plausible otherwise (S6.2; Table S6) |
|  |  | Eastern chipmunk | 2×10^4^ | 0.6 |  |
|  |  | Short-tailed shrew | 9.8×10^3^ | 0.6 |  |
|  |  | Masked shrew | 2×10^4^ | 0.7 |  |
|  |  | Eastern gray squirrel | 5×10^3^ | 0.9 |  |
|  |  | Virginia opossum | 1.5×10^3^ | 1.0 |  |
|  |  | Raccoon | 2×10^3^ | 0.8 |  |
|  |  | Striped skunk | 2.5×10^3^ | 0.8 |  |
|  |  | White-tailed deer | 500 | 1.0 |  |
|  |  | American robin | 2×10^4^ | 0.7 |  |
|  |  | Veery | 1.2×10^4^ | 0.7 |  |
|  |  | Wood thrush | 8×10^3^ | 0.8 |  |
| Load growth rate *r* (per day) | Normal, floor 0.01 | White-footed mouse | 0.35 | 0.06 | White-footed mouse: Consistent (Donahue et al. 1987; Lindsay et al. 1997; Bourgeois et al. 2024). American robin: rise slower than observed (Richter et al. 2000; S6.6). Others: Plausible; no time course published (S6.2) |
|  |  | Eastern chipmunk | 0.25 | 0.05 |  |
|  |  | Short-tailed shrew | 0.30 | 0.06 |  |
|  |  | Masked shrew | 0.28 | 0.06 |  |
|  |  | Eastern gray squirrel | 0.15 | 0.04 |  |
|  |  | Virginia opossum | 0.10 | 0.03 |  |
|  |  | Raccoon | 0.12 | 0.04 |  |
|  |  | Striped skunk | 0.12 | 0.04 |  |
|  |  | White-tailed deer | 0.08 | 0.03 |  |
|  |  | American robin | 0.25 | 0.06 |  |
|  |  | Veery | 0.20 | 0.05 |  |
|  |  | Wood thrush | 0.18 | 0.05 |  |
| Time to half plateau *t*_0_ (d) | Normal, floor 1 d | White-footed mouse | 10 | 2 | As for *r* |
|  |  | Eastern chipmunk | 12 | 3 |  |
|  |  | Short-tailed shrew | 10 | 2 |  |
|  |  | Masked shrew | 10 | 2 |  |
|  |  | Eastern gray squirrel | 18 | 4 |  |
|  |  | Virginia opossum | 20 | 5 |  |
|  |  | Raccoon | 18 | 4 |  |
|  |  | Striped skunk | 16 | 4 |  |
|  |  | White-tailed deer | 20 | 5 |  |
|  |  | American robin | 12 | 3 |  |
|  |  | Veery | 12 | 3 |  |
|  |  | Wood thrush | 14 | 3 |  |
| Feeding survival *S* (proportion of attached larvae that feed to repletion; constant factor in *a*) | Lognormal | White-footed mouse | 0.49 | 0.05 | Literature: Keesing et al. 2009, via Levi et al. 2016, Table 1 |
|  |  | Eastern chipmunk | 0.24 | 0.08 |  |
|  |  | Short-tailed shrew | 0.49 | 0.10 | Assumed by Levi et al. 2016, Table 1: mouse value (their alternative: 0.24, chipmunk value) |
|  |  | Masked shrew | 0.49 | 0.10 | Assumed by Levi et al. 2016, Table 1: mouse value (their alternative: 0.24) |
|  |  | Eastern gray squirrel | 0.17 | 0.15 | Literature: Keesing et al. 2009, via Levi et al. 2016, Table 1 |
|  |  | Virginia opossum | 0.035 | 0.20 |  |
|  |  | Raccoon | 0.24 | 0.15 | Assumed by Levi et al. 2016, Table 1: chipmunk value (their alternative: 0.04, opossum value) |
|  |  | Striped skunk | 0.24 | 0.15 | Assumed by Levi et al. 2016, Table 1: chipmunk value (their alternative: 0.04) |
|  |  | White-tailed deer | 0.49 | 0.10 | Assumed by Levi et al. 2016, Table 1: poor groomer, mouse value |
|  |  | American robin | 0.27 | 0.12 | Levi et al. 2016, Table 1: ground-foraging birds (Keesing et al. 2009 veery value), applied to each bird species |
|  |  | Veery | 0.27 | 0.12 | Literature: Keesing et al. 2009, via Levi et al. 2016, Table 1 |
|  |  | Wood thrush | 0.27 | 0.12 | As for American robin |

Among-individual log-SDs for *S* are assumed; no study reports individual variation in feeding success (S6.2). Because the lognormal is mean-preserving, species means of *a*, *H*_pot_ and *H*_cap_, and therefore the ranks in Fig. 4, do not depend on them.

##### Table S2b. Host ecology and contact

**Table S2b. Lyme disease: host ecology and contact.**

| **Parameter** | **Distribution among individuals** | **Host** | **Value** | **SD used in simulation** | **Source and basis** |
| --- | --- | --- | --- | --- | --- |
| Host density *D* (individuals per ha) | Fixed | White-footed mouse | 40 | — | Literature: Levi et al. 2016, Table 1 (compiled from LoGiudice et al. 2003; Keesing et al. 2009) |
|  |  | Eastern chipmunk | 20 | — |  |
|  |  | Short-tailed shrew | 25 | — |  |
|  |  | Masked shrew | 25 | — |  |
|  |  | Eastern gray squirrel | 8.1 | — |  |
|  |  | Virginia opossum | 1 | — | Literature: LoGiudice et al. 2003; Keesing et al. 2009 (Levi et al. 2016 use 0.2 as base and 1 as alternative; S6.2) |
|  |  | Raccoon | 0.2 | — | Literature: Levi et al. 2016, Table 1 (compiled from LoGiudice et al. 2003; Keesing et al. 2009) |
|  |  | Striped skunk | 0.05 | — |  |
|  |  | White-tailed deer | 0.25 | — |  |
|  |  | American robin | 15 | — | Assumed split of the pooled ground-foraging bird density, 31.6 per ha (LoGiudice et al. 2003; Levi et al. 2016); the three values sum to 29 |
|  |  | Veery | 8 | — |  |
|  |  | Wood thrush | 6 | — |  |
| Post-grooming burden *B* (engorged larvae per host per 72 h) | Fixed | White-footed mouse | 27.8 | — | Literature: LoGiudice et al. 2003, Table 1, via Levi et al. 2016, Table 1 |
|  |  | Eastern chipmunk | 36 | — |  |
|  |  | Short-tailed shrew | 62.9 | — |  |
|  |  | Masked shrew | 55.5 | — |  |
|  |  | Eastern gray squirrel | 142 | — |  |
|  |  | Virginia opossum | 254 | — |  |
|  |  | Raccoon | 127 | — |  |
|  |  | Striped skunk | 66.8 | — |  |
|  |  | White-tailed deer | 239 | — |  |
|  |  | American robin | 8 | — | Plausible: within the per-bird range used by Levi et al. 2016 (1.7, LoGiudice et al. 2003; 11.4, Keesing et al. 2009) |
|  |  | Veery | 6 | — |  |
|  |  | Wood thrush | 4 | — |  |
| Larvae attaching per host per 72 h, *B*/*S* | Fixed (derived) | White-footed mouse | 56.7 | — | Derived: post-grooming burden ÷ feeding survival (Levi et al. 2016, Eqs 16–17) |
|  |  | Eastern chipmunk | 150.0 | — |  |
|  |  | Short-tailed shrew | 128.4 | — |  |
|  |  | Masked shrew | 113.3 | — |  |
|  |  | Eastern gray squirrel | 835.3 | — |  |
|  |  | Virginia opossum | 7,257.1 | — |  |
|  |  | Raccoon | 529.2 | — |  |
|  |  | Striped skunk | 278.3 | — |  |
|  |  | White-tailed deer | 487.8 | — |  |
|  |  | American robin | 29.6 | — |  |
|  |  | Veery | 22.2 | — |  |
|  |  | Wood thrush | 14.8 | — |  |
| Contact θ = *B*/(3*S*) (larvae attaching per host per day) | Fixed (derived) | White-footed mouse | 18.9 | — | Derived. Among-individual variation in contact is used only in Fig. 2 (Table S5) |
|  |  | Eastern chipmunk | 50.0 | — |  |
|  |  | Short-tailed shrew | 42.8 | — |  |
|  |  | Masked shrew | 37.8 | — |  |
|  |  | Eastern gray squirrel | 278.4 | — |  |
|  |  | Virginia opossum | 2,419.0 | — |  |
|  |  | Raccoon | 176.4 | — |  |
|  |  | Striped skunk | 92.8 | — |  |
|  |  | White-tailed deer | 162.6 | — |  |
|  |  | American robin | 9.9 | — |  |
|  |  | Veery | 7.4 | — |  |
|  |  | Wood thrush | 4.9 | — |  |
| Realized reservoir competence (comparison only) | Fixed | White-footed mouse | 0.921 | — | Literature: LoGiudice et al. 2003, Table 1, via Levi et al. 2016, Table 1 (birds pooled). Used only in the check against mean *a* (Section 5; Table S5) |
|  |  | Eastern chipmunk | 0.550 | — |  |
|  |  | Short-tailed shrew | 0.418 | — |  |
|  |  | Masked shrew | 0.512 | — |  |
|  |  | Eastern gray squirrel | 0.147 | — |  |
|  |  | Virginia opossum | 0.026 | — |  |
|  |  | Raccoon | 0.013 | — |  |
|  |  | Striped skunk | 0.097 | — |  |
|  |  | White-tailed deer | 0.046 | — |  |
|  |  | American robin | 0.117 | — |  |
|  |  | Veery | 0.117 | — |  |
|  |  | Wood thrush | 0.117 | — |  |

##### Table S2c. Shared parameters

**Table S2c. Lyme disease: shared parameters.**

| **Parameter** | **Distribution among individuals** | **Host** | **Value** | **SD used in simulation** | **Source and basis** |
| --- | --- | --- | --- | --- | --- |
| Infectiousness to larvae, φ | Fixed | All | Hill on load: *K* = 10^4^ (relative units), *n* = 1.5 | — | *K* is a scale convention (only *L*_max_/*K* is identifiable). *n*: Derived (approximate) from Zinck et al. 2023; the individual-level relation is untested (Bourgeois et al. 2024) |
| Behavior *b* (infection-induced change in contact) | Fixed | All | 1 | — | Assumed; the *b* slot in the code carries *S* (Table S2a) |
| Host survival *S*(τ) within the season | Fixed | All | 1 (not modeled) | — | *B. burgdorferi* is not lethal to these hosts; the season bounds the integral |
| Transmission window; step (d) | Fixed | All | 60; 1 | — | Consistent: larval peak in August–September (LoGiudice et al. 2003; Ostfeld et al. 2006; Huang et al. 2019) |

#### Table S3. Chytridiomycosis (*Batrachochytrium dendrobatidis*, Bd)

Load units: zoospore equivalents (ZE) per swab. Load grows logistically from *L*_0_ toward *L*_cap_ at rate *g*. Infectiousness is *a* = *b* ∫ φ(*L*(τ)) *S*(τ) dτ, bounded by natural lifespan (bullfrog) or by background plus load-dependent disease mortality (northern leopard frog, NLF; mountain yellow-legged frog, MYLF).

**Table S3. Chytridiomycosis parameters.**

| **Parameter** | **Distribution among individuals** | **Host** | **Value** | **SD used in simulation** | **Source and basis** |
| --- | --- | --- | --- | --- | --- |
| Initial load *L*_0_ (ZE) | Lognormal | American bullfrog | 100 | 0.5 | Plausible (S6.3): Voyles et al. 2012a; DiRenzo et al. 2014; Jani et al. 2021 |
|  |  | Northern leopard frog | 100 | 0.5 |  |
|  |  | Mountain yellow-legged frog | 100 | 0.5 |  |
| Load growth rate *g* (per day) | Normal, floor 0.001 | American bullfrog | 0.08 | 0.02 | NLF, MYLF: Plausible (S6.3). Bullfrog: experimental bullfrogs usually clear or nearly clear infection (Byrne et al. 2018; S6.6) |
|  |  | Northern leopard frog | 0.1 | 0.025 |  |
|  |  | Mountain yellow-legged frog | 0.12 | 0.03 |  |
| Load capacity *L*_cap_ (ZE) | Lognormal | American bullfrog | 5×10^5^ | 0.6 | MYLF: Consistent (Knapp et al. 2022). Bullfrog and NLF: above reported loads (Greenspan et al. 2012; Buck et al. 2015; S6.6) |
|  |  | Northern leopard frog | 1.5×10^6^ | 0.55 |  |
|  |  | Mountain yellow-legged frog | 2×10^6^ | 0.5 |  |
| Mean natural lifespan (d) | Lognormal | American bullfrog | 1,825 | 0.4 | Plausible (S6.3): AnAge |
| Background mortality hazard *h*_0_ (per day) | Fixed | Northern leopard frog | 0.001 | — | Consistent: annual survival 0.69 (Knapp et al. 2022; AnAge) |
|  |  | Mountain yellow-legged frog | 0.001 | — |  |
| Load threshold for disease mortality (ZE) | Fixed | Northern leopard frog | 3×10^4^ | — | MYLF: Consistent (Vredenburg et al. 2010). NLF: adult Bd mortality not documented (Voordouw et al. 2010; S6.6); froglets die (Pask et al. 2013) |
|  |  | Mountain yellow-legged frog | 5×10^3^ | — |  |
| Disease-mortality slope β (per ZE above threshold per day) | Lognormal | Northern leopard frog | 8×10^-8^ | 0.4 | MYLF: Consistent; implied median time to death 63 d vs mortality from 3–4 weeks after exposure (Jani & Briggs 2018). NLF: S6.6 |
|  |  | Mountain yellow-legged frog | 5×10^-7^ | 0.4 |  |
| Behavior *b* | Lognormal | American bullfrog | 1 | 0.2 | Plausible; direction supported (Jani & Briggs 2018; Voyles et al. 2012b; Chatfield et al. 2013; Sauer et al. 2018) |
|  |  | Northern leopard frog | 0.85 | 0.25 |  |
|  |  | Mountain yellow-legged frog | 0.7 | 0.3 |  |
| Susceptibility ε (beta-Poisson) | *r* ~ Beta(α, β); dose lognormal | American bullfrog | *r* ~ Beta(3, 500); dose 1,000 zoospores; mean ε 0.95 | dose 0.3 | Bullfrog: higher than reported (Peterson & McKenzie 2014). MYLF: Consistent (Jani & Briggs 2018) |
|  |  | Northern leopard frog | *r* ~ Beta(4, 400); dose 1,000 zoospores; mean ε 0.99 | dose 0.3 |  |
|  |  | Mountain yellow-legged frog | *r* ~ Beta(5, 300); dose 1,000 zoospores; mean ε 1.00 | dose 0.3 |  |
| Integration window; step (d) | Fixed | American bullfrog | 9,000; 2 | — | Numerical horizons; survival near zero by the window end for NLF and MYLF |
|  |  | Northern leopard frog | 540; 1 | — |  |
|  |  | Mountain yellow-legged frog | 120; 0.5 | — |  |
| Shedding/infectiousness, φ | Fixed | All | Hill on load: *K* = 10^4^ ZE, *n* = 2.5 | — | Assumed: no load threshold reported; shedding rises with load (Reeder et al. 2012; DiRenzo et al. 2014; Wilber et al. 2022; S6.6) |
| Contact × zoospore persistence, θ (per day) | Lognormal × Beta(5, 5) | All | 2.0 × 0.5 | 0.4 (contact) | Placeholder. Zoospore death rates: Voyles et al. 2012a; seasonal contact: Wilber et al. 2022 |

#### Table S4. Low-pathogenicity avian influenza (LPAI)

Load units: EID_50_/mL of swab medium. Shedding follows *L*(τ) = *L*_max_ (τ/τ_p_)^α^ exp(−α(τ/τ_p_ − 1)), peaking at τ_p_. Infectiousness is *a* = *b* ∫ φ(*L*(τ)) *S*(τ) dτ over 21 days, with *S*(τ) = exp(−*h*_0_τ).

**Table S4. Low-pathogenicity avian influenza parameters.**

| **Parameter** | **Distribution among individuals** | **Host** | **Value** | **SD used in simulation** | **Source and basis** |
| --- | --- | --- | --- | --- | --- |
| Peak load *L*_max_ (EID_50_/mL) | Lognormal | Mallard | 5×10^4^ | 0.7 | Consistent: Brown et al. 2012a; Criado et al. 2021 |
|  |  | Wood duck | 1.5×10^4^ | 0.8 | Plausible: no titers published; shorter shedding than mallards (Costa et al. 2011b; S6.4) |
|  |  | Laughing gull | 3×10^3^ | 1.0 | Depends on virus: 10^1.5^–10^6.3^ across gull-origin viruses (Criado et al. 2021); other origins rarely infect (Bahnson et al. 2020; S6.6) |
| Time to peak shedding τ_p_ (d) | Normal, floor 0.5 d | Mallard | 2.5 | 0.625 | Literature: peak shedding 2–3 d after inoculation (Brown et al. 2012a); SD scaled to keep the v7 CV |
|  |  | Wood duck | 3 | 0.8 | Plausible (S6.4) |
|  |  | Laughing gull | 2 | 0.6 | Consistent: early peaks (day 1) for all virus groups (Criado et al. 2021) |
| Shedding curve shape α | Fixed | Mallard | 3 | — | Plausible: gives about 2 weeks of shedding in mallards (Brown et al. 2012a; Latorre-Margalef et al. 2009) |
|  |  | Wood duck | 3 | — |  |
|  |  | Laughing gull | 4 | — |  |
| Behavior *b* | Lognormal | Mallard | 1 | 0.1 | Literature: no behavioral change after infection (mallard, Jourdain et al. 2010; wood duck, Costa et al. 2011b; laughing gull, Criado et al. 2021). SDs assumed |
|  |  | Wood duck | 1 | 0.15 |  |
|  |  | Laughing gull | 1 | 0.2 |  |
| Background mortality hazard *h*_0_ (per day) | Fixed | All | 5×10^-4^ | — | Annual survival 0.83; published adult survival 0.54–0.62 (Roberts et al. 2023; Otis & Dukes 1995). Changes 21-day survival by at most 2.4% |
| Susceptibility ε (beta-Poisson) | *r* ~ Beta(α, β); dose lognormal | Mallard | *r* ~ Beta(6, 120); dose 10^4^ EID_50_; mean ε 1.000 | dose 0.3 | Dose: Brown et al. 2012b. As coded, ε ≈ 1 for all three species; experiments separate them on infection success (S6.6) |
|  |  | Wood duck | *r* ~ Beta(4, 160); dose 10^4^ EID_50_; mean ε 1.000 | dose 0.3 |  |
|  |  | Laughing gull | *r* ~ Beta(2.5, 220); dose 10^4^ EID_50_; mean ε 1.000 | dose 0.4 |  |
| Shedding/infectiousness, φ | Fixed | All | Hill on load: *K* = 10^4^ EID_50_/mL, *n* = 2 | — | *K*: dose used to mimic natural exposure (Brown et al. 2012b). *n*: Assumed (Breban et al. 2009 use exponential escape) |
| Contact × persistence in water, θ (per day) | Lognormal × Beta(6, 4) | All | 2.0 × 0.6 | 0.3 (contact) | Placeholder.  . Persistence 9–100 d (Stallknecht et al. 1990; Brown et al. 2009); contact consistent with Niqueux et al. 2013 |
| Integration window; step (d) | Fixed | All | 21; 0.1 | — | Shedding ends within about 2 weeks (Brown et al. 2012a; Latorre-Margalef et al. 2009) |

#### Table S5. Cross-system and analysis settings

These are design choices rather than biological estimates; the contact CV and the Fig. 3B band have literature anchors.

**Table S5.** Cross-system and analysis settings.

| **Setting** | **Value** | **Used in** | **Basis and source** |
| --- | --- | --- | --- |
| Monte Carlo sample | 4,000 individuals per species | Figs 1, 3, 4 | Numerical choice. Seed 20260925 (R set.seed; Python numpy.random.default_rng). The two pipelines use different random-number streams and agree within Monte Carlo error |
| Covariation sample | 40,000 individuals per species (8,000 for the bullfrog) | Fig. 2 | Numerical choice |
| Among-individual CV of contact | 1 (lognormal; contact expressed relative to its mean) | Fig. 2 | Literature for Lyme: larval burdens on white-footed mice, CV 0.95–1.16 (Tufts et al. 2021); negative binomial *k* = 0.86 (Devevey & Brisson 2012). Applied to the other systems as an extrapolation; nestling biting data suggest a larger CV for West Nile virus (Caillouët et al. 2013) |
| Correlation ρ between load magnitude and contact | −0.7, −0.4, 0, 0.4, 0.7 (Gaussian copula; marginal distributions unchanged) | Fig. 2 | Assumed scenario range; no estimates exist. Mice that carry many ticks in one season tend to carry many in others (Devevey & Brisson 2012), which supports a positive correlation but gives no value |
| Self-replacement benchmark | Mean contact at which 10% of individuals replace themselves at ρ = 0 (1 ÷ 90th percentile of θ*a*) | Fig. 2 | Design choice; holds mean contact fixed across ρ |
| Exposure–transmission contact inflation | exp(ρσ^2^), σ^2^ = ln(1 + CV^2^) | Main text Section 2; Appendix S1, Sections 5.4–5.5 | Analytical result (Dye & Hasibeder 1986) |
| Sobol sensitivity design | First- and total-order indices of *a*; *N* = 16,384 base samples (4,096 for the bullfrog); 200 bootstrap resamples for 95% intervals; inputs drawn from the simulation distributions | Appendix S3, Fig. S1 | Saltelli and Jansen estimators (Saltelli et al. 2010) |
| Sobol centering (v9) | Outputs centered on the mean of the two base samples before the first-order estimator is applied | Appendix S3, Fig. S1 | Numerical stabilization; leaves the expected index unchanged. Across eight seeds the mouse load-magnitude index ranged 0.62–0.88 without centering and 0.74–0.77 with it |
| Fig. 3B band of tick-side probability *q* | 0.0081 to 0.35 | Fig. 3B | Product of: survival through molt and winter, 0.18–0.47 (Lindsay et al. 1998); nymphal host-finding, 0.15–0.83 (Tufts et al. 2021); infection of the fed-upon host, 0.30–0.90 (Assumed; S6.2) |
| Break-even contact | West Nile virus: θ *= 1/*a *(median among birds with* a *> 0). Lyme:* q* *= 1/(*a*θ) | Fig. 3 | Definition (Eq. 6) |
| Reservoir-competence check | Least-squares regression of log_10_ realized reservoir competence on log_10_ mean *a*, nine mammals; Spearman rank correlation | Section 5 | Reservoir competence from LoGiudice et al. 2003 (Table S2b) |
| Numerical integration | Trapezoidal rule; steps 0.05 d (West Nile virus) to 2 d (bullfrog) | All | Steps finer than the fastest load change in each system (Tables S1–S4) |
| Box 1 inputs | Individual viremia of zebra finches with and without corticosterone, interpolated linearly between days 0, 2, 4, 6 and 14; birds that died held at their last titer; φ as in Table S1; ε = 1; step 0.01 d; attractiveness ×2 for treated birds | Box 1 | Data: Gervasi et al. 2017 (Dryad, doi:10.5061/dryad.1s9g1). Attractiveness: Gervasi et al. 2016 |

### S6. Biological plausibility of values with no literature estimate

Each note covers a value tagged Plausible or Assumed in Tables S1–S5: what bounds it, what it was chosen to represent, and where it is only an assumption. Section S6.6 lists caveats for decisions made within the parameterization.

#### S6.1 West Nile virus

**Infectiousness function φ.** Komar et al. (2003) report infectiousness only as a mean per infectious day for each species. Applied to the daily means, the refit Hill function (half-saturation 10^8.79^, *n* = 4.49) gives 0.68, 0.56, 0.32, 0.13 and 0.13 for blue jay, house sparrow, American robin, European starling and mourning dove, against published values of 0.68, 0.53, 0.36, 0.12 and 0.11 (our calculation).

**Behavior *b* (mean 1, log-SD 0.15).** No study reports a variance in per-bird feeding success, so the mean of 1 simply defines *b* relative to the bird's pre-infection state. The SD is conservative: mosquitoes consistently chose one of two healthy house finches, foot-stomp rate explained about half the difference, and stressed birds were fed on twice as often (Darbro et al. 2007; Gervasi et al. 2016). An among-bird CV of about 15% is therefore best read as a lower bound on behavioral heterogeneity.

**Biting rate θ (0.3 per day).** No source gives bites per adult bird per day. Nestlings receive 0.7 attempted feeds per night on average (1.8 for house sparrows), so 0.3 successful bites per day is the right order of magnitude once failed attempts are excluded (Caillouët et al. 2013).

#### S6.2 Lyme disease

**Susceptibility ε.** ε is the probability that a host becomes infected after an infected nymph bite. Field prevalence is related to ε but also depends on how often hosts are bitten (90% in mice, 75% in chipmunks; Mather et al. 1989). Across species, prevalence and infectivity are strongly correlated (*r* = 0.88; Brunner et al. 2008), and the coded ε values preserve that ranking: shrews intermediate, squirrels low, opossum and raccoon very low. Brunner et al. call the opossum and raccoon estimates "uninformative" because single positive ticks from these hosts may be false positives, so those two values are assumptions with wide uncertainty. Their Table 1 separates prevalence (π) from infectivity (φ) for the LoGiudice hosts, which lets ε and φ(*L*_max_) be checked separately rather than only as a product. Assuming a perfect test, the mouse (π 1.0, φ 0.914) and short-tailed shrew (π 0.831, φ 0.505) match the coded values closely (ε 0.95, φ 0.92; ε 0.85, φ 0.49). For the chipmunk (π 0.971, φ 0.569) and masked shrew (π 0.701, φ 0.537), the products agree with the coded values but the coded split places less in ε (0.75) and more in φ (0.74) (S6.6). For squirrels, deer, raccoon, opossum and skunk, perfect-test prevalence (0.31–1.0) is well above the coded ε (0.05–0.25); allowing for false positives lowers it to 0.0–0.28.

**Plateau load and growth (*L*_max_, *r*, *t*_0_) for the species without a time course.** The plateau ratio *L*_max_/*K* is the only identifiable quantity. Table S6 compares ε × φ(*L*_max_) with published reservoir competence. Slower rise for low-competence hosts (*r* 0.08–0.15 vs 0.35 per day) is an assumption: body size predicts the plateau, but a spirochete growth model predicts only a 1.5–4-day difference in lag between a mouse and a raccoon (Barbour et al. 2015). The assumption matters: relative to a host at plateau from day 0, the rise lowers 60-day infectiousness by 10–17% in the competent species but by 34–46% in squirrel, opossum, raccoon, skunk and deer, widening the gap between high- and low-competence hosts by up to about 1.7-fold (our calculation; feeding survival multiplies both cases equally and does not change these percentages).

**Table S6.** Lyme disease: model ε × φ(*L*_max_) compared with published realized reservoir competence (RC). RC: LoGiudice et al. (2003, Table 1), birds pooled; Keesing et al. (2009) report 28% for the veery and Ginsberg et al. (2005) 16.1% for wild-caught robins.

| **Host** | **ε** | **φ(*L*_max_)** | **ε × φ(*L*_max_) (%)** | **Published RC (%)** |
| --- | --- | --- | --- | --- |
| White-footed mouse | 0.95 | 0.92 | 87.2 | 92.1 |
| Eastern chipmunk | 0.75 | 0.74 | 55.4 | 55.0 |
| Short-tailed shrew | 0.85 | 0.49 | 41.9 | 41.8 |
| Masked shrew | 0.75 | 0.74 | 55.4 | 51.2 |
| Eastern gray squirrel | 0.25 | 0.26 | 6.5 | 14.7 |
| Virginia opossum | 0.08 | 0.05 | 0.4 | 2.6 |
| Raccoon | 0.15 | 0.08 | 1.2 | 1.3 |
| Striped skunk | 0.15 | 0.11 | 1.7 | 9.7 |
| White-tailed deer | 0.05 | 0.01 | 0.1 | 4.6 |
| American robin | 0.65 | 0.74 | 48.0 | 11.7 |
| Veery | 0.50 | 0.57 | 28.4 | 11.7 |
| Wood thrush | 0.40 | 0.42 | 16.7 | 11.7 |

**Hill steepness *n* = 1.5.** A logistic relation between infection probability and log load is a Hill function. Across 11 strains, the slope of nymphal infection on log ear load implies *n* between about 0.4 and 2 (Zinck et al. 2023), and 1.5 lies inside that interval. Whether the same relation holds among individual *Peromyscus leucopus* is untested; one study found no correlation (Bourgeois et al. 2024).

**Feeding survival *S*.** *S* was measured by placing 100 larvae on each host and counting those that fed to repletion (Keesing et al. 2009) for the mouse, chipmunk, gray squirrel, opossum and veery. For the other hosts, the values are the assumptions of Levi et al. (2016): shrews groom like mice (alternative: like chipmunks), raccoons and skunks like chipmunks (alternative: like opossums), and deer are poor groomers like mice. Under the alternatives, shrew contact would roughly double while their competence would halve, and raccoon and skunk contact would rise sixfold while their competence would fall sixfold (our arithmetic from Table S2). The among-individual log-SDs (0.05–0.20) are assumptions; no study reports individual variation in feeding success.

**Bird burdens, densities and *S*.** Levi et al. (2016) pool ground-foraging birds (density 31.6 per ha; burden 1.7 or 11.4 larvae per bird; *S* = 0.27). The per-species values here (burdens 8, 6 and 4; densities 15, 8 and 6 per ha) lie within those bounds, and all three birds take *S* = 0.27.

**Opossum density (1 per ha).** This is the value of LoGiudice et al. (2003) and Keesing et al. (2009); Levi et al. (2016) use 0.2 per ha as their base value. Opossums contribute less than 0.1% of community capacity at 1 per ha; at 0.2 per ha their share would be five times smaller.

**Fig. 3B band, probability that the fed-upon host becomes infected (30–90%).** No source gives this number. The range spans the coded ε of the hosts most nymphs feed on (mouse 0.95, chipmunk 0.75, robin 0.65) and allows for nymphs feeding on low-ε hosts.

#### S6.3 Chytridiomycosis

**Initial load *L*_0_ = 100 ZE.** One encysted zoospore produces its first sporangium in 2.5–3.9 days at 23 °C (Voyles et al. 2012a). A natural, low-dose exposure should therefore yield on the order of 10^1^–10^2^ ZE in the first week; high experimental doses give 10^3^–10^4^ (Jani et al. 2021). *L*_0_ matters mainly through time to death, ln(*L*_cap_/*L*_0_)/*g*.

**Growth rates for leopard frog and MYLF (0.10, 0.12 per day).** These are slower than in hypersusceptible *Atelopus* (about 0.30 per day; DiRenzo et al. 2014) and far below growth in culture at 23 °C (about 0.6), which suits cool montane and temperate hosts. The MYLF value gives a doubling time of about 6 days and median death at about 9 weeks, within a factor of 2–3 of experimental mortality onset (3–4 weeks; Jani & Briggs 2018). On-host growth depends on temperature (Wilber et al. 2022), so a constant rate is a simplification.

**Behavior *b* (0.85, 0.70).** Infected *Rana* lose appetite and weight, moribund MYLF show abnormal posture, and leopard frog jumping velocity declines (Jani & Briggs 2018; Voyles et al. 2012b; Chatfield et al. 2013). The direction is supported; the magnitudes are illustrative.

**Bullfrog lifespan (5 years).** It implies annual survival of about 0.82 and a 4% chance of reaching the 16-year captive maximum (AnAge). No wild survival estimate was found. Bullfrog infectiousness scales almost linearly with this value, so it should be described as an assumption.

**Contact rate and zoospore persistence.** No encounter rates have been measured. Zoospores die at about 1 per day at 23 °C and 0.13 per day at 4 °C (Voyles et al. 2012a); with a contact rate of 2 per day, the fraction of zoospores encountered before dying is 2/(2 + 1) ≈ 0.67 at warm temperatures, so a mean persistence factor of 0.5 is of the right order. Leopard frog contact should be seasonal (zero on land; Wilber et al. 2022).

#### S6.4 Low-pathogenicity avian influenza

**Wood duck peak load and timing.** No titers exist. Wood ducks shed detectable virus for 2–4 days, compared with 4–9 days in mallards given related viruses (Costa et al. 2011b; Costa et al. 2010), and a shorter window at the same detection limit implies a lower peak. Placing wood duck half a log_10_ below mallard, peaking by day 3, is an interpolation consistent with those bounds.

**Hill *K* = 10^4^ EID_50_/mL.** No study relates shedding titer to transmission. *K* sits at the dose one challenge study chose to mimic natural exposure (Brown et al. 2012b), so it marks the titer at which infection from one exposure becomes likely. *n* = 2 is a modeling choice; the only mechanistic form published is exponential escape (Breban et al. 2009).

**Behavior SDs (0.10–0.20).** With *b* = 1 for all three species, the SDs represent individual differences in contact unrelated to infection. They are assumptions; in the Sobol analysis (Appendix S3, Fig. S1) behavior has first-order indices of 0.05 (mallard) and 0.01 (laughing gull).

**Persistence factor and contact rate.** Persistence is published in days (9–100 days depending on temperature and salinity), which cannot be converted to a 0–1 factor. A contact rate of 2 per day is consistent with *R*_0_ of 5.5–42.7 in housed ducks over a week-long infectious period (Niqueux et al. 2013). For laughing gulls, which shed mainly from the oropharynx, a water-borne contact term is the weakest assumption (Criado et al. 2021).

#### S6.5 Cross-system settings

**Contact CV of 1 outside Lyme disease.** Only the Lyme value is measured. Fig. 2 expresses contact relative to its mean, so the CV sets how strongly a given correlation ρ can reweight individuals; a larger CV, as the nestling biting data suggest for West Nile virus, would be expected to strengthen the effects shown.

#### S6.6 Caveats

**ε–φ split for the chipmunk and masked shrew (Lyme; Fig. 3).** Their products ε × φ(*L*_max_) match published reservoir competence (Table S6), but Brunner et al. (2008, Table 1) attribute more to prevalence (0.971, 0.701) and less to infectivity (0.569, 0.537) than the coded ε (0.75) and φ(*L*_max_) (0.74). Infected larvae per host (Fig. 3A–B) depend on φ but not on ε, so a split following Brunner et al. would lower both species' Fig. 3 output by roughly 25–30% (our calculation); competence, potential and capacity (Fig. 4) depend on the product and would change little.

**Lyme infectivity does not wane.** In *P. leucopus* and robins, infectivity to larvae declines within the 60-day window in several studies (78% at day 13 to 45% at day 40 in mice; robins from 92% at days 28–30 to 68% at days 39–41; Lindsay et al. 1997; Hanincová et al. 2008; Richter et al. 2000), although one older study reports a decline lasting over 200 days (Donahue et al. 1987). The flat plateau makes *a* an upper bound for species that wane.

**American robin rise and competence (Lyme).** Robins infected 29% of larvae at days 6–9 and 85% at days 12–15 (Richter et al. 2000); the coded curve rises more slowly. ε × φ(*L*_max_) of 48% lies between the wild-caught (16.1%) and laboratory-infected (81.8%) estimates (Ginsberg et al. 2005).

**Opossum, skunk and deer below published reservoir competence (Lyme).** Model values are 0.4%, 1.7% and 0.1% against 2.6%, 9.7% and 4.6%. The xenodiagnostic assay has a specificity of about 0.97, so published values near 3% cannot be distinguished from false positives (Brunner et al. 2008); the skunk estimate rests on four animals.

**Lyme load units are relative.** Published loads come in non-convertible units (Barbour et al. 2009; Baum et al. 2012; Zinck et al. 2023); only *L*_max_/*K* matters.

**West Nile virus daily means are survivor-based.** Komar et al. (2003, Table 2) report log_10_ of arithmetic means, and later days include only surviving birds (our inference from the robin and blue jay entries). Applying φ to these means inflates infectiousness by roughly 0.4–1.0 log_10_ at the peak (our inference). Even after the refit, the simulated dove (mean *a* 0.33) ranks above the starling (0.25), the reverse of Komar's index (0.19 vs 0.22).

**Laughing gull shedding depends on the virus.** Gulls shed 10^4.5^–10^6.3^ EID_50_/mL of two gull-origin viruses but only 10^1.5^–10^1.7^ of two others (Criado et al. 2021), and shorebird- and mallard-origin viruses rarely infected them (Bahnson et al. 2020). The coded peak (3×10^3^) represents a poorly replicating strain.

**LPAI susceptibility is ≈1 for all three species.** The per-particle probabilities imply an ID_50_ of about 15–62 EID_50_, two to three log_10_ below measured LPAI infectious doses in wild birds (about 10^3.6^–10^4.2^; Hall et al. 2013), and experiments separate these species most sharply on infection success (Brown et al. 2012a; Costa et al. 2011b; Criado et al. 2021).

**Bullfrog load and infectiousness (Bd).** The code holds bullfrog load near 5×10^5^ ZE for years, but experimental bullfrogs carry low loads and often clear infection (Greenspan et al. 2012; Byrne et al. 2018); the Hill threshold would make realistic low loads nearly non-infectious, yet low-load bullfrogs transmit (Greenspan et al. 2012). The two assumptions err in opposite directions.

**Northern leopard frog mortality (Bd).** Adult NLF showed no Bd-related mortality in field and laboratory studies (Voordouw et al. 2010); froglets did die (Pask et al. 2013). The coded load-dependent mortality fits the froglet stage.

### References

AnAge: The Animal Ageing and Longevity Database. Entries for *Lithobates catesbeianus* and *Lithobates pipiens*. https://genomics.senescence.info (accessed October 2026).

Bahnson, C. S., Poulson, R. L., Hollander, L. P., Crum Bradley, J. A., & Stallknecht, D. E. (2020). Susceptibility of laughing gulls (*Leucophaeus atricilla*) and mallards (*Anas platyrhynchos*) to ruddy turnstone (*Arenaria interpres morinella*) origin type A influenza viruses. *Journal of Wildlife Diseases, 56*(1), 167–174. PMID 31532732.

Barbour, A. G., Bunikis, J., Fish, D., & Hanincová, K. (2015). Association between body size and reservoir competence of mammals bearing *Borrelia burgdorferi* at an endemic site in the northeastern United States. *Parasites & Vectors, 8*, 299. https://doi.org/10.1186/s13071-015-0903-5

Barbour, A. G., Bunikis, J., Travinsky, B., Hoen, A. G., Diuk-Wasser, M. A., Fish, D., & Tsao, J. I. (2009). Niche partitioning of *Borrelia burgdorferi* and *Borrelia miyamotoi* in the same tick vector and mammalian reservoir species. *The American Journal of Tropical Medicine and Hygiene, 81*(6), 1120–1131. https://doi.org/10.4269/ajtmh.2009.09-0208

Baum, E., Hue, F., & Barbour, A. G. (2012). Experimental infections of the reservoir species *Peromyscus leucopus* with diverse strains of *Borrelia burgdorferi*, a Lyme disease agent. *mBio, 3*(6), e00434-12. https://doi.org/10.1128/mBio.00434-12

Bourgeois, J. S., You, S. S., Clendenen, L. H., Shrestha, M., Petnicki-Ocwieja, T., Telford, S. R., & Hu, L. T. (2024). Comparative reservoir competence of *Peromyscus leucopus*, C57BL/6J, and C3H/HeN for *Borrelia burgdorferi* B31. *Applied and Environmental Microbiology, 90*(7), e00822-24. https://doi.org/10.1128/aem.00822-24

Breban, R., Drake, J. M., Stallknecht, D. E., & Rohani, P. (2009). The role of environmental transmission in recurrent avian influenza epidemics. *PLoS Computational Biology, 5*(4), e1000346. https://doi.org/10.1371/journal.pcbi.1000346

Brown, J. D., Berghaus, R. D., Costa, T. P., Poulson, R., Carter, D. L., Lebarbenchon, C., & Stallknecht, D. E. (2012a). Intestinal excretion of a wild bird-origin H3N8 low pathogenic avian influenza virus in mallards (*Anas platyrhynchos*). *Journal of Wildlife Diseases, 48*(4), 991–998. https://doi.org/10.7589/2011-09-280

Brown, J. D., Goekjian, G., Poulson, R., Valeika, S., & Stallknecht, D. E. (2009). Avian influenza virus in water: Infectivity is dependent on pH, salinity and temperature. *Veterinary Microbiology, 136*(1–2), 20–26. https://doi.org/10.1016/j.vetmic.2008.10.027

Brown, J. D., Poulson, R., Carter, D., Lebarbenchon, C., Pantin-Jackwood, M., Spackman, E., Shepherd, E., Killian, M., & Stallknecht, D. (2012b). Susceptibility of avian species to North American H13 low pathogenic avian influenza viruses. *Avian Diseases, 56*(4 Suppl.), 969–975. https://doi.org/10.1637/10158-040912-Reg.1

Brunner, J. L., LoGiudice, K., & Ostfeld, R. S. (2008). Estimating reservoir competence of *Borrelia burgdorferi* hosts: Prevalence and infectivity, sensitivity, and specificity. *Journal of Medical Entomology, 45*(1), 139–147. https://doi.org/10.1603/0022-2585(2008)45[139:ERCOBB]2.0.CO;2

Buck, J. C., Hua, J., Brogan, W. R., Dang, T. D., Urbina, J., Bendis, R. J., Stoler, A. B., Blaustein, A. R., & Relyea, R. A. (2015). Effects of pesticide mixtures on host-pathogen dynamics of the amphibian chytrid fungus. *PLoS ONE, 10*(7), e0132832. https://doi.org/10.1371/journal.pone.0132832

Byrne, A. Q., Poorten, T. J., Voyles, J., Willis, C. K. R., & Rosenblum, E. B. (2018). Opening the file drawer: Unexpected insights from a chytrid infection experiment. *PLoS ONE, 13*(5), e0196851. https://doi.org/10.1371/journal.pone.0196851

Caillouët, K. A., Robertson, C. W., Wheeler, D. C., Komar, N., & Bulluck, L. P. (2013). Vector contact rates on eastern bluebird nestlings do not indicate West Nile virus transmission in Henrico County, Virginia, USA. *International Journal of Environmental Research and Public Health, 10*(12), 6366. https://doi.org/10.3390/ijerph10126366

Chatfield, M. W. H., Brannelly, L. A., Robak, M. J., Freeborn, L., Lailvaux, S. P., & Richards-Zawacki, C. L. (2013). Fitness consequences of infection by *Batrachochytrium dendrobatidis* in northern leopard frogs (*Lithobates pipiens*). *EcoHealth, 10*(1), 90–98. https://doi.org/10.1007/s10393-013-0833-7

Costa, T. P., Brown, J. D., Howerth, E. W., & Stallknecht, D. E. (2010). Effect of a prior exposure to a low pathogenic avian influenza virus in the outcome of a heterosubtypic low pathogenic avian influenza infection in mallards (*Anas platyrhynchos*). *Avian Diseases, 54*(4), 1286–1291. https://doi.org/10.1637/9480-072210-Reg.1

Costa, T. P., Brown, J. D., Howerth, E. W., Stallknecht, D. E., & Swayne, D. E. (2011b). Homo- and heterosubtypic low pathogenic avian influenza exposure on H5N1 highly pathogenic avian influenza virus infection in wood ducks (*Aix sponsa*). *PLoS ONE, 6*(1), e15987. https://doi.org/10.1371/journal.pone.0015987

Criado, M. F., Moresco, K. A., Stallknecht, D. E., & Swayne, D. E. (2021). Low-pathogenicity influenza viruses replicate differently in laughing gulls and mallards. *Influenza and Other Respiratory Viruses, 15*(6), 701–706. https://doi.org/10.1111/irv.12878

Darbro, J. M., Dhondt, A. A., Vermeylen, F. M., & Harrington, L. C. (2007). *Mycoplasma gallisepticum* infection in house finches (*Carpodacus mexicanus*) affects mosquito blood feeding patterns. *The American Journal of Tropical Medicine and Hygiene, 77*(3), 488–494. PMID 17827365.

Devevey, G., & Brisson, D. (2012). The effect of spatial heterogeneity on the aggregation of ticks on white-footed mice. *Parasitology, 139*(7), 915–925. https://doi.org/10.1017/S003118201200008X

DiRenzo, G. V., Langhammer, P. F., Zamudio, K. R., & Lips, K. R. (2014). Fungal infection intensity and zoospore output of *Atelopus zeteki*, a potential acute chytrid supershedder. *PLoS ONE, 9*(3), e93356. https://doi.org/10.1371/journal.pone.0093356

Donahue, J. G., Piesman, J., & Spielman, A. (1987). Reservoir competence of white-footed mice for Lyme disease spirochetes. *The American Journal of Tropical Medicine and Hygiene, 36*(1), 92–96. https://doi.org/10.4269/ajtmh.1987.36.92

Dye, C., & Hasibeder, G. (1986). Population dynamics of mosquito-borne disease: Effects of flies which bite some people more frequently than others. *Transactions of the Royal Society of Tropical Medicine and Hygiene, 80*(1), 69–77. https://doi.org/10.1016/0035-9203(86)90199-9

Gervasi, S. S., Burgan, S. C., Hofmeister, E., Unnasch, T. R., & Martin, L. B. (2017). Stress hormones predict a host superspreader phenotype in the West Nile virus system. *Proceedings of the Royal Society B, 284*(1859), 20171090. PMID 28724737. Data: Dryad, https://doi.org/10.5061/dryad.1s9g1

Gervasi, S. S., Burkett-Cadena, N., Burgan, S. C., Schrey, A. W., Hassan, H. K., Unnasch, T. R., & Martin, L. B. (2016). Host stress hormones alter vector feeding preferences, success, and productivity. *Proceedings of the Royal Society B, 283*(1836), 20161278. https://doi.org/10.1098/rspb.2016.1278

Ginsberg, H. S., Buckley, P. A., Balmforth, M. G., Zhioua, E., Mitra, S., & Buckley, F. G. (2005). Reservoir competence of native North American birds for the Lyme disease spirochete, *Borrelia burgdorferi*. *Journal of Medical Entomology, 42*(3), 445–449. https://doi.org/10.1603/0022-2585(2005)042[0445:RCONNA]2.0.CO;2

Greenspan, S. E., Calhoun, A. J. K., Longcore, J. E., & Levy, M. G. (2012). Transmission of *Batrachochytrium dendrobatidis* to wood frogs (*Lithobates sylvaticus*) via a bullfrog (*L. catesbeianus*) vector. *Journal of Wildlife Diseases, 48*(3), 575–582. https://doi.org/10.7589/0090-3558-48.3.575

Hall, J. S., Krauss, S., Franson, J. C., TeSlaa, J. L., Nashold, S. W., Stallknecht, D. E., Webby, R. J., & Webster, R. G. (2013). Avian influenza in shorebirds: Experimental infection of ruddy turnstones (*Arenaria interpres*) with avian influenza virus. *Influenza and Other Respiratory Viruses, 7*(1), 85–92. https://doi.org/10.1111/j.1750-2659.2012.00358.x

Hanincová, K., Ogden, N. H., Diuk-Wasser, M., Pappas, C. J., Iyer, R., Fish, D., Schwartz, I., & Kurtenbach, K. (2008). Fitness variation of *Borrelia burgdorferi* sensu stricto strains in mice. *Applied and Environmental Microbiology, 74*(1), 153–157. https://doi.org/10.1128/AEM.01567-07

Huang, C.-I., Kay, S. C., Davis, S., Tufts, D. M., Gaffett, K., Tefft, B., & Diuk-Wasser, M. A. (2019). High burdens of *Ixodes scapularis* larval ticks on white-tailed deer may limit Lyme disease risk in a low biodiversity setting. *Ticks and Tick-borne Diseases, 10*(2), 258–268. https://doi.org/10.1016/j.ttbdis.2018.10.013

Jani, A. J., & Briggs, C. J. (2018). Host and aquatic environment shape the amphibian skin microbiome but effects on downstream resistance to the pathogen *Batrachochytrium dendrobatidis* are variable. *Frontiers in Microbiology, 9*, 487. https://doi.org/10.3389/fmicb.2018.00487

Jani, A. J., Bushell, J., Arisdakessian, C. G., Belcaid, M., Boiano, D. M., Brown, C., & Knapp, R. A. (2021). The amphibian microbiome exhibits poor resilience following pathogen-induced disturbance. *The ISME Journal, 15*(6), 1628–1640. https://doi.org/10.1038/s41396-020-00875-w

Jourdain, E., Gunnarsson, G., Wahlgren, J., Latorre-Margalef, N., Bröjer, C., Sahlin, S., Svensson, L., Waldenström, J., Lundkvist, Å., & Olsen, B. (2010). Influenza virus in a natural host, the mallard: Experimental infection data. *PLoS ONE, 5*(1), e8935. https://doi.org/10.1371/journal.pone.0008935

Keesing, F., Brunner, J., Duerr, S., Killilea, M., LoGiudice, K., Schmidt, K., Vuong, H., & Ostfeld, R. S. (2009). Hosts as ecological traps for the vector of Lyme disease. *Proceedings of the Royal Society B, 276*(1675), 3911–3919. https://doi.org/10.1098/rspb.2009.1159

Knapp, R. A., Joseph, M. B., Smith, T. C., Hegeman, E. E., Vredenburg, V. T., Erdman, J. E., Boiano, D. M., Jani, A. J., & Briggs, C. J. (2022). Effectiveness of antifungal treatments during chytridiomycosis epizootics in populations of an endangered frog. *PeerJ, 10*, e12712. https://doi.org/10.7717/peerj.12712

Komar, N., Langevin, S., Hinten, S., Nemeth, N., Edwards, E., Hettler, D., Davis, B., Bowen, R., & Bunning, M. (2003). Experimental infection of North American birds with the New York 1999 strain of West Nile virus. *Emerging Infectious Diseases, 9*(3), 311–322. https://doi.org/10.3201/eid0903.020628

Latorre-Margalef, N., Gunnarsson, G., Munster, V. J., Fouchier, R. A. M., Osterhaus, A. D. M. E., Elmberg, J., Olsen, B., Wallensten, A., Haemig, P. D., Fransson, T., Brudin, L., & Waldenström, J. (2009). Effects of influenza A virus infection on migrating mallard ducks. *Proceedings of the Royal Society B, 276*(1659), 1029–1036. https://doi.org/10.1098/rspb.2008.1501

Levi, T., Keesing, F., Holt, R. D., Barfield, M., & Ostfeld, R. S. (2016). Quantifying dilution and amplification in a community of hosts for tick-borne pathogens. *Ecological Applications, 26*(2), 484–498. https://doi.org/10.1890/15-0122

Lindsay, L. R., Barker, I. K., Surgeoner, G. A., McEwen, S. A., & Campbell, G. D. (1997). Duration of *Borrelia burgdorferi* infectivity in white-footed mice for the tick vector *Ixodes scapularis* under laboratory and field conditions in Ontario. *Journal of Wildlife Diseases, 33*(4), 766–775. https://doi.org/10.7589/0090-3558-33.4.766

Lindsay, L. R., Barker, I. K., Surgeoner, G. A., McEwen, S. A., Gillespie, T. J., & Addison, E. M. (1998). Survival and development of the different life stages of *Ixodes scapularis* (Acari: Ixodidae) held within four habitats on Long Point, Ontario, Canada. *Journal of Medical Entomology, 35*(3), 189–199. https://doi.org/10.1093/jmedent/35.3.189

LoGiudice, K., Ostfeld, R. S., Schmidt, K. A., & Keesing, F. (2003). The ecology of infectious disease: Effects of host diversity and community composition on Lyme disease risk. *Proceedings of the National Academy of Sciences, 100*(2), 567–571. https://doi.org/10.1073/pnas.0233733100

Mather, T. N., Wilson, M. L., Moore, S. I., Ribeiro, J. M., & Spielman, A. (1989). Comparing the relative potential of rodents as reservoirs of the Lyme disease spirochete (*Borrelia burgdorferi*). *American Journal of Epidemiology, 130*(1), 143–150. https://doi.org/10.1093/oxfordjournals.aje.a115306

Niqueux, É., Picault, J.-P., Amelot, M., Allée, C., Lamandé, J., Guillemoto, C., Pierre, I., Massin, P., Blot, G., Briand, F.-X., Rose, N., & Jestin, V. (2013). Quantitative transmission characteristics of different H5 low pathogenic avian influenza viruses in Muscovy ducks. *Veterinary Microbiology, 168*(1), 78–87. https://doi.org/10.1016/j.vetmic.2013.10.020

Ostfeld, R. S., Canham, C. D., Oggenfuss, K., Winchcombe, R. J., & Keesing, F. (2006). Climate, deer, rodents, and acorns as determinants of variation in Lyme-disease risk. *PLoS Biology, 4*(6), e145. https://doi.org/10.1371/journal.pbio.0040145

Otis, D. L., & Dukes, W. F., Jr. (1995). Improving wood duck survival estimates in the mid-Atlantic region by using nest box banding programs. *Proceedings of the Annual Conference of the Southeastern Association of Fish and Wildlife Agencies, 49*, 618–628.

Pask, J. D., Cary, T. L., & Rollins-Smith, L. A. (2013). Skin peptides protect juvenile leopard frogs (*Rana pipiens*) against chytridiomycosis. *Journal of Experimental Biology, 216*(15), 2908–2916. https://doi.org/10.1242/jeb.084145

Peterson, A. C., & McKenzie, V. J. (2014). Investigating differences across host species and scales to explain the distribution of the amphibian pathogen *Batrachochytrium dendrobatidis*. *PLoS ONE, 9*(9), e107441. https://doi.org/10.1371/journal.pone.0107441

Reeder, N. M. M., Pessier, A. P., & Vredenburg, V. T. (2012). A reservoir species for the emerging amphibian pathogen *Batrachochytrium dendrobatidis* thrives in a landscape decimated by disease. *PLoS ONE, 7*(3), e33567. https://doi.org/10.1371/journal.pone.0033567

Richter, D., Spielman, A., Komar, N., & Matuschka, F.-R. (2000). Competence of American robins as reservoir hosts for Lyme disease spirochetes. *Emerging Infectious Diseases, 6*(2), 133–138. https://doi.org/10.3201/eid0602.000205

Roberts, A. J., Hostetler, J. A., Stiller, J. C., Devers, P. K., & Link, W. A. (2023). Population dynamics and harvest management of eastern mallards. *Journal of Wildlife Management*, e22405. https://doi.org/10.1002/jwmg.22405

Saltelli, A., Annoni, P., Azzini, I., Campolongo, F., Ratto, M., & Tarantola, S. (2010). Variance based sensitivity analysis of model output. Design and estimator for the total sensitivity index. *Computer Physics Communications, 181*, 259–270. https://doi.org/10.1016/j.cpc.2009.09.018

Sardelis, M. R., Turell, M. J., Dohm, D. J., & O'Guinn, M. L. (2001). Vector competence of selected North American *Culex* and *Coquillettidia* mosquitoes for West Nile virus. *Emerging Infectious Diseases, 7*(6), 1018–1022. https://doi.org/10.3201/eid0706.010617

Sauer, E. L., Fuller, R. C., Richards-Zawacki, C. L., Sonn, J., Sperry, J. H., & Rohr, J. R. (2018). Variation in individual temperature preferences, not behavioural fever, affects susceptibility to chytridiomycosis in amphibians. *Proceedings of the Royal Society B, 285*(1885). https://doi.org/10.1098/rspb.2018.1111

Stallknecht, D. E., Kearney, M. T., Shane, S. M., & Zwank, P. J. (1990). Effects of pH, temperature, and salinity on persistence of avian influenza viruses in water. *Avian Diseases, 34*(2), 412–418. PMID 2142421.

Tufts, D. M., McClure, M., & Diuk-Wasser, M. A. (2021). *Ixodes scapularis* (Acari: Ixodidae) nymphal survival and host-finding success in the eastern United States. *Journal of Medical Entomology, 58*(2), 929–938. https://doi.org/10.1093/jme/tjaa246

Voordouw, M. J., Adama, D., Houston, B., Govindarajulu, P., & Robinson, J. (2010). Prevalence of the pathogenic chytrid fungus, *Batrachochytrium dendrobatidis*, in an endangered population of northern leopard frogs, *Rana pipiens*. *BMC Ecology, 10*, 6. https://doi.org/10.1186/1472-6785-10-6

Voyles, J., Johnson, L. R., Briggs, C. J., Cashins, S. D., Alford, R. A., Berger, L., Skerratt, L. F., Speare, R., & Rosenblum, E. B. (2012a). Temperature alters reproductive life history patterns in *Batrachochytrium dendrobatidis*, a lethal pathogen associated with the global loss of amphibians. *Ecology and Evolution, 2*(9), 2241–2249. https://doi.org/10.1002/ece3.334

Voyles, J., Vredenburg, V. T., Tunstall, T. S., Parker, J. M., Briggs, C. J., & Rosenblum, E. B. (2012b). Pathophysiology in mountain yellow-legged frogs (*Rana muscosa*) during a chytridiomycosis outbreak. *PLoS ONE, 7*(4), e35374. https://doi.org/10.1371/journal.pone.0035374

Vredenburg, V. T., Knapp, R. A., Tunstall, T. S., & Briggs, C. J. (2010). Dynamics of an emerging disease drive large-scale amphibian population extinctions. *Proceedings of the National Academy of Sciences, 107*(21), 9689–9694. https://doi.org/10.1073/pnas.0914111107

Wilber, M. Q., Ohmer, M. E. B., Altman, K. A., Brannelly, L. A., LaBumbard, B. C., Le Sage, E. H., McDonnell, N. B., Muñiz Torres, A. Y., Nordheim, C. L., Pfab, F., Richards-Zawacki, C. L., Rollins-Smith, L. A., Saenz, V., Voyles, J., Wetzel, D. P., Woodhams, D. C., & Briggs, C. J. (2022). Once a reservoir, always a reservoir? Seasonality affects the pathogen maintenance potential of amphibian hosts. *Ecology, 103*(9), e3759. https://doi.org/10.1002/ecy.3759

Zinck, C. B., Raveendram Thampy, P., Uhlemann, E.-M. E., Adam, H., Wachter, J., Suchan, D., Cameron, A. D. S., Rego, R. O. M., Brisson, D., Bouchard, C., Ogden, N. H., & Voordouw, M. J. (2023). Variation among strains of *Borrelia burgdorferi* in host tissue abundance and lifetime transmission determine the population strain structure in nature. *PLoS Pathogens, 19*(8), e1011572. https://doi.org/10.1371/journal.ppat.1011572
