## Appendix S3 for "Formalizing host competence"

**Appendix S3. Sources of individual variation in infectiousness: a variance decomposition**

*Supporting Information for Gibson et al., Formalizing host competence*

**S3.1 Purpose and scope**

This appendix reports the variance decomposition that the main text summarizes at the start of Section 4. To determine which host traits drive individual variation in competence, we decomposed variance in infectiousness, *a*, with Sobol analyses (Fig. S1) though we did not include susceptibility. Susceptibility matters at the population level because heterogeneity in susceptibility determines who becomes infected (Dwyer et al. 1997; Langwig et al. 2017). However, too few data from dose-dependent experimental exposures were available for most species. So we could not investigate it meaningfully here.

**S3.2 Design**

For a high- and a low-competence species in each of the four systems (for chytridiomycosis, the tolerant bullfrog and the mountain yellow-legged frog, in which infection is lethal), we estimated first-order and total-order Sobol indices of infectiousness, *a* (main text Eq. 5), with respect to every parameter that varies among individuals. Parameters were drawn from the same among-individual distributions used to simulate individuals (Appendix S2), not from uniform ranges, so the indices partition the among-individual variance that was actually simulated. We drew two independent base samples of N = 16,384 parameter sets (4,096 for the bullfrog, whose integration window spans 9,000 days) and, for each parameter, a third matrix in which that parameter’s column in the first sample was replaced by its column in the second. First-order indices used the Saltelli estimator and total-order indices the Jansen estimator (Saltelli et al. 2010). Model outputs were centered on the mean of the two base samples before the first-order estimator was applied; this leaves its expected value unchanged and reduces seed noise. Intervals are the 2.5th and 97.5th percentiles of 200 bootstrap resamples. Indices are unitless proportions of the among-individual variance in *a* (0–1); small negative values, and first-order indices slightly above total-order ones, reflect Monte Carlo error. Susceptibility ε and contact θ do not enter *a* and are excluded, as are the Hill parameters K and n, which are fixed across individuals. For Lyme disease, the behavior term is the feeding survival of attached larvae, which occupies the b slot in the code (Appendix S2, Table S2a). The analysis is figs.py::sobol_within in the Python pipeline, which produced Fig. S1 and Table S7; the R pipeline (02_host_competence_figures.Rmd) gives the same indices within Monte Carlo error.

**S3.3 Results**

The parameter that explained most among-individual variation in infectiousness (i.e., the first-order Sobol index, or the proportion of the variance in *a* among individuals of a species that was attributable to one parameter alone (unitless, 0–1; Saltelli et al. 2010)) differed among disease systems. Load magnitude dominated in avian influenza (mallard 0.54, laughing gull 0.84), West Nile virus (blue jay 0.52, mourning dove 0.95), and Lyme disease (white-footed mouse 0.76, gray squirrel 0.92) with load timing (mallard 0.35) and infection duration (blue jay 0.27) secondary in the high-competence species. In chytrid, by contrast, load mattered little because loads far exceeded the threshold for infectiousness: natural lifespan dominated in the bullfrog (0.76), and behavior (0.67), load growth rate (0.18) and disease-induced mortality (0.10) in the mountain yellow-legged frog. In every system but chytrid, load magnitude explained more of the variation in the low-competence species than in the high-competence one, because low-competence species’ loads sit near the threshold at which small differences in load produced large differences in infectiousness.

Considered together, consequential variation in host infectiousness arose through one of three paths: differences in pathogen load where transmission requires load to exceed a threshold, differences in how long peak load lasts where transmission is confined to a brief window, or differences in survival where infections persist such that some individuals transmit for far longer than others. These alternatives emphasize that hosts could be similarly competent in different ways. For instance, in chytridiomycosis, a highly susceptible but vulnerable host would be practically incompetent because disease-induced mortality would truncate the transmission window. By contrast, a highly-resistant host could be very competent if it was also tolerant (e.g., low per pathogen effects on host health) of what pathogen burden it maintained.

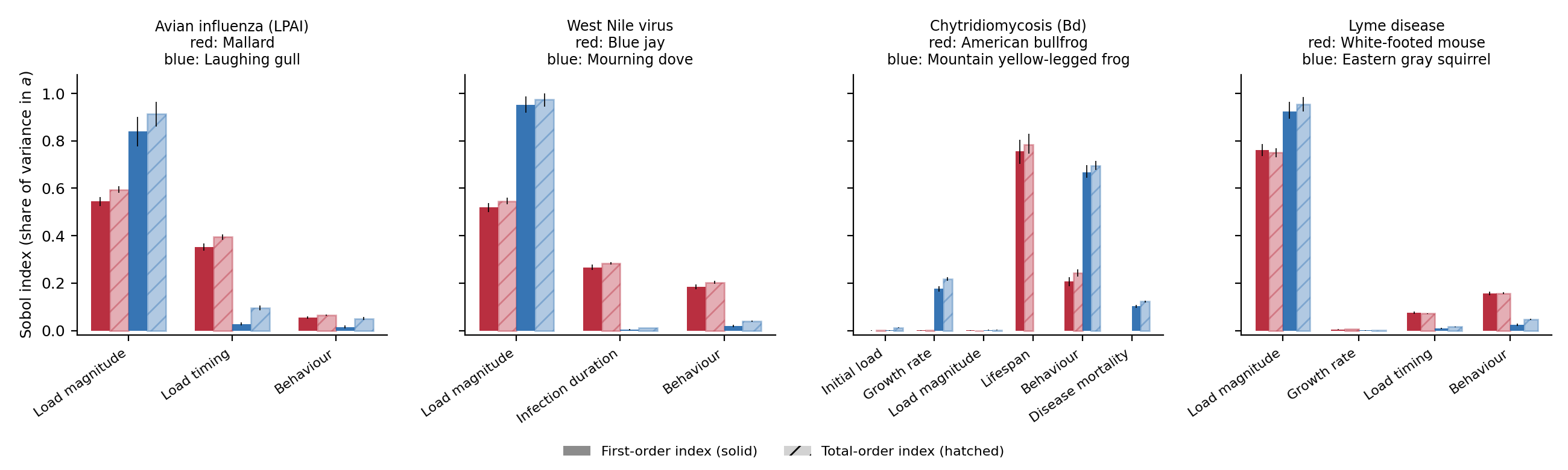

**Figure S1.** Sources of among-individual variation in infectiousness, *a*, for a high-competence (red) and a low-competence (blue) species in each system. Parameters were drawn from the same distributions used to simulate individuals rather than from uniform ranges, and first-order (solid) and total-order (hatched) indices were estimated with the Saltelli and Jansen estimators (Saltelli et al. 2010; N = 16,384 base samples, 4,096 for the bullfrog), with 95% bootstrap intervals. Load magnitude is peak or plateau load (for West Nile virus, an individual’s offset in log_10_ viremia). Lifespan is natural lifespan (bullfrog); disease mortality is the load-dependent mortality coefficient (mountain yellow-legged frog). The Hill parameters K and n are fixed in the simulations and are therefore excluded; their influence concerns model structure rather than individual variation (Appendix S1, Section 6).

**Table S7.** First- and total-order Sobol indices of infectiousness, *a*, with 95% bootstrap intervals (200 resamples), for the analysis in Fig. S1. Values are rounded to two decimals; “0.00” marks indices below 0.005 in absolute value.

| **System** | **Species (competence)** | **Parameter** | **First-order index (95% interval)** | **Total-order index (95% interval)** |
| --- | --- | --- | --- | --- |
| LPAI | Mallard (high) | Load magnitude | 0.54 (0.52–0.56) | 0.59 (0.58–0.61) |
|  |  | Load timing | 0.35 (0.34–0.37) | 0.39 (0.38–0.40) |
|  |  | Behavior | 0.05 (0.05–0.06) | 0.07 (0.06–0.07) |
|  | Laughing gull (low) | Load magnitude | 0.84 (0.78–0.90) | 0.91 (0.86–0.96) |
|  |  | Load timing | 0.03 (0.02–0.03) | 0.10 (0.09–0.11) |
|  |  | Behavior | 0.01 (0.01–0.02) | 0.05 (0.04–0.06) |
| West Nile virus | Blue jay (high) | Load magnitude | 0.52 (0.50–0.54) | 0.55 (0.53–0.56) |
|  |  | Infection duration | 0.27 (0.26–0.28) | 0.28 (0.28–0.29) |
|  |  | Behavior | 0.18 (0.17–0.19) | 0.20 (0.20–0.21) |
|  | Mourning dove (low) | Load magnitude | 0.95 (0.92–0.99) | 0.97 (0.94–1.00) |
|  |  | Infection duration | 0.00 (0.00–0.01) | 0.01 (0.01–0.01) |
|  |  | Behavior | 0.02 (0.02–0.02) | 0.04 (0.04–0.04) |
| Chytridiomycosis | American bullfrog (tolerant) | Initial load | 0.00 (0.00–0.00) | 0.00 (0.00–0.00) |
|  |  | Growth rate | 0.00 (0.00–0.00) | 0.00 (0.00–0.00) |
|  |  | Load magnitude | 0.00 (0.00–0.00) | 0.00 (0.00–0.00) |
|  |  | Lifespan | 0.76 (0.70–0.80) | 0.78 (0.75–0.83) |
|  |  | Behavior | 0.21 (0.19–0.23) | 0.24 (0.23–0.26) |
|  | Mountain yellow-legged frog (lethal) | Initial load | 0.00 (0.00–0.00) | 0.01 (0.01–0.01) |
|  |  | Growth rate | 0.18 (0.16–0.19) | 0.22 (0.21–0.23) |
|  |  | Load magnitude | 0.00 (0.00–0.00) | 0.00 (0.00–0.00) |
|  |  | Disease mortality | 0.10 (0.10–0.11) | 0.12 (0.12–0.13) |
|  |  | Behavior | 0.67 (0.65–0.70) | 0.70 (0.68–0.71) |
| Lyme disease | White-footed mouse (high) | Load magnitude | 0.76 (0.74–0.79) | 0.75 (0.73–0.77) |
|  |  | Growth rate | 0.00 (0.00–0.01) | 0.01 (0.01–0.01) |
|  |  | Load timing | 0.07 (0.07–0.08) | 0.07 (0.07–0.07) |
|  |  | Behavior (feeding survival) | 0.16 (0.15–0.16) | 0.16 (0.15–0.16) |
|  | Eastern gray squirrel (low) | Load magnitude | 0.92 (0.89–0.96) | 0.95 (0.92–0.98) |
|  |  | Growth rate | 0.00 (0.00–0.00) | 0.00 (0.00–0.00) |
|  |  | Load timing | 0.01 (0.01–0.01) | 0.02 (0.01–0.02) |
|  |  | Behavior (feeding survival) | 0.02 (0.02–0.03) | 0.05 (0.04–0.05) |

**References**

Dwyer, G., Elkinton, J. S., & Buonaccorsi, J. P. (1997). Host heterogeneity in susceptibility and disease dynamics: tests of a mathematical model. *American Naturalist*, 150, 685–707.

Langwig, K. E., Wargo, A. R., Jones, D. R., Viss, J. R., Rutan, B. J., Egan, N. A., et al. (2017). Vaccine effects on heterogeneity in susceptibility and implications for population health management. *mBio*, 8, e00796-17.

Saltelli, A., Annoni, P., Azzini, I., Campolongo, F., Ratto, M., & Tarantola, S. (2010). Variance based sensitivity analysis of model output. Design and estimator for the total sensitivity index. *Computer Physics Communications*, 181, 259–270.
